# A segmental duplication-mediated deletion leads to neocentromere formation in orangutans

**DOI:** 10.64898/2026.04.09.717302

**Authors:** Luciana de Gennaro, DongAhn Yoo, Letizia Pistacchia, Rosaria Magrone, Alessia Daponte, Francesco Perrone, Francesco Ravasini, F. Kumara Mastrorosa, Keisuke K. Oshima, Cesare Polano, Kendra Hoekzema, Katherine M. Munson, Julie Wertz, Fabio Marroni, Claudia R. Catacchio, Francesca Antonacci, Daan Noordermeer, Francesco Montinaro, Glennis A. Logsdon, Beniamino Trombetta, Evan E. Eichler, Mario Ventura

## Abstract

Centromeres ensure faithful chromosome segregation, yet how new centromeres arise and replace canonical ones remains poorly understood. Here, we investigate a polymorphic centromere repositioning event on the orangutan chromosome 10 using near-telomere-to-telomere assemblies, epigenetic profiling, and population-scale data. We identify striking heterogeneity in canonical centromeres, ranging from large, higher-order repeat α-satellite arrays to short, monomeric α-satellite tracts, alongside the emergence of neocentromeres lacking α-satellite DNA. We show a segmental duplication-mediated deletion of 3.6 Mbp that removed the higher-order repeat array, promoting centromere repositioning and neocentromere formation. Phylogenetic analyses reveal complex evolutionary dynamics, including introgression and incomplete lineage sorting in orangutan lineages. These findings demonstrate that centromere identity can evolve through structural variation and epigenetic reprogramming, highlighting its remarkable plasticity in primate genomes.

---

The centromere (CEN), the primary constriction of the chromosome visible during metaphase, is essential for proper chromosome segregation due to its role as the assembly site for the kinetochore complex (*1, 2*). Its activity may occasionally relocate to a *de novo* ectopic location, lacking α-satellite (α-sat) DNA, and generate neocentromeres(*3–8*). Several investigations have reported neocentromeres in both pathological rearrangements and in individuals with a normal karyotype, in which the centromeric function shifts to a nearby euchromatic site, while the canonical α-sat region remains physically intact but inactive(*9–11*). Neocentromere formation may also be associated with deletions at the original kinetochore site, resulting in a substantial reduction of α-sat DNA and contributing to centromere inactivation and neocentromere emergence(*12*).

Furthermore, neocentromere positioning is not entirely random: several recurrent loci in human, such as the region at 15q24-26, overlap with ancestral centromeres that were inactivated during primate evolution, suggesting the existence of genomic contexts permissive to centromere reactivation(*13*). However, recent high-resolution analyses indicate that neocentromeric regions do not share obvious sequence-based hallmarks, such as specific gene content or repeat composition, but are instead characterized by reduced heterozygosity across haplotypes, suggesting population-level or chromatin-associated constraints rather than a universal DNA sequence signature(*12*).

The formation of evolutionary new centromeres (ENCs) represents the evolutionary counterpart to neocentromeres: a repositioned centromeric activity that has become fixed and inherited within a species lineage(*7*). ENCs initially appear in sequence-agnostic regions as neocentromeres before gradually acquiring the canonical hallmarks of a mature centromere, such as *de novo* α-sat arrays(*14, 15*). ENCs are recognized as a significant force in genomic evolution, acting independently of classical chromosomal rearrangements(*13, 14, 16, 17*).

Despite the high frequency in primate genome evolution, a detailed understanding of the molecular mechanisms driving the *de novo* formation and evolutionary fixation of centromeres among primates remains unclear. In particular, while recent studies have begun to hypothesize the mechanisms underlying *de novo* formation of human neocentromeres, the complete processes required for the formation of human neocentromeres and ENCs are still largely unknown(*12, 18*), as the limited availability of fully assembled allelic sequences and comprehensive epigenetic profiles constrains our ability to precisely track both the new centromere formation and the simultaneous behavior of the canonical centromere.

Sequencing and cytogenetic studies have reported putative ENCs in both Sumatran (*Pongo abelii*; PAB) and Bornean (*Pongo pygmaeus*, PPY) orangutan species, with an ENC at chromosome 10 (chr10_hsa12) occurring at ∼26% allele frequency(*11, 19, 20*). Preliminary studies of this region have identified the active centromeric domain as spanning roughly 605 kbp. Even though these species have been sequenced before, it is only with the recent emergence of near-complete, high-quality centromeric assemblies(*21*) that we are now able to investigate their complete sequence composition and structural architecture.

Here, we investigate the molecular and epigenetic landscape of the genomic region where the centromere activity repositioning occurred, by generating three new high-quality orangutan genome assemblies, one PAB and two PPY, each exhibiting a distinct centromere status on chromosome 10. Leveraging deep long-read sequencing (PacBio high-fidelity [HiFi] and Oxford Nanopore Technologies [ONT]) combined with Micro-C for haplotype phasing, we assembled all chromosome 10 haplotypes to near-telomere-to-telomere (T2T) resolution, with each represented by a single contig spanning almost the entire chromosome. This level of completeness enables the characterization of sequence organization, DNA methylation, and transcriptional activity across both canonical and neocentromeric regions using complementary functional assays, including long-read isoform sequencing (Iso-Seq). In addition, by integrating Illumina whole-genome sequencing data from 52 orangutans (25 PAB and 27 PPY), including both publicly available and newly generated datasets, we extend our analyses to a population scale. Together, these data provide a framework for investigating the structural and epigenetic features associated with centromere repositioning and for exploring the evolutionary processes underlying the emergence and maintenance of neocentromeres in orangutans.

## Centromere and neocentromere diversity across Sumatran and Bornean orangutans

Cytogenetic and sequence analyses were performed on a combined population of 17 orangutans representing both the Sumatran (PAB) and Bornean (PPY) species, which diverged approximately 0.96 million years ago (Mya)(*21*) (Fig. 1A; 1B). We found that the putative ENCs in orangutans were missing α-sat sequences, thus representing neocentromeres (NEOs) rather than ENCs. In this regard, cytogenetic profiling revealed the presence of sex chromosome aneuploidy and mosaicism in some cell lines and identified three centromeric states on chr10_hsa12 (canonical CEN/CEN, heterozygous CEN/NEO, and homozygous NEO/NEO), with distinct distributions across species (Supplementary Text 1; fig. S1; table S1). In our sample set, neocentromeres showed a higher prevalence in Bornean individuals, occurring in 33.3% of haplotypes (4 out of 14 haplotypes from 7 individuals), compared to 25% in Sumatran orangutans (5 out of 20 haplotypes from 10 individuals), where canonical configurations predominated (table S1). Population structure analyses, using 35 orangutan genomes from publicly available data and encompassing both chromosome 10 and the neocentromeric region (Supplementary Text 2), demonstrated a separation of the two species and identified a Borneo × Sumatran hybrid between the two clusters (Fig. 1C). A principal component analysis (PCA) of the neocentromeric region revealed specific stratification within the Sumatran lineage that correlates with neocentromere genotype (Fig. 1D). Based on this data, three individuals representative of the different species and neocentromere configurations (PAB16 CEN/NEO, PPY17 CEN/NEO, and PPY15 NEO/NEO) (Fig. 1E) were selected for long-read sequencing, enabling a high-resolution study of their structure and evolution (Supplementary Text 1-2).

**Fig. 1.**
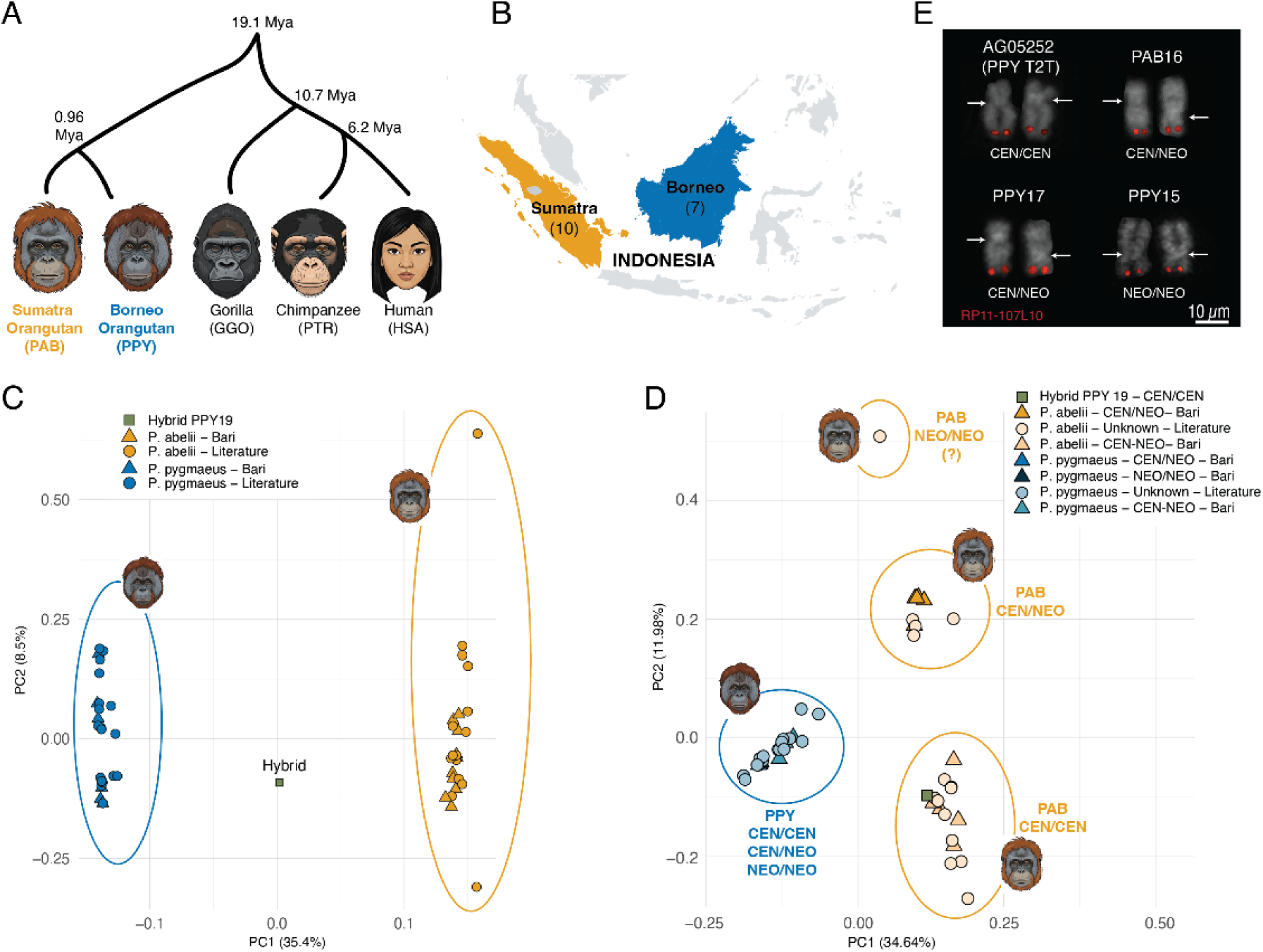
Orangutan cohort phylogeny, geographic distribution, cytogenetic characterization, and PCAs. **(A)** Phylogenetic tree showing evolutionary relationships and divergence times among the Sumatran orangutan (*Pongo abelii*, PAB), Bornean orangutan (*Pongo pygmaeus*, PPY), gorilla (*Gorilla gorilla*, GGO), chimpanzee (*Pan troglodytes*, PTR), and human (*Homo sapiens*, HSA). Node values indicate estimated divergence times in million years ago (Mya) based on Yoo et al., 2025(*21*). In particular Sumatra-Borneo 0.96 (0.92-1.00), human-chimpanzee 6.2 (5.6-6.3) Mya, gorilla-human-chimpanzee 10.7 (10.6-10.9), all great ape 19.1 (18.2-19.6). **(B)** Geographic distribution in Southeast Asia-Indonesia of our orangutan samples (the number of samples is reported in brackets). **(C)** Principal component analysis (PCA) carried out for the complete orangutan sequence for chromosome 10 and **(D)** on the neocentromeric region, including 250 kbp of its flanking regions (homologous to NC_071995.2:82,842,621–83,474,431 in mPonAbe1-v2.1_pri). In (D), cluster labels indicate centromeric status. In PAB, the co-clustering of our 10 cytogenetically characterized samples with the 15 publicly available sequenced genomes enabled the assignment of centromeric status to the latter. The symbol (?) denotes a tentative assignment, as it refers to a separate sample whose status cannot be validated because no sample in our cohort displays the corresponding phenotype (NEO/NEO); accordingly, its cluster assignment remains uncertain. In PPY, no distinct clustering was observed. **(E)** Cytogenetic characterization of neocentromere status using fluorescence *in situ* hybridization (FISH) for AG05252 (PPY T2T), PAB16, PPY17, PPY15. The last three cell lines were selected for long-read sequencing as representatives of the observed neocentromeric diversity. Across the full cohort (n = 17), the distribution of centromere (CEN) and neocentromere (NEO) alleles was as follows: among Sumatra samples (n = 10), 5 were CEN/CEN, and 5 were CEN/NEO; among Borneo samples (n = 7), 4 were CEN/CEN, 3 were CEN/NEO, and 1 was NEO/NEO (table S1).

## α-sat organization and centromere activity across orangutan chromosome 10 haplotypes

Combining PacBio HiFi reads, ultra-long ONT reads, and Illumina short reads, together with Micro-C, we generated high-quality, near-T2T haplotype-resolved assemblies for three representative orangutans using Verkko (PAB16, PPY17, PPY15) (v2.2.1;(*22*)). The high contiguity (haploid N50 ∼138–146 Mbp), baseline accuracy (quality value [QV] ∼56–59), and gene completeness >98% of the assemblies allowed reliable haplotype reconstruction of chromosome 10 (Supplementary Text 3). The assembled contigs from chromosome 10 generated in this study have been made publicly accessible (Data, code, and materials availability). Comparative analysis revealed structural and epigenetic heterogeneity at centromeric and neocentromeric loci (Supplementary Text 4). We observed differences in (i) α-sat array length (from ∼118 kbp to >2.1 Mbp), (ii) genomic organization of the repetitive regions (monomeric α-sat versus higher-order repeat (HOR) structures), (iii) degree of LINE1 interspersion (3–18 LINE1 elements >5 kbp embedded within α-sat arrays across haplotypes), (iv) sequence identity (<85% in monomeric/neocentromeric regions versus >95% in α-sat HOR arrays), and (v) functional activation marked by the presence of centromere dip regions (CDRs) and CENP-A/CENP-C enrichment (Fig. 2 and fig. S2).

**Fig. 2.**
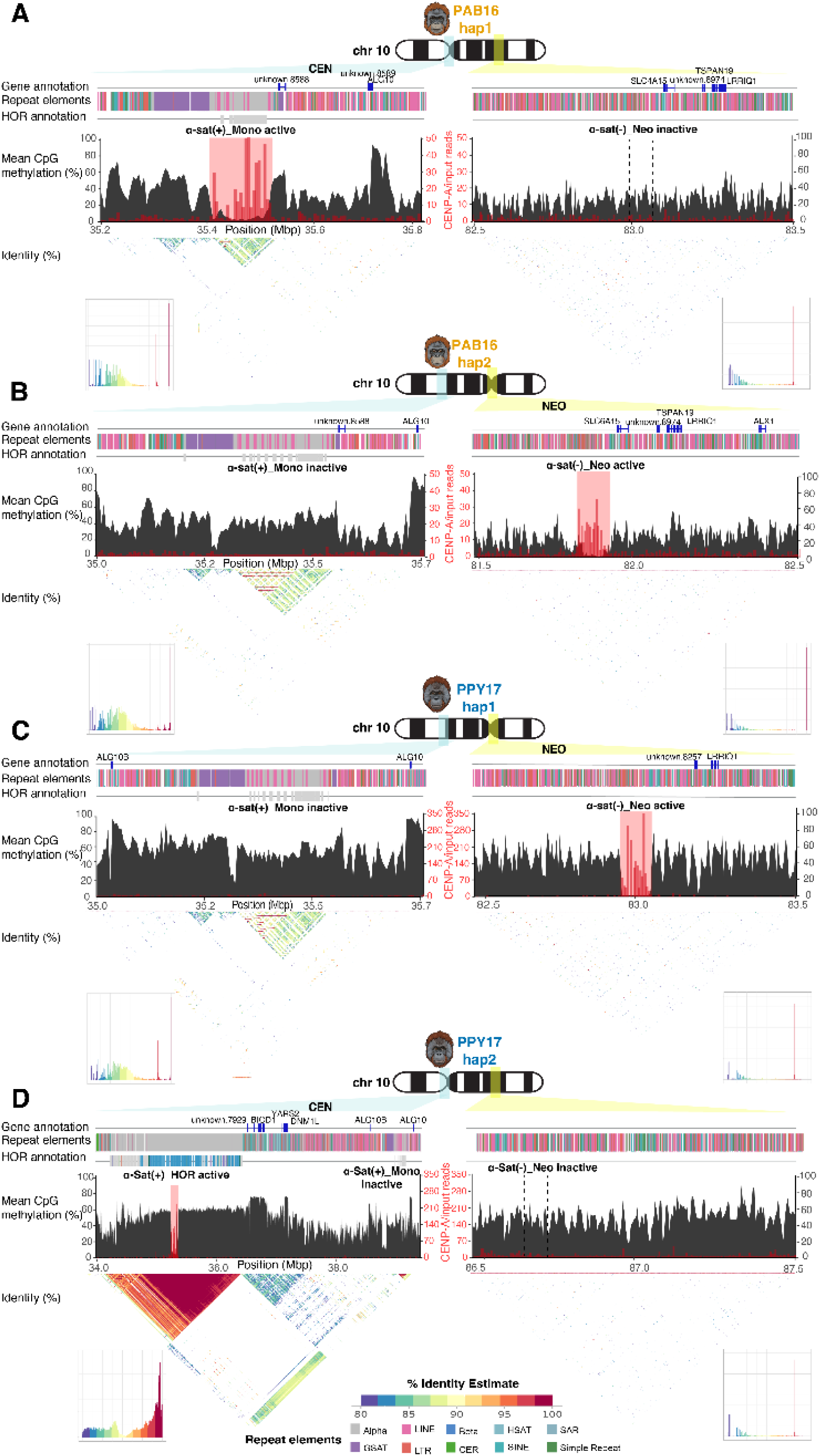
Architecture of the centromere and neocentromere regions of chromosome 10. Comparative representation of the organization of the centromere of chromosome 10 (chr10) in samples PAB16 and PPY17, which are both heterozygous for the neocentromere. For each haplotype (hap1 & hap2), we depict the centromeric (left; light blue in the ideogram) and neocentromeric (right; light yellow in the ideogram) regions. The following are shown from top to bottom: (i) gene annotation; (ii) composition of repeat element classes (including alpha-satellite [α-sat], HSAT, LTR, LINE, 1NE, and simple repeats); (iii) organization of α-sat units, distinguishing between α-sat(+)_Mono, α-sat(+)_HOR, and α-sat(−)_Neo, further classified as active or inactive based on methylation profiles; (iv) methylation profile (dark grey) and ratio (red) of CENP-A chip reads/input reads (5 kbp bins), and (v) sequence identity percentage. Each panel represents a different haplotype for each individual. Specifically, panel **(A)** shows PAB16_hap1, **(B)** PAB16_hap2, **(C)** PPY17_hap1, and **(D)** PPY17_hap2. Additional details, including the corresponding information for PPY15, are provided in fig. S2. Note: Given differences in the size of the α-sat HOR array, each plot is scaled differently.

Available orangutan T2T assemblies(*21*), hereafter referred to as PPY_T2T (mPonPyg2-v2.1) and PAB_T2T (mPonAbe1-v2.1), show that both PAB and PPY chromosome 10, all in a canonical CEN/CEN configuration, have centromeres typically characterized by a single short stretch of monomeric α-sat DNA located within the canonical centromeric interval, spanning ∼116-164 kbp in PAB and ∼221 kbp in a single PPY haplotype(*21*). An exception is observed in the alternative PPY_T2T haplotype (mPonPyg2-v2.1_alt; PPY_T2T_alt), which harbors two distinct α-sat loci: a large and active HOR-containing array of ∼1.87 Mbp together with a smaller and inactive monomeric array of ∼209 kbp (tables S10-S11).

Using RepeatMasker(*23*) optimized for primates to annotate α-sat regions (ALR/alpha entries) in our three representative samples, we identified α-sat arrays on chromosome 10 in all six analyzed haplotypes, with striking heterogeneity in size, organization, and genomic context (Fig. 2, fig. S2, table S11). Array length varied by an order of magnitude, from ∼118 kbp to >2.1 Mbp, and substantially differed across homologous chromosomes within the same individual. In PAB16 and PPY15, the four haplotypes contained short α-sat arrays (<250 kbp) located within the canonical centromeric interval (detailed α-sat locations are provided in table S11). In contrast, PPY17 haplotype 2 (PPY17_hap2) uniquely harbored two distinct α-sat loci, exactly as the PPY_T2T_alt haplotype(*21*): a small array (∼212 kbp), present in both PAB and PPY orangutans, and a much larger (∼2.1 Mbp) array, located ∼2.4 Mbp upstream, revealing a complex, multi-array centromeric landscape. Importantly, α-sat arrays shorter than ∼250 kbp did not form continuous satellite blocks, but instead displayed a mosaic architecture, in which α-sat monomers, particularly at the edges, were extensively interspersed with LINE elements (Fig. 2).

Annotation of gene content across these regions revealed that the presence or absence of the expanded α-sat stretch does not alter the expression of flanking genes. Coding sequences remain intact across all haplotypes, as α-sat arrays are confined to nongenic, repetitive intervals. CENdetectHOR(*24*) analysis showed that only PPY17_hap2 contained an α-sat HOR structure (α-sat(+)_HOR), predominantly composed of a 12-mer with multiple variant HORs, which covers ∼1.1 Mbp, whereas all other haplotypes exhibited strictly monomeric α-sat organization (α-sat(+)_Mono) (Fig. 2, figs. S2-S3, tables S12-S13).

To assess the internal sequence organization, we generated dot plots for each region using ModDotPlot(*25*) with an 80% sequence identity threshold. In α-sat(+)_Mono regions, we identified a lower percent of sequence identity (<90%) than the one typical of great apes canonical centromeres(*21, 26*), independent of methylation state or centromere activity (Fig. 2). On the contrary, the α-sat(+)_HOR array in PPY17_hap2, exhibited high sequence identity (>95%), mirroring the long-range homogeneity observed in common primate centromeres (Fig. 2)(*21, 27*).

Putative neocentromeric regions were initially mapped based on synteny with the locus originally hypothesized by Tolomeo et al. (2017)(*19*). However, these coordinates were further refined and validated by integrating DNA methylation profiling (CDR-Finder) and CENP-A/C enrichment patterns (ChIP-Seq), precisely localizing the boundaries of functional neocentromeres. Functional activity was identified by profiling DNA methylation via CDR-Finder (v1.0.1,(*28*)), revealing that the hypomethylated region (CDR)(*29*) (table S11), indicative of centromeric activity, was restricted to the α-sat region only in PAB16_hap1 (120 kbp) and PPY17_hap2 (140 kbp in the α-sat(+)_HOR), while in PAB16_hap2, PPY15_hap1 and hap2, and PPY17_hap1, the CDR (100–120 kbp each) was in the neocentromeric region (Fig. 2, table S11). These patterns were consistent with CENP-A/C ChIP-seq enrichment, which mapped to the neocentromeric region in both PPY15 haplotypes. In contrast, haplotype-specific centromere activity was observed in the other two samples: in PAB16, haplotype 1 (hap1) showed CENP-A/C enrichment at the canonical centromere, whereas haplotype 2 (hap2) localized it to the neocentromeric region; similarly, in PPY17, hap1 was associated with the neocentromere and hap2 with the canonical centromere.

Based on α-sat presence, HOR organization, and functional annotation, we classified centromeric regions as α-sat(+)_Mono_active (PAB16_hap1), α-sat(+)_Mono_inactive (PAB16_hap2, PPY15_hap1, PPY15_hap2, PPY17_hap1), and α-sat(+)_HOR_active (PPY17_hap2 proximal array). At the neocentromeric regions, detailed sequence analysis revealed an almost complete absence of long-range sequence identity, with only sparse, highly diverged structures (<85% similarity) lacking α-sat DNA, even in haplotypes in which the neocentromere is functionally active (α-sat(−)_Neo active/inactive). Similarly, hypomethylation and CENP-A/CENP-C enrichment identified the active canonical centromeric regions (table S11).

As above-mentioned, the distinctive structural pattern identified in PPY17_hap2 was also observed upon analysis of the most recent Bornean orangutan T2T assembly (mPonPyg2-v2.0_pri; PPY_T2T)(*21*). To assess population prevalence of the α-sat(+)_HOR, we generated a 101 bp FISH probe targeting the α-sat HOR domain and screened the full collection of PPY (8) and PAB (10) individuals (Supplementary Text 5; fig. S4). Strong hybridization signals were detected on one chromosome 10 homolog in four PPY individuals (PPY12, PPY17, PPY19, PPY_T2T) (fig. S5), whereas PAB individuals lacked any signal, consistent with the absence of the large α-sat HOR array (figs. S6-S8; table S16). This finding suggests that the large α-sat HOR centromeric array is a species-specific feature of the PPY lineage but that it is also structurally polymorphic. Immuno-FISH and ChIP-Seq showed that when the large α-sat HOR organization is present, it functions as an active centromere. In the monomeric haplotypes, CENP-C signals were instead located at the α-sat(+)_Mono or α-sat(+)_Neo regions, depending on the hypomethylation profile (fig. S9).

Micro-C analysis showed that the overall 3D organization of chromosome 10 is highly similar across the three sequenced individuals (Supplementary Text 6; fig. S10). Zoomed-in Micro-HiC maps revealed consistent chromatin organization at the canonical centromere across all three cell lines, where the centromere exactly coincides with a discrete sub-TAD that overlaps the hypermethylated domain surrounding the centromere (fig. S11). In contrast, the neocentromeric locus in all samples coincides with a wide TAD boundary located within a B compartment (fig. S12). A small sub-TAD–like domain overlapping the active neocentromere, the region of hypomethylation and CENP-A deposition, is detectable in all cell lines (fig. S13). In PPY15, however, where the neocentromere is active on both alleles, this domain appears more strongly insulated, with reduced interactions with neighboring domains.

## Segmental duplications mediate the 3.6 Mbp HOR centromere deletion, and LINEs promote alphoid movement

To investigate the organization of chromosome 10 centromeres, we first annotated repetitive elements using an integrated repeat-masking framework and identified segmental duplications (SDs) >90% identical and >1 kbp in length (Supplementary Text 7). Notably, only the large centromeres, PPY_T2T_hap2 and PPY17_hap2 (Fig. 3A) that harbor the large tract of α-sat HORs (∼2 Mbp), were flanked by 252 kbp of SDs with ∼99.25% sequence identity, whereas haplotypes lacking the large centromeres had a single copy of the 252 kbp unit and, therefore, classified as unique sequence (Fig. 3A). This genomic architecture suggested that the large α-sat HOR tract was likely deleted evolutionarily by non-allelic homologous recombination (NAHR) by the directly oriented 252 kbp units (Fig. 3B).

**Fig. 3.**
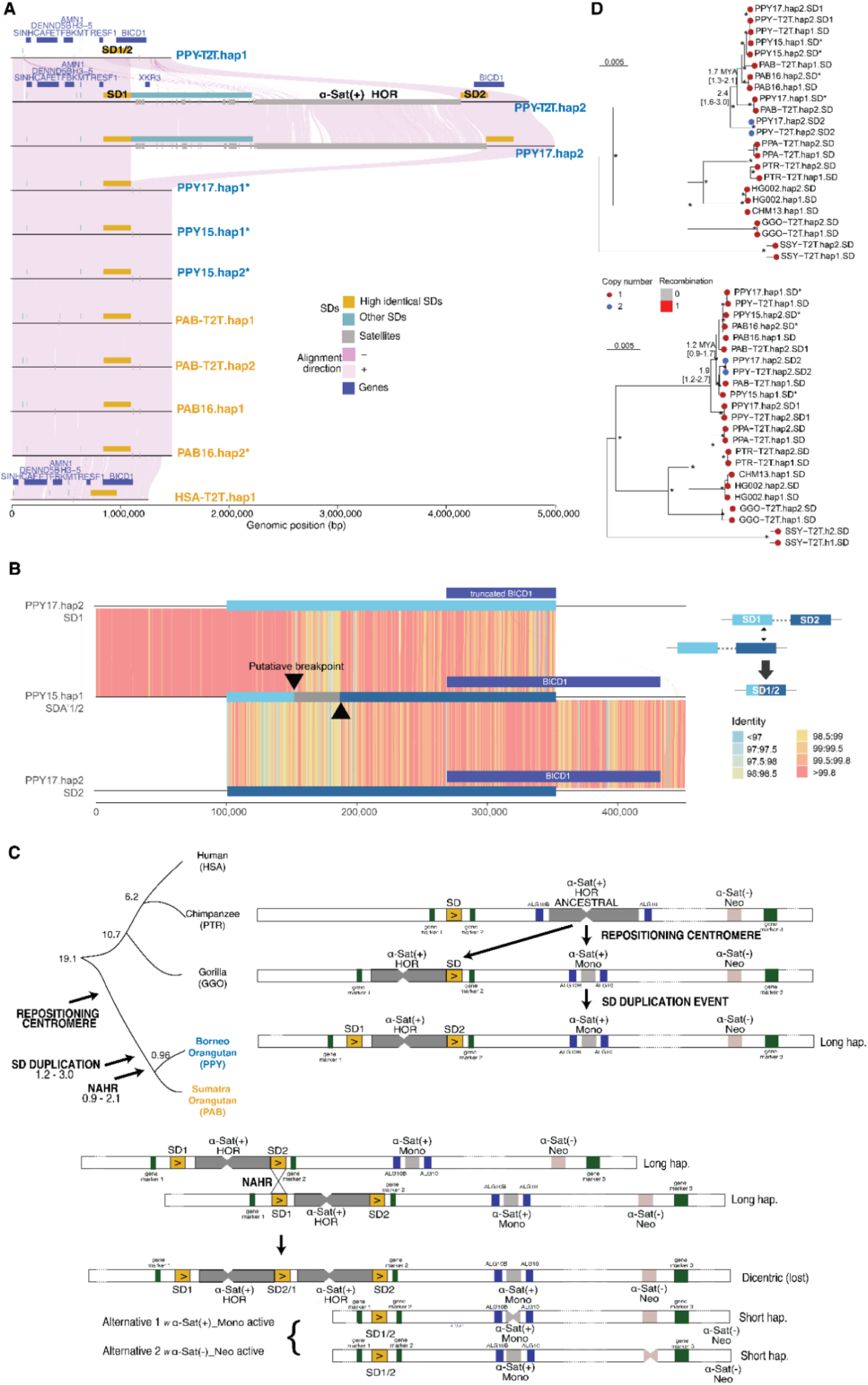
Segmental duplication (SD) mediating non-allelic homologous recombination (NAHR) and centromere deletion. **(A)** Local alignment (minimap2) of Sumatran (PAB) and Bornean (PPY) orangutan genomes with respect to the 252 kbp lineage-specific SD (orange) (SD1/2), along with gene, satellite, and other SD annotations. **(B)** Alignment of a recombined SD (SD1/2 of PPY15_hap1) compared to SD1 and SD2 of PPY17_hap2. The alignment blocks are binned into 1 kbp fragments and colored by sequence identity. The shift in sequence identity identifies the putative ancestral NAHR breakpoint interval (arrows and grey bar). **(C)** Putative model of the orangutan ancestral chr10 centromere, which, after a repositioning event, has originated the actual long haplotype. NAHR can then mediate the HOR deletion and the subsequent activation of the α-sat(+)_mono or of the neocentromere. **(D)** Maximum likelihood phylogenetic trees of the SD1/2 duplication. Two multiple sequence alignments were constructed from both the 5’ and 3’ ends of the SD from both PAB and PPY and using other nonhuman primates as an outgroup (PPA: bonobo, PTR: chimpanzee, HG002 & CHM13: humans, GGO: gorilla and SSY: siamang), which all carry a single copy of the 252 kbp unit (unique sequence). Among the orangutans, SD1 vs. SD2 copies are distinguished, as well as whether the haplotypes are the result of a recombination event (SD1/2) or represent the ancestral state (SD, SD1, and SD2). The coalescent times are reported in million years ago (Mya), and the interval is computed from 100 replicates. *haplotypes correspond to a neocentromere

The conservation of local gene order and flanking-sequence synteny across all haplotypes, along with the 3.6 Mbp size reduction observed in all neocentromere and PAB haplotypes (Fig. 3A), supports an SD-mediated deletion by NAHR rather than by independent rearrangements (Fig. 3C). In contrast, haplotypes lacking this deletion invariably display centromeric activity at the α-sat HOR array. Because NAHR between SDs would be predicted to create a hybrid locus composed of both SD1 and SD2 sequences, we compared the sequence identity from the single-copy 252 kbp unit of PPY15_hap1 to SD1 and SD2 from PPY17_hap2 (Fig. 3B). From the 5’ flanking sequence of the single-copy unit, we find >99.8% identical alignment synteny, shared with the left copy (SD1), extending to a recombination interval (marked by black arrows in Fig. 3B) characterized by a localized decrease in sequence identity. We hypothesize that this region represents a transition zone where the sequence shifts from SD1-specific to SD2-specific variants, identifying it as the likely site of the NAHR event that generated the single-copy unit (Fig. 3B) associated with neocentromeres.

To estimate the timing of the duplication and deletion event, we built two maximum-likelihood phylogenetic trees corresponding to 50 kbp sequences sampled from the left and right ends of the 252 kbp unit, respectively (Fig. 3D). Among the sequences of the duplicated pair, the coalescent times were estimated to be 2.4 (with a 1.6-3.0 confidence interval) Mya and 1.9 (1.2-2.7) Mya, for the left and right subsequences, respectively. This suggests that the duplication event likely occurred between 1.2 and 3.0 Mya, predating the speciation of the two orangutan species. As a second analysis, we estimated the coalescent time for all orangutan haplotypes bearing a single copy of the 252 kbp sequence. The deepest coalescent time between recombined copies was found to be 1.7 (1.3-2.1) Mya and 1.2 (0.9-1.7) Mya, selecting from the 5’ and 3’ end of the SD, respectively. This suggests that the deletion of chromosome 10 centromeres among the majority of orangutans occurred soon after the duplication event between 0.9-2.1 Mya.

Comparative analyses across primate T2T assemblies showed that the genomic domain orthologous to the 252 kbp SD unit is conserved on the chromosome homologous to the orangutan chromosome 10 in chimpanzee, bonobo, gorilla, Sumatran orangutan, human, and macaque (Supplementary Text 7). However, in these species, the orthologous sequence is present as a single-copy locus, whereas only the Bornean orangutan haplotypes carrying the large HOR-centromere show the derived duplicated configuration (SD1/SD2) flanking the α-sat array. This indicates that the duplicated architecture is orangutan-specific and supports a mechanism in which the SD pair arose after the ancestral primate single-copy state diverged, thereby creating the substrate for subsequent NAHR-mediated deletion.

Chromosome 10 synteny analysis further supports this mechanism. The short monomeric α-sat array that is retained in most orangutan haplotypes is flanked by *ALG10B* and *ALG10* (Supplementary Text 7-8; figs. S14-S15). These gene markers flank the syntenic centromere across the primate lineage(*21*) suggesting it is the ancestral location (figs. S14-S15). This suggests that the present-day short α-sat locus likely corresponds to the relic of an older ancestral centromere. Under this scenario, a centromere repositioning event in the orangutan ancestral lineage first displaced the functional centromere away from the ancestral *ALG10B*–*ALG10* interval, leaving behind only a reduced monomeric α-sat remnant in orangutans, while a larger HOR-containing array became established ∼2 Mbp away. The subsequent emergence of the 252 kbp SD pair would then have created the substrate for NAHR, ultimately leading to subsequent deletion of the intervening HOR centromere.

Gene annotation and Iso-Seq analysis also revealed that the duplicated 252 kbp unit encompasses expressed *BICD1* gene models (Supplementary Text 8). Two transcript isoforms were supported by more than 10 full-length reads, whereas a third low-abundance model showed a disrupted predicted open reading frame. The longer *BICD1* isoform mapped specifically to the downstream duplicated copy (SD2), whereas a shorter isoform was detected from both SD1 and SD2 in PPY17_hap2, indicating that the duplication generated paralog-specific transcripts and amino acid substitutions (figs. S16-S17), although AlphaFold-based structural modeling(*30*) did not indicate major folding differences. These data show that the SD architecture associated with centromere remodeling also intersects a transcriptionally active gene locus, potentially creating additional functional consequences beyond centromere instability.

To investigate the evolutionary history of the α-sat–associated region, we analyzed long LINE elements (>5 kbp), as molecular fossils of past transposable element activity (Supplementary Text 7). We found enrichment for L1, including L1P1, L1PA2, L1PA3, and L1PA4 (table S21), although their distributions differed among lineages (tables S22-26). In detail, in human and chimpanzee L1P1 and multiple L1PA elements were predominantly present, whereas bonobo showed a majority of L1PA2 together with L1PA3 and L1PA4; gorilla showed only L1PA3 and L1PA4 elements, while in the PPY_T2T_alt only L1PA3 elements were detected within the α-sat region. The differential distribution of LINE subfamilies across species suggests lineage-specific waves of LINE activity. In particular, the enrichment of relatively younger L1 elements(*31*) within orangutan α-sat regions supports a mechanism in which centromere repositioning occurred in a genomic environment remodeled by recent transposable element insertions.

## Introgression and incomplete lineage sorting shape the evolution of the neocentromeric region

Phylogenetic and coalescent analyses indicate that the evolutionary history of the neocentromeric region is uncoupled from the remainder of chromosome 10 (Supplementary Text 9). Phylogenetic analysis of entire chromosome haplotypes reflected the established divergence between PPY and PAB species, grouping the six PPY and the four PAB haplotypes into two well-supported clades (fig. S18), confirming the separation between the Bornean and Sumatran groups observed in the PCA (Fig. 1D). In contrast, the neocentromeric region exhibits a strikingly distinct evolutionary pattern: the PAB haplotype carrying the neocentromere (PAB16_hap2) clusters within the PPY clade, whereas non-neocentromeric PAB haplotypes form the expected separate cluster (Fig. 4A). Within PPY, it is not possible to discriminate between neocentromeric and non-neocentromeric individuals, indicating that the presence of the neocentromere does not affect their phylogenetic relationships.

**Fig. 4.**
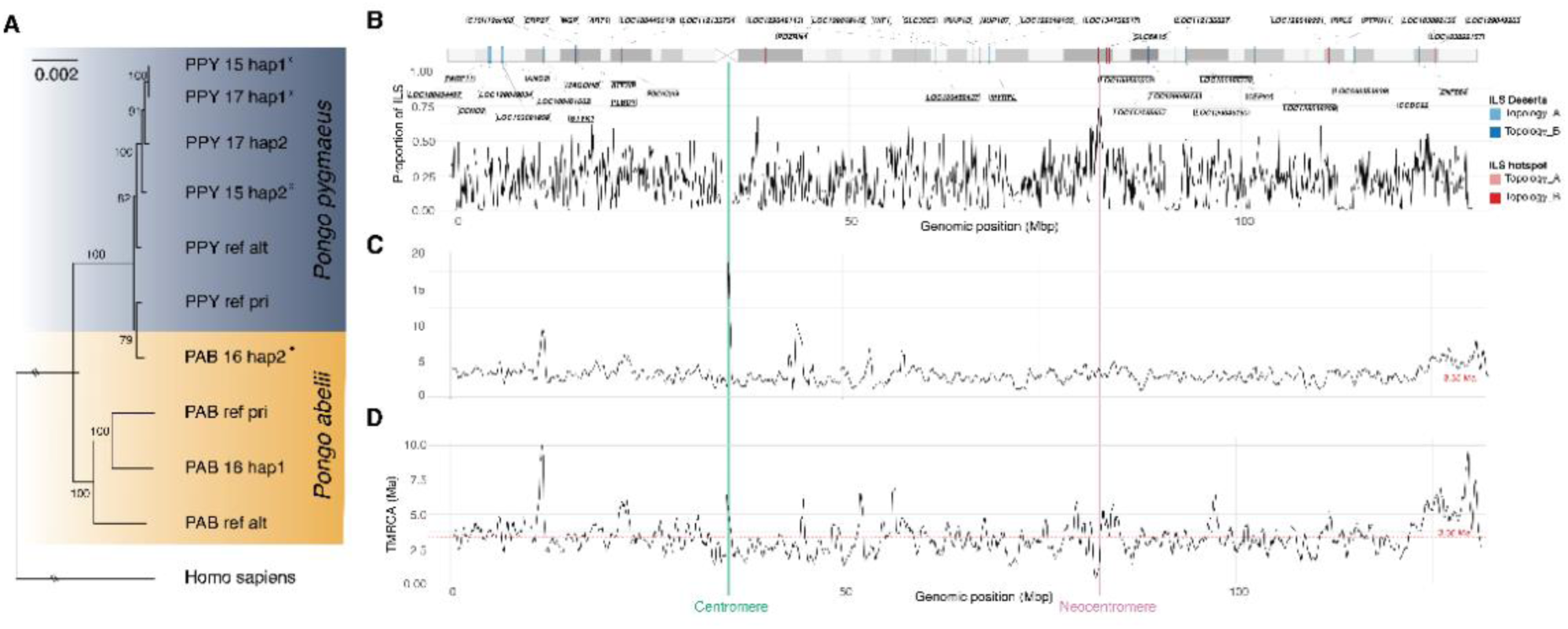
Phylogenetic relationships, incomplete lineage sorting (ILS), and TMRCA patterns across orangutan neocentromeres. **(A)** Maximum-likelihood phylogenetic tree inferred from the neocentromeric region, showing relationships among *Pongo* haplotypes; orange and blue indicate *PAB* and *PPY* haplotypes, respectively. *indicates haplotypes with a neocentromere. Numbers at nodes indicate bootstrap support. *Homo sapiens* was used as the outgroup. **(B)** Gene annotation of the 20 windows with the highest (hotspot, red) and lowest (desert, blue) ILS proportion. **(C)** Sliding-window estimates of time to the most recent common ancestor (TMRCA) along chromosome 10 in the comparison between PAB16_hap1 (a non-neocentromeric *PAB* haplotype) and PPY15_hap1 (a neocentromeric *PPY* haplotype). **(D)** Sliding-window estimates of TMRCA along chromosome 10 in the comparison between neocentromeric haplotypes from both species (PAB16_hap2 vs. PPY15_hap1). Windows of 500 kbp were analysed with a 100 kbp step. The red line indicates the mean TMRCA across the entire chromosome. Green and red vertical lines indicate the centromeric and the neocentromeric regions, respectively.

The discordance is further supported by time to the most recent common ancestor (TMRCA) estimates. Comparisons involving the non-neocentromeric haplotypes of PAB and PPY (the latter carrying the active neocentromere) yield coalescent times of ∼3.3–3.6 Mya, which predate the species divergence and are consistent with the average TMRCA of the entire chromosome (∼3.4 Mya) (Fig. 4C-D). By contrast, the neocentromeric PAB haplotype shows markedly more recent coalescence with both PPY haplotypes (∼0.32–0.42 Mya). As this timeframe significantly postdates the species divergence, it suggests a possible secondary introgression of the neocentromeric region from PPY into PAB. This event likely introduced the neocentromeric region, harbouring an already active neocentromere or one predisposed to activation, into the PAB genomic background. Selection scans along chromosome 10 further highlight lineage-specific evolutionary dynamics (Supplementary Text 9; fig. S19). In PPY, both canonical centromeres and neocentromeres show evidence of strong selective sweeps, whereas PAB exhibits no signature of selection, indicating differential selective pressures on centromeric regions between the two species.

Incomplete lineage sorting (ILS) pattern along orangutan chromosome 10 reveals a pronounced elevation at the neocentromeric region (Supplementary Text 10): across chromosome 10, the mean ILS proportion is ∼21.8%, whereas the neocentromeric region exhibits an ILS of ∼60.9%, strongly favoring a topology in which the PPY15 neocentromeric haplotype is closer to PAB16 neocentromeric haplotype than to the non-neocentromeric one. (Topology B; fig. S20). Window-based coalescent analysis confirms a shallow divergence between PAB haplotypes and a deep coalescent with PPY15 (fig. S21).

Gene annotation within ILS hotspots and deserts indicates that the neocentromeric region resides in a gene desert (Fig. 4B). Nearby hotspots similarly exhibit minimal genic content, with only two windows in proximity to the neocentromere harboring *SLC6A15* and *LOC112136027*, suggesting that neocentromere formation and maintenance may be facilitated by gene-poor, structurally permissive chromosomal contexts(*5, 14, 16, 32–37*) (fig. S22; table S30).

## Discussion

Neocentromeres and evolutionary new centromeres (ENCs) provide a powerful system to investigate how centromeric function emerges, stabilizes, and is maintained during evolution. Although neocentromeres have been reported in both clinical(*38*) and evolutionary contexts(*11, 16, 34, 39*), we still know very little about the molecular architecture of neocentromeric regions and the mechanisms governing their formation, maturation, and long-term evolutionary stability. By leveraging orangutan populations where a neocentromere segregates at roughly 26% allele frequency, we combined haplotype-resolved assemblies with epigenetic and population-level analyses to provide a framework linking centromere repositioning, structural variation, and long-term evolutionary outcomes.

In most primates, centromeres are characterized by large, homogeneous arrays of α-sat DNA, which have long been considered a defining feature of centromeric identity and function(*40*). A striking feature of orangutan chromosome 10 is the widespread absence of canonical α-sat HOR arrays across most haplotypes. Instead, the majority of chromosomes harbor unexpectedly short α-sat arrays (roughly 120 kbp), organized as monomers, and heavily interspersed with LINE elements. These features are inconsistent with the classical model of primate centromeres(*41–43*), suggesting a substantial deviation from the canonical architecture. A single exception is represented by one haplotype (PPY17_hap2), which still carries a large 2 Mbp α-sat HOR array alongside a downstream region containing short α-sat arrays found in the other orangutan genomes. Population-scale FISH and immuno-FISH analyses in both orangutan species show that this HOR organization is present in multiple Bornean individuals, including the alternative PPY reference haplotype(*21*), but absent in Sumatran orangutans, indicating that this architecture constitutes a large structural polymorphism confined to the PPY species. CENP-C enrichment confirms that these HOR arrays are functionally active when present. However, their restricted distribution indicates that a canonical HOR-based organization is not required for centromere function but rather represents one of several alternative centromeric states.

The SD analysis revealed an extraordinary genomic context surrounding the α-sat HOR array. The large α-sat HOR is flanked by two approximately 252 kbp orangutan SDs (∼99.25% identity) in direct orientation that create a substrate for NAHR(*44*). Our data suggest that NAHR-mediated deletion of the intervening α-sat HOR array has generated shorter haplotypes lacking a canonical centromere. Such rearrangements would be expected to produce unstable chromosomal configurations (e.g., dicentric chromosomes, Fig. 4), thereby imposing strong selective pressure for the rapid establishment of an alternative centromeric site. In this context, both the activation of residual α-sat monomers and the emergence of neocentromeres can be interpreted as compensatory mechanisms that restore chromosome stability following structural disruption. Notably, this structural context intersects a transcriptionally active locus. The SD unit encompasses the gene *BICD1* (Fig. 2 and Fig. S7.1), which encodes a coiled-coil adaptor involved in dynein-mediated vesicle transport through interaction with Rab6(*45*), with distinct transcript isoforms arising from the two copies. This indicates that centromere-associated rearrangements can also produce local transcriptional consequences and lead to the emergence of new/modified genes even if only transiently within the orangutan lineage.

Comparative analysis indicates that the short monomeric α-sat locus present in most orangutan haplotypes is flanked by *ALG10B* and *ALG10*, a configuration conserved across primates and associated with ancestral centromeric domains (fig. S14). This strongly suggests that the original chromosome 10 centromere resided at this interval and was subsequently repositioned approximately 2 Mbp upstream, where a large HOR-containing array persists in a subset of PPY haplotypes. In this framework, the short α-sat locus represents a residual vestige of the ancestral centromere, whereas the large HOR array corresponds to a later active centromeric state that subsequently became susceptible to SD-mediated NAHR. Together, these observations indicate that chromosome 10 has undergone at least three successive transitions: centromere repositioning, SD birth, and NAHR-mediated centromere loss, followed by stabilization of alternative centromeric states, including α-sat-poor canonical centromeres and neocentromeres.

The distribution of LINE elements further supports this model. In humans, chimpanzees, and gorillas, the active centromere lies between the *ALG10B*-*ALG10* interval and is enriched in older L1PA4/L1PA3 elements, with L1PA4 age (18 Mya) overlapping the divergence time of great apes(*31*). In orangutans, centromeric activity has shifted outside this region and is enriched for solely younger L1PA3 elements (12.8 Mya)(*31*), indicating that repositioning occurred within a genomic context shaped by subsequent L1PA3 expansion, after divergence from the African ape–human lineage(*31*). Together, these observations support a model in which centromere repositioning on chromosome 10 was mediated by the movement and/or reorganization of a composite LINE–α-sat block rather than by the relocation of centromeric chromatin alone, as recently reported in human and *Opsariichthys*(*46, 47*). In this framework, the interval between *ALG10B* and *ALG10* represents the ancestral centromere-associated domain, whereas the orangutan lineage experienced a secondary displacement of the LINE–α block, followed by lineage-specific amplification, structural remodeling, and stabilization of alternative centromeric states.

Across chromosome 10 orangutan haplotypes, most of them appear to have evolved toward a derived state in which loss of the canonical α-sat HOR array is accompanied by epigenetic repositioning of centromeric function to alternative genomic locations. The neocentromeric regions identified here lack α-sat DNA yet display localized hypomethylation and CENP-A/C enrichment, indicating fully functional centromeric activity. These observations reinforce the view that centromere identity is determined primarily by epigenetic signatures rather than being dictated by a strict DNA-sequence blueprint(*12, 19, 48–50*). Notably, these neocentromeres consistently arise within gene-poor regions, supporting the idea that such genomic contexts provide permissive substrates for centromere repositioning by minimizing functional constraints(*11, 13, 39, 51*). Micro-C analysis further shows that the neocentromeric region overlaps with TAD-like chromatin structure present in all samples, which adopts a more discrete, insulated configuration when neocentromeric activity is established, suggesting that local 3D architecture may facilitate and stabilize centromere function. Despite their stability, these neocentromeres have not transitioned into an ENC throughout the acquisition of α-sat arrays, as observed in other systems(*7, 16, 52–54*). This suggests that satellite seeding is not an inevitable outcome of centromere activity but may instead be constrained by local structural or epigenetic features that limit satellite expansion at this particular locus.

At the population level, the evolutionary history of the neocentromeric region seems decoupled from the rest of the chromosome. Phylogenetic analyses indicate that the neocentromere likely arose within the Bornean lineage and was subsequently introgressed into Sumatran orangutans, consistent with known patterns of gene flow between these species(*55–57*). This introgression may have introduced a genomic background permissive to neocentromere activation, potentially influencing chromatin state and centromere competence in the Sumatran lineage. This hypothesis is supported by distinct and localized selection signatures at both centromeric and neocentromeric sites in PPY but not in PAB. Such lineage-specific dynamics are consistent with the “centromere paradox,” in which essential centromeric function is maintained despite the rapid turnover of centromeric DNA(*58, 59*). Models of centromere drive predict episodic selective sweeps arising from meiotic competition, especially under female meiotic asymmetry(*58, 60*). In this context, the selective sweep observed in PPY suggests that the neocentromere activation is relatively recent and may still be undergoing functional optimization.

These findings support a dynamic view of centromere biology(*27, 58*), in which canonical centromeres, inactive remnants, and fully functional neocentromeres can coexist within populations, and even within single individuals, highlighting the remarkable plasticity of centromeric identity. Crucially, this plasticity demonstrates that faithful chromosome segregation can be maintained despite extensive turnover of the underlying DNA sequence. By leveraging near-complete assemblies and integrated epigenetic analyses, our study provides a framework for understanding how centromere identity is established, maintained, and reshaped during evolution. Future experimental-induced functional perturbations, such as CRISPR-Cas9 depletions, will be essential to dissect the relative contributions of DNA sequence and epigenetic regulation in defining centromere function.

## Acknowledgments

We thank Tonia Brown for careful proofreading of the manuscript, and Ivan Alexandrov and Rachel O’Brein for insightful discussions and valuable feedback.

## Funding

This research was funded by the National Recovery and Resilience Plan (NRRP), Mission 4, Component 2, Investment 1.1, Call for tenders No. 104, published on 2 February 2022 by the Italian Ministry of University and Research (MUR), funded by the European Union—NextGenerationEU—Project Title ‘Telomere-to-telomere sequencing: the new era of Centromere and neocentromere eVolution (CenVolution)’, CUP H53D23003260006, grant assignment decree no. 1015 adopted on 7 July 2023 by the Italian MUR (MV, BT, FMa). Research reported in this publication was supported, in part, by the National Human Genome Research Institute of the National Institutes of Health (NIH) under grants R01HG002385 and R01HG010169 (to EEE). The content is solely the responsibility of the authors and does not necessarily represent the official views of the NIH. EEE is an investigator of the Howard Hughes Medical Institute.

## Author contributions

Conceptualization: MV, BT, FMa, EEE

Methodology: LdG, DAY, FMo, BT, LP, DN, AD, FKM

Software: LdG, DAY, JW

Investigation: LdG, DAY, RM, LP, FR, FP, AD, KH, KMM

Formal analysis: LdG, DAY, FMo, FP, AD, RM, LP, FR, DN, CP, KKO

Data curation: LdG, DAY, AD

Validation: LdG, DAY, MV

Visualization: LdG, DAY, FA, EEE, GAL, MV

Supervision: MV, BT, EEE, GAL

Funding acquisition: MV, BT, EEE, Fma

Writing – original draft: LdG, DAY, LP, FR, RM, AD, FP, Fma

Writing – review & editing: LdG, MV, EEE, GAL, CRC, FA, FMo, FKM

## Competing interests

EEE is a scientific advisory board (SAB) member of Variant Bio, Inc. All other authors have no affiliations with or involvement in any organization or entity with any financial or non-financial interest in the subject matter or materials discussed in this manuscript.

## Data, code, and materials availability

All data and materials generated in this study are available in the main text or the supplementary materials to the research community for the purpose of reproducing and extending the analyses. The chromosome 10 haplotype-resolved assemblies generated in this study have been deposited in NCBI under BioProject accession numbers PRJNA1442736 (PAB16_hap1), PRJNA1442739 (PAB16_hap2), PRJNA1442737 (PPY15_hap1), PRJNA1442744 (PPY15_hap2), PRJNA1442745 (PPY17_hap1), and PRJNA1442746 (PPY17_hap2). Corresponding BioSample accessions are SAMN56359922 (PAB16), SAMN56359927 (PPY15), and SAMN56360226 (PPY17). At the time of submission, some records are undergoing processing and validation at NCBI; all data will be publicly accessible upon completion of the submission process. All other data supporting the findings of this study are available within the main text and the Supplementary Materials or from the corresponding author upon reasonable request.

## Supplementary Materials

Supplementary Text Figs. S1 to S22

Tables S1 to S30

References (*61-118*)

## Reference

1. L. E. Kursel, H. S. Malik, Centromeres. Curr Biol 26, R487–R490 (2016).

2. C. Y. Y. Wong, B. C. H. Lee, K. W. Y. Yuen, Epigenetic regulation of centromere function. Cell Mol Life Sci 77, 2899–2917 (2020).

3. C. M. Wade et al., Genome sequence, comparative analysis, and population genetics of the domestic horse. Science 326, 865–867 (2009).

4. Z. Gong et al., Repeatless and repeat-based centromeres in potato: implications for centromere evolution. Plant Cell 24, 3559–3574 (2012).

5. K. Wang, Y. Wu, W. Zhang, R. K. Dawe, J. Jiang, Maize centromeres expand and adopt a uniform size in the genetic background of oat. Genome Res 24, 107–116 (2014).

6. F. M. Piras et al., Phylogeny of horse chromosome 5q in the genus Equus and centromere repositioning. Cytogenet Genome Res 126, 165–172 (2009).

7. M. Rocchi, N. Archidiacono, W. Schempp, O. Capozzi, R. Stanyon, Centromere repositioning in mammals. Heredity (Edinb) 108, 59–67 (2012).

8. L. Carbone et al., Evolutionary movement of centromeres in horse, donkey, and zebra. Genomics 87, 777–782 (2006).

9. L. E. Voullaire, H. R. Slater, V. Petrovic, K. H. Choo, A functional marker centromere with no detectable alpha-satellite, satellite III, or CENP-B protein: activation of a latent centromere? Am J Hum Genet 52, 1153–1163 (1993).

10. D. J. Amor et al., Human centromere repositioning “in progress”. Proc Natl Acad Sci U S A 101, 6542–6547 (2004).

11. M. Ventura et al., Recurrent sites for new centromere seeding. Genome Res 14, 1696–1703 (2004).

12. S. J. Hoyt et al., Haplotype-Resolved Genomics Reveals Conserved Chromatin Architecture and Epigenetic Constraints of Human Neocentromeres. bioRxiv, (2025).

13. M. Ventura et al., Neocentromeres in 15q24-26 map to duplicons which flanked an ancestral centromere in 15q25. Genome Res 13, 2059–2068 (2003).

14. M. F. Cardone et al., Independent centromere formation in a capricious, gene-free domain of chromosome 13q21 in Old World monkeys and pigs. Genome Biol 7, R91 (2006).

15. M. Ventura, N. Archidiacono, M. Rocchi, Centromere emergence in evolution. Genome Res 11, 595–599 (2001).

16. M. Ventura et al., Evolutionary formation of new centromeres in macaque. Science 316, 243–246 (2007).

17. G. Giannuzzi et al., Hominoid fission of chromosome 14/15 and the role of segmental duplications. Genome Res 23, 1763–1773 (2013).

18. N. Altemose et al., Complete genomic and epigenetic maps of human centromeres. Science 376, eabl4178 (2022).

19. D. Tolomeo et al., Epigenetic origin of evolutionary novel centromeres. Sci Rep 7, 41980 (2017).

20. D. P. Locke et al., Comparative and demographic analysis of orang-utan genomes. Nature 469, 529–533 (2011).

21. D. Yoo et al., Complete sequencing of ape genomes. Nature 641, 401–418 (2025).

22. M. Rautiainen et al., Telomere-to-telomere assembly of diploid chromosomes with Verkko. Nat Biotechnol 41, 1474–1482 (2023).

23. S. Tempel, Using and understanding RepeatMasker. Methods Mol Biol 859, 29–51 (2012).

24. A. Daponte, et al., CENdetectHOR: a comprehensive tool for CENtromere profiling and HOR detection. bioRxiv, 2025.2001.2007.631657 (2025).

25. A. P. Sweeten, M. C. Schatz, A. M. Phillippy, ModDotPlot-rapid and interactive visualization of tandem repeats. Bioinformatics 40, (2024).

26. G. A. Logsdon et al., The structure, function and evolution of a complete human chromosome 8. Nature 593, 101–107 (2021).

27. G. A. Logsdon et al., The variation and evolution of complete human centromeres. Nature 629, 136–145 (2024).

28. F. K. Mastrorosa et al., Identification and annotation of centromeric hypomethylated regions with CDR-Finder. Bioinformatics 40, (2024).

29. A. Gershman et al., Epigenetic patterns in a complete human genome. Science 376, eabj5089 (2022).

30. M. G. Krokidis et al., AlphaFold3: An Overview of Applications and Performance Insights. Int J Mol Sci 26, (2025).

31. H. Khan, A. Smit, S. Boissinot, Molecular evolution and tempo of amplification of human LINE-1 retrotransposons since the origin of primates. Genome Res 16, 78–87 (2006).

32. C. Ketel et al., Neocentromeres form efficiently at multiple possible loci in Candida albicans. PLoS Genet 5, e1000400 (2009).

33. K. A. Maggert, G. H. Karpen, The activation of a neocentromere in Drosophila requires proximity to an endogenous centromere. Genetics 158, 1615–1628 (2001).

34. O. J. Marshall, A. C. Chueh, L. H. Wong, K. H. Choo, Neocentromeres: new insights into centromere structure, disease development, and karyotype evolution. Am J Hum Genet 82, 261–282 (2008).

35. S. Nasuda, S. Hudakova, I. Schubert, A. Houben, T. R. Endo, Stable barley chromosomes without centromeric repeats. Proc Natl Acad Sci U S A 102, 9842–9847 (2005).

36. A. M. Olszak et al., Heterochromatin boundaries are hotspots for de novo kinetochore formation. Nat Cell Biol 13, 799–808 (2011).

37. H. Su et al., Dynamic chromatin changes associated with de novo centromere formation in maize euchromatin. Plant J 88, 854–866 (2016).

38. D. J. Amor, K. H. Choo, Neocentromeres: role in human disease, evolution, and centromere study. Am J Hum Genet 71, 695–714 (2002).

39. O. Capozzi et al., Evolutionary descent of a human chromosome 6 neocentromere: a jump back to 17 million years ago. Genome Res 19, 778–784 (2009).

40. L. L. Sullivan, K. Chew, B. A. Sullivan, α satellite DNA variation and function of the human centromere. Nucleus 8, 331–339 (2017).

41. I. Alexandrov, A. Kazakov, I. Tumeneva, V. Shepelev, Y. Yurov, Alpha-satellite DNA of primates: old and new families. Chromosoma 110, 253–266 (2001).

42. C. Alkan et al., Genome-wide characterization of centromeric satellites from multiple mammalian genomes. Genome Res 21, 137–145 (2011).

43. C. Alkan et al., Organization and evolution of primate centromeric DNA from whole-genome shotgun sequence data. PLoS Comput Biol 3, 1807–1818 (2007).

44. P. Stankiewicz, J. R. Lupski, Genome architecture, rearrangements and genomic disorders. Trends Genet 18, 74–82 (2002).

45. Y. Liu et al., Bicaudal-D uses a parallel, homodimeric coiled coil with heterotypic registry to coordinate recruitment of cargos to dynein. Genes Dev 27, 1233–1246 (2013).

46. L. L. Sullivan, K. A. Maloney, A. J. Towers, S. G. Gregory, B. A. Sullivan, Human centromere repositioning within euchromatin after partial chromosome deletion. Chromosome Res 24, 451–466 (2016).

47. J.-H. Tai et al., Transposons accelerate chromosomal speciation by centromere expansion and chromosome fission. bioRxiv, 2025.2002.2013.638213 (2025).

48. G. A. Logsdon, M. R. Vollger, E. E. Eichler, Long-read human genome sequencing and its applications. Nat Rev Genet, (2020).

49. M. Murillo-Pineda et al., Induction of spontaneous human neocentromere formation and long-term maturation. J Cell Biol 220, (2021).

50. C. Salinas-Luypaert et al., DNA methylation influences human centromere positioning and function. Nat Genet 57, 2509–2521 (2025).

51. M. F. Cardone et al., Evolutionary history of chromosome 11 featuring four distinct centromere repositioning events in Catarrhini. Genomics 90, 35–43 (2007).

52. K. Nagaki et al., Sequencing of a rice centromere uncovers active genes. Nat Genet 36, 138–145 (2004).

53. H. Yan et al., Genomic and genetic characterization of rice Cen3 reveals extensive transcription and evolutionary implications of a complex centromere. Plant Cell 18, 2123–2133 (2006).

54. H. Zhao et al., Gene Expression and Chromatin Modifications Associated with Maize Centromeres. G3 (Bethesda) 6, 183–192 (2015).

55. C. Becquet, M. Przeworski, A new approach to estimate parameters of speciation models with application to apes. Genome Res 17, 1505–1519 (2007).

56. M. P. Mattle-Greminger et al., Genomes reveal marked differences in the adaptive evolution between orangutan species. Genome Biol 19, 193 (2018).

57. A. Nater et al., Morphometric, Behavioral, and Genomic Evidence for a New Orangutan Species. Curr Biol 27, 3487–3498.e3410 (2017).

58. S. Henikoff, K. Ahmad, H. S. Malik, The centromere paradox: stable inheritance with rapidly evolving DNA. Science 293, 1098–1102 (2001).

59. H. S. Malik, S. Henikoff, Major evolutionary transitions in centromere complexity. Cell 138, 1067–1082 (2009).

60. M. A. Lampson, B. E. Black, Cellular and Molecular Mechanisms of Centromere Drive. Cold Spring Harb Symp Quant Biol 82, 249–257 (2017).

