## Supplemental Text and Figures for "A segmental duplication-mediated deletion leads to neocentromere formation in orangutans"

Science
AAAS

### Supplementary Materials for

#### A segmental duplication-mediated deletion leads to neocentromere formation in orangutans

**Author:** Luciana de Gennaro[1], DongAhn Yoo[2], Letizia Pistacchia[3], Rosaria Magrone[1], Alessia Daponte[1], Francesco Perrone[1], Francesco Ravasini[3,4], F. Kumara Mastrorosa[2], Keisuke K. Oshima[8], Cesare Polano[5], Kendra Hoekzema[2], Katherine M. Munson[2], Julie Wertz[2], Fabio Marroni[5], Claudia R. Catacchio[1], Francesca Antonacci[1], Daan Noordermeer[6], Francesco Montinaro[1,7], Glennis A. Logsdon[8], Beniamino Trombetta[3], Evan E. Eichler[2,9], Mario Ventura[1]*

**Affiliations:**
[1] Department of Biosciences, Biotechnology and Environment, University of Bari, Bari, 70124, Italy
[2] Department of Genome Sciences, University of Washington School of Medicine, Seattle, WA 98195, USA
[3] Department of Biology and Biotechnologies 'Charles Darwin', Sapienza University of Rome, 00185 Rome, Italy
[4]BiRC – Section for Bioinformatics and Computational Biology, Aarhus University, 8000 Aarhus, Denmark
[5]Department of Agricultural, Food, Environmental and Animal Sciences, Via delle Scienze 206, University of Udine, 33100, Udine, Italy
[6]Université Paris-Saclay, CEA, CNRS, Institute for Integrative Biology of the Cell (I2BC), 91190 Gif-sur-Yvette, France
[7]Institute of Genomics, University of Tartu, Tartu, Estonia
[8]Department of Genome Sciences, University of Washington School of Medicine, Seattle, WA, USA; Present address: Department of Genetics, Epigenetics Institute, Perelman School of Medicine, University of Pennsylvania, Philadelphia, PA, USA
[9]Howard Hughes Medical Institute, University of Washington, Seattle, WA 98195, USA

**The PDF file includes:**

**Supplementary Text**
**Figs. S1 to S22**

**Other Supplementary Material for this manuscript includes the following:**

**Tables S1 to S30**

#### Supplementary Text

#### 1. Samples

##### Methods

We analyzed 17 orangutan lymphoblastoid cell lines (LCLs) (Table S1), comprising six *Pongo pygmaeus* (PPY) and ten *Pongo abelii* (PAB) and one hybrid PPYxPAB individuals. All cell lines were cultured in RPMI medium supplemented with 16% fetal bovine serum (FBS), 1% L-glutamine (L-Glut), and 1% penicillin-streptomycin (PenStrep).

**Cytogenetics characterization**

We cytogenetically characterized all 17 cell lines to determine species, sex, and the neocentromere status. Species identification was confirmed by examining the short arm of chr2_hsa3, which differs in length between the two species(*61*). Sex determination and neocentromere presence/absence were assessed using fluorescent *in situ* hybridization (FISH) with probes specific for the PAR region (RP11-990G10) and the q arm of chromosome 10 (RP11-107L10; human hg38 location chr12:124,381,868-124,562,842), respectively. FISH experiments were conducted on metaphase spreads prepared from lymphoblastoid cell lines using the standard procedure(*62*). DNA for FISH probes was extracted from selected BACs using the Bio-Rad Quantum Prep Plasmid Miniprep Kit (Cat. no. 7326100). The FISH protocol followed was as described by Ventura et al. (2004)(*11*). Briefly, 200 ng of each DNA probe, labeled via nick-translation with Cy3-dUTP, was precipitated with human Cot-1 DNA by alcohol precipitation, denatured at 70°C for 2 minutes, and hybridized overnight at 37°C. Post-hybridization washes were performed at 60°C in 0.1× SSC (three times, high stringency). Slides were counterstained with DAPI, producing a Q-banding pattern. Fluorescent signals from Cy3 and DAPI were captured separately using a Leica DMRXA epifluorescence microscope equipped with a cooled CCD camera (Princeton Instruments) and recorded as greyscale images. Final image pseudo-colorization and merging were performed in Adobe PhotoshopTM.

##### Results

Cytogenetic screening of 17 orangutan LCLs identified six *P. pygmaeus* (PPY), ten P. abelii (PAB), and one hybrid individual (PPY19) heterozygous for a pericentric inversion on chromosome 2(*61, 63*). FISH with the specific BAC probes RP11-990G10 and RP11-107L10(*64*) to assess chromosomal sex and neocentromeric status on chromosome 10 (Fig. S1) revealed that our cohort includes seven male and ten female individuals across both species. Among Sumatran individuals, seven were female, and three were male; in the Bornean group, three were female, and three were male and the hybrid resulted male. Additional cytogenetic evidence on sex chromosomes revealed three individuals with chromosomal abnormalities: one Sumatran male (PAB10) with XXY karyotype, one Sumatran female (PAB13) showing mosaicism (XX/XXX), and another Sumatran male (PAB16) with approximately 60% mosaic trisomy of chromosome 10.

FISH with the RP11-107L10 probe identified three distinct centromere configurations: ten individuals were homozygous for the canonical centromere on chromosome 10 (CEN10); seven samples were heterozygous for the neocentromere on chromosome 10 (NEO10); and only one sample, PPY15, was homozygous for NEO10. Specifically, among Sumatran orangutans, five were CEN/NEO, and the remaining five were CEN/CEN. In the Bornean group, three individuals carried the neocentromere (two CEN/NEO and one NEO/NEO), while four were CEN/CEN. The hybrid individual was a CEN/CEN. In parallel, we confirmed the CEN/CEN configuration and male sex of the AG05252 (PPY_T2T) reference individual (Fig. 1C) and therefore used it as a fully canonical Bornean reference genome in subsequent analyses. A detailed overview of the orangutan cohort is provided in Table S1.

#### 2. Genomic sequencing and PCA

##### Methods

All 17 orangutan lymphoblastoid cell lines were processed for Illumina sequencing of genomic DNA and RNA-seq. Additionally, only for three of these cell lines, PAB16, PPY15, and PPY17, was long-read sequencing performed, starting from high-molecular-weight (HMW) DNA. These three lines underwent whole-genome sequencing: long, accurate reads were obtained with Pacific Biosciences (PacBio) high-fidelity (HiFi), while Oxford Nanopore Technologies (ONT) sequencing produced ultra-long reads. Furthermore, Micro-C data were made for these three cell lines to perform haplotype phasing.

**Illumina sequencing**
All 17 orangutan lymphoblastoid cell lines were paired-end sequenced using the NovaSeq X platform (Illumina), with a read length of 150 bp and a throughput of 90 Gbp per sample, corresponding to approximately 30× coverage. Genomic DNA was extracted using the QIAamp DNA Blood Mini Kit (Qiagen Inc., Cat. no. 51104) according to the manufacturer's instructions. Macrogen Europe performed sequencing.

**Ultra-Long-ONT sequencing**
Ultra-HMW DNA was extracted from PAB16, PPY15, and PPY17 cell lines using a phenol-chloroform extraction protocol(*48*). Briefly, 3-5 × 10^7 cells were lysed in a buffer containing 10 mM Tris-Cl (pH 8.0), 0.1 M EDTA (pH 8.0), 0.5% w/v SDS, and 20 mg/mL RNase A (Qiagen, 19101) for 1 hour at 37°C. 200 µg/mL Proteinase K (Qiagen, 19131) was added, and the solution was incubated at 50°C for 2 hours. DNA was purified by two rounds of 25:24:1 phenol-chloroform-isoamyl alcohol extraction, followed by a chloroform back-extraction. DNA was precipitated with ethanol and was solubilized in EEB (ONT) at 4°C for two days. Libraries were constructed using the Ultra-Long DNA Sequencing Kit V14 (SQK-ULK114) following the manufacturer's protocol. For monarch DNA, two extractions were combined for library prep, and the final elution volume was doubled from the protocol. For phenol-chloroform-extracted DNA, approximately 40 µg was input into library prep, and the final elution volume ranged from 2× to 4× the protocol volume, depending on the amount of DNA visualized during the cleanup step. Final libraries were left at room temperature overnight to solubilize. Finally, 75 µL of library was loaded onto a primed FLO-PRO114M R10.4.1 flow cell for sequencing on the PromethION, with two nuclease washes and reloads after 24 and 48 hours of sequencing.

**PacBio HiFi sequencing**
PacBio HiFi data were generated per the manufacturer's recommendations at the University of Washington Long Reads Sequencing Center. At all steps, quantification was performed with Qubit dsDNA HS (Thermo Fisher, Q32854), measured on a DS-11 FX (Denovix), and the size distribution was checked using FEMTO Pulse (Agilent, M5330AA & FP-1002-0275). HMW DNA was extracted from cultured cells using the Monarch HMW DNA Extraction Kit (NEB, T3050L) with a lysis shaking speed of 1000 rpm. After QC, samples were pre-sheared with Megaruptor 3 DNAFluid+ (Diagenode, B06010003 & E07020001) before final shearing to a size of ~20 kbp using Megaruptor 3 Hydropores (Diagenode, E07010003) settings 28/31 or 28/30 (depending on original length distribution) and used to generate PacBio HiFi libraries via the SMRTbell Prep Kit 3.0 (PacBio, 102-182-700) using barcoded adapters (PacBio, 102-009-200). Size selection was performed with Pippin HT using a high-pass cutoff of 17 kbp (Sage Science, HTP0001 & HPE7510). Libraries were sequenced on the Revio platform on

SMRT Cells 25M with Revio Chemistry V1 (PacBio, 102-817-900) with Adaptive Loading and 30-hour movies.

##### Micro-C seq

For Micro-C, PAB16, PPY15, and PPY17 cells were sent to the University of California, Santa Cruz (UCSC), where cell preparation, crosslinking, and library preparation were performed using the Dovetail Micro-C Kit (Cantata Bio), according to the manufacturer's instructions. The prepared libraries were then sequenced at the Northwest Genomics Center on a NovaSeq X 10B platform with a 300-cycle flow cell, yielding approximately 40× genomic coverage for haplotype phasing.

##### Chip-seq

Chromatin immunoprecipitation followed by sequencing (ChIP–seq) for CENP-A and CENP-C was performed on samples PAB16, PPY15, and PPY17. Library preparation, sequencing, and data analysis were carried out according to the protocol described by Daponte et al., 2026(*65*).

##### Principal component analysis (PCA)

A PCA was carried out using a dataset comprising our 17 samples analysed together with 35 additional samples from the literature(*20, 57, 66*) and was performed both on chromosome 10 and on the neocentromeric region, including 250 kbp of its flanking regions, to provide a broader context. At the whole-chromosome level, the multi-sample VCF obtained from short-read sequencing was filtered for minor allele frequency (MAF) >0.05 using PLINK 1.9 (option *--maf*)(*67*). Additionally, markers in linkage disequilibrium (LD) were filtered with PLINK 1.9 (*--indep-pairwise 50 5 0.5*), and sites with more than 10% missingness across samples were removed. As for the neocentromeric region and its flanking regions, the VCF was filtered with a lower MAF cutoff of 1% and with LD pruning (50, 5, 0.5). PCAs were then performed with PLINK 1.9 (option *--pca*), and the results were visualized in R (2022.12.0+353) using the ggplot2 v3.5.2(*68*) package.

#### Results

##### Illumina

Illumina paired-end sequencing was performed to support polishing and variant calling. Sequencing depth ranged from 30.95× (PAB9) to 39.24× (PAB1), with total read counts ranging between ~355 million and ~805 million reads. All samples demonstrated high base quality scores, with over 97% of bases reaching Q20 and over 92% reaching Q30. Together, these datasets provide robust coverage and quality for each individual, enabling reliable *de novo* assembly, polishing, detection of structural variation, and downstream functional and comparative analyses (Table S2).

##### Oxford Nanopore Technologies (ONT)

ONT long-read sequencing was performed using PromethION flow cells, generating ultra-long reads over 100 kbp with a total coverage of 21.46× for PAB16, 21.36× for PPY15, and 57.23× for PPY17. The ONT reads had an average read N50 of 123.07 kbp, 142.91 kbp, and 93.16 kbp for PAB16, PPY15, and PPY17, respectively. Per-sample QuickStat analysis revealed read counts ranging from approximately 198,600 to 1.85 million, with 100 kbp coverage values ranging from 92.71 to 155.89 (Table S3).

##### PacBio HiFi

PacBio HiFi sequencing was performed on a PacBio Revio sequencer, yielding 7,711,174 reads for PAB16 (52.13× coverage), 7,795,612 reads for PPY15 (53.79× coverage), and 10,139,567 reads for PPY17 (59.38× coverage). The average HiFi read N50 ranged from 18.03 kbp (PPY17) to 21.89 kbp (PPY15). Sub-coverage metrics (15 kbp and 18 kbp bins) showed consistent high-quality read-length distributions across samples (Table S3).

**Micro-C and TADs analysis**

Micro-C sequencing yielded high-quality data across all samples. For PAB16, a total of 242.5 Gbp (39×) was obtained, with 223.5 Gbp (36×) bases above Q30 and an average Phred quality score of 38.32, corresponding to 92.1% of bases with Q30 or higher. Similarly, PPY15 generated 254.1 Gbp (41×) of sequence data, including 235.1 Gbp (38×) above Q30, with a mean quality score of 38.41 and 92.4% Q30 bases. Finally, PPY17 produced 218.2 Gbp (35×) total data, with 203.2 Gbp (32×) above Q30, a mean quality score of 38.56, and 93.1% of bases at Q30 or higher (Table S4).

**Chip-seq**

ChIP-seq analyses performed on PAB16, PPY15, and PPY17 LCLs using both anti–CENP-A and anti–CENP-C antibodies revealed overlapping enrichment peaks in all samples, consistent with the predicted location of the active centromere (Figs. 2 and S4.1, Supplementary Text 4). In PAB16, CENP-A and CENP-C binding regions were detected at positions 35,409,523–35,520,539 on hap1, corresponding to the hypomethylated α-sat(+)_Mono active region, and at 81,809,926–81,904,080 on hap2, within the hypomethylated α-sat(−)_Neo active region. Similarly, in PPY17, both proteins showed enrichment peaks at 82,973,593–83,175,416 on hap1, overlapping the hypomethylated α-sat(−)_Neo region, and at 35,178,468–35,290,552 on hap2, within the hypomethylated α-sat(+)_HOR active region. Finally, in PPY15, CENP-A and CENP-C peaks were observed on both haplotypes within hypomethylated regions of the α-sat(-)_Neo active domain, mapping to 82,973,593–83,175,416 on hap1 and to 83,014,811–83,090,006 on hap2.

**PCA of the orangutan cohort**

The population structure of our 17 samples was analyzed alongside 35 additional samples from the literature to provide a broader context for genetic variation (Table S5). The PCA based on the entire chromosome 10 (Fig. 1C) revealed a clear separation between PAB and PPY. The first principal component (PC1, 35.35% of the variance) distinguishes the two species, whereas PC2 (8.47% of the variance) highlights within-species structure. The individual PPY19 occupies an intermediate position between the two main clusters, consistent with its status as a Borneo × Sumatra hybrid. The PCA restricted to the neocentromeric region (Fig. 1D) revealed a similar pattern of species differentiation, although the separation between clusters is slightly less pronounced than for the whole chromosome. Of note, within the PAB cluster, an additional internal separation (along the PC2) can be observed, distinguishing heterozygotes carrying the neocentromere from homozygous WT individuals. Moreover, among the PAB samples from the literature, one sample is positioned separately from the others, which could hypothetically represent a NEO/NEO genotype; however, this cannot be confirmed because the corresponding cell line is unavailable. This distinction, however, is detectable only among our sequenced samples, as the status of samples from the literature is otherwise unavailable. In contrast, no such internal separation is evident within the PPY cluster.

243

#### 3. Genome assemblies

##### Methods

The genome assemblies of the three representative samples (PAB16, PPY15, and PPY17) were generated using Verkko (v2.2.1;(*22*)), a widely adopted assembler known for producing high-quality, phased genomes. Verkko first constructs a backbone using the highly accurate HiFi reads (*--hifi*), then incorporates ultra-long ONT sequences (*--nano*) to improve resolution and scaffolding, ensuring excellent assembly performance even in repetitive regions. By integrating Micro-C data (*--hic1 R1.fofn --hic2 R2.fofn*), Verkko further enables haplotype phasing at the chromosomal scale, ultimately producing a fully diploid assembly. In addition, the parameter *--screen-human-contaminants* was used to remove all sequences matching the typical human contaminants from the main assembly output.

```
verkko -d VerkkoWithHic/ --hifi $( cat HiFi.fofn ) --nano $( cat ONT.fofn ) --hic1
$( cat /HiC_R1.fofn ) --hic2 $( cat HiC_R2.fofn ) --screen-human-contaminants
```

**Genome assembly assessment**

All assemblies were annotated for potential misassemblies using Flagger (v0.3.3; https://github.com/mobinasri/flagger) and NucFlag (https://github.com/logsdon-lab/NucFlag). Assembly quality was further assessed by computing quality values (QVs) with Merqury (v1.3;(*69*)), and gene completeness was evaluated using Compleasm (v0.2.2;(*70*)). Alignments were produced using the *assembly_eval* pipeline (https://github.com/EichlerLab/assembly_eval) with default parameters. The *type_map* setting was specified according to the downstream tool: Winnowmap (v2.03) for Flagger, and pbmm2 (v1.10.0; https://github.com/PacificBiosciences/pbmm2) for NucFlag. Flagger was executed using the *flagger_end_to_end_no_variant_calling_no_ref.wdl* workflow provided in the repository, taking as input HiFi reads mapped to each assembly. NucFlag was run on the BAM files of PacBio HiFi reads aligned to each assembly, and, when available, a BED file specifying genomic regions of interest was provided. NucFlag generated a BED file of predicted misassemblies and a coverage plot showing the first- and second-most frequent bases, along with flagged misassembled regions. QV estimation with Merqury used a hybrid database of 21-mers collected from Illumina and HiFi reads via Meryl (v1.3; https://github.com/marbl/merqury/wiki/1.-Prepare-meryl-dbs). Compleasm was run using Illumina reads only to assess gene completeness for each assembly.

**Chromosome 10 assignment and contig orientation assessment**

To assess the coverage of chromosome 10 in the genome assemblies and to identify the contig(s) mapping to this chromosome, we aligned each assembly to the appropriate T2T reference using minimap2 (v2.28; parameters: *-cx asm20 -t 24*;(*71*)). Specifically, assemblies from *Pongo pygmaeus* (PPY) samples were aligned to the *mPomPyg2_v2.0_pri* reference, while the PAB16 sample (*Pongo abelii*) was aligned to *mPonAbe1_v2.0_pri*.

```
minimap2 -cx asm20 -t 24 -a Reference.genome.fa /assembly.haplotype1.fasta >
sample_hap1_VS_RefT2T.sam
for sam in *.sam; do samtools view -Sb "$sam" > "${sam%.sam}.bam"; done
```

The resulting alignments were filtered using SAMtools (v1.21;(*72*)) *view* to extract reads mapping to chromosome 10 (NC_072383.2 for PPY and NC_071995.2 for PAB). The resulting *.bam* and *.bai* (obtained with *samtools index*) files were loaded into Integrative Genomics

Viewer (IGV v2.18.4;(*70*)) to visualize the aligned contigs and evaluate their orientation relative to the reference chromosome 10.

```
samtools view -bh sample_hap1_VS_RefT2T.bam "NC_071995.2" > sample_hap1_VS_RefT2T_chr10.bam
samtools index sample_hap1_VS_RefT2T_chr10.bam
```

When contigs mapping to chromosome 10 were found to have a negative orientation, they were reverse-complemented using seqtk (v1.4; https://github.com/lh3/seqtk). First, the relevant contig was extracted from the assembly using *seqtk subseq*, then reverse-complemented with *seqtk seq -r*. During this process, the name of the flipped contig was modified by appending the suffix _v1. To reconstruct the corrected assembly, all contigs except the one to be flipped were extracted from the original assembly using *samtools faidx -r*, and the reverse-complemented contig was then added in place of the original.

```
echo -e contig2flip_name > contig2flip.bed
seqtk subseq assembly.haplotype1.fasta contig2flip.bed > contig2flip.fasta
seqtk seq -r contig2flip.fasta | sed 's/^>.*$/>v1/' > contigFlipped.fasta
awk '{print $1}' assembly.haplotype1.fasta.fai > contigs2keep.txt
grep -v "contig2flip" contigs2keep.txt > contigs2keep_minus_contig2flip.txt
samtools faidx -r contigs2keep_minus_contig2flip.txt assembly.haplotype1.fasta >
assembly.haplotype1_v1.fasta
cat contigFlipped.fasta >> assembly.haplotype1_v1.fasta
samtools faidx assembly.haplotype1_v1.fasta
```

#### Results

All assemblies generated with Verkko showed high contiguity and base-level accuracy. Haploid N50 values ranged from 137.7 Mbp (PPY15_hap1) to 146.0 Mbp (PAB16_hap2), while estimated QVs spanned from 55.6 to 59.1, indicating very low base-level error rates (Table S6). Assessment of assembly completeness and redundancy revealed that single-copy sequences comprised the majority of each assembly. The proportions of collapsed (Col), duplicated (Dup), and erroneous (Err) bases were minimal across all haplotypes, with most bases assigned to the haplotype-specific sequence (Hap), accounting for over 93% to 99% of each assembly. For example, in PPY17, 99.6% of bases were correctly assigned to the haplotypes, with only 0.12% collapsed, 0.23% duplicated, and 0.07% erroneous (Table S7). Gene completeness analysis using Compleasm(*73*) (Table S8) indicated that more than 98% of single-copy genes were complete in all haplotypes, with duplicated genes accounting for less than 0.7%. Fragmented genes were rare (<0.4%), and missing genes ranged from 0.13% (PPY15_hap2) to 3.48% (PAB16_hap1), consistent with highly complete assemblies.

All primary contigs were unequivocally assigned and correctly oriented to chromosome 10, as indicated by the "_v1" suffix, denoting finalized orientation. Both haplotypes of each assembly consistently anchor to chromosome 10 reference (NC_071995.2; ~132.28 Mbp), with notable length differences ranging from ~130.98 Mbp (PAB16_hap2) to ~135.71 Mbp (PPY17_hap2) (Table S9). Despite this variation, each haplotype achieves near-complete coverage of chromosome 10, supporting near-T2T assemblies, with only terminal telomeric regions unresolved.

#### 4. Centromere analyses

##### Methods

**Repeat annotation and detection of α-sat arrays**
Repetitive elements were annotated using RepeatMasker (v4.1.6;(*23*)) with parameters optimized for primate genomes (*-species Primates*). Each assembly was screened in sensitive mode (*-s*) using the NCBI search engine (*-e ncbi*), producing soft-masked output (*-xsmall*) for downstream analyses. The following command was used:

```
RepeatMasker -s -e ncbi -xsmall -species Primates -pa 16 assembly.fasta
```

The resulting annotations were subsequently filtered to retain only entries classified as ALR/alpha, corresponding to canonical α-sat monomers. These filtered intervals were then merged and manually inspected to define continuous α-sat blocks, which were used to delineate putative centromeric coordinates for each contig and haplotype.

**Methylation and CDR definition**
All ONT sample data were base called with Guppy v6.3.7 (or newer) or Dorado v0.7.3 (or newer) to generate BAM files containing methylation tags for each sequencing read. Methyl-reads were extracted from the BAM files and aligned to the corresponding sample assembly using winnowmap v2.03(*74*). The resulting BAM files were used as input for CDR-Finder (v1.0.1)(*28*) to generate the CDR calls and associated plots.

**NucFlag**
Potential misassembled regions were identified using NucFlag (v0.3.3;(*75*)). NucFlag, based on NucFreq, detects local deviations in nucleotide frequencies that may indicate assembly errors. Nucleotide frequency plots and misassembly region files were generated from HiFi read alignments to the species-specific reference genome using the software's default parameters.

**ModDotPlot**
To assess internal sequence organization, dot plots were generated individually for each centromeric and neocentromeric region using ModDotPlot (v0.8.4;(*25*)), with a minimum identity threshold of 80% (*--identity 80*). Sequences were extracted from the orangutan genome assemblies using *samtools faidx*, spanning the annotated centromere or neocentromere and including 100 kbp of flanking sequence on either side to provide local genomic context.

```
samtools faidx assembly.fasta chr10:(start-100bp)-(end+100bp) > selectedRegion.fasta
moddotplot static -f selectedRegion.fasta --identity 80
```

These extracted regions were used as input for dot plot generation to assess the internal repeat structure and sequence organization and to visualize self-similarity within each region.

**Pairwise sequence identity heatmaps and SVbyEye plot**
To identify homologous regions across different haplotypes, particularly within centromeric and neocentromeric loci, and to visually inspect potential structural rearrangements/diversity, a specific targeted genomic region was first extracted from the reference assembly using *samtools faidx* and aligned against a set of target assemblies using minimap2 (with parameters *-x asm20 --eqx -c -t 50* and *-secondary=no*). For each target, alignments longer than 200 kbp were retained, and the region with the highest cumulative alignment length was identified and

extracted based on those coordinates. The resulting sequences were then pairwise-aligned in a predefined order using minimap2, and the *.paf* files were merged into a single alignment file. This combined file was subsequently processed with the *rb break-paf* tool (run with the *-m 100* parameter) to split the alignments into blocks of at least 100 bp, thereby removing shorter, potentially spurious alignments. The processed alignment data were then imported into R and visualized using the SVbyEye R package (https://github.com/daewoooo/SVbyEye), enabling clear visualization of structural differences across assemblies. In addition to structural variant visualization, this approach proved useful for retrieving and comparing syntenic region coordinates across haplotypes, particularly within complex centromeric and neocentromeric loci, by analyzing and interpreting the structure and continuity of alignments in the resulting PAF files.

**CENdetectHOR**

To characterize the higher-order repeat (HOR) structure in the three samples (PAB16, PPY15, PPY17), we ran the CENdetectHOR pipeline(*24*), providing as input the complete sequences of both haplotypes of chromosome 10, along with the human α-sat consensus sequence (GenBank: X07685.1). The only parameter modified from the default settings was the k-mer size, which we set to 9 bp. The resulting trees were then screened using CENdetectHOR's graphical interface, PhylotreeGUI, to identify potential HORs in each sequence. All monomers corresponding to each family, whether forming monomeric arrays or HORs, were extracted and aligned using MAFFT(*76*). Consensus sequences for each family were generated with EMBOSS cons(*77*). A Levenshtein distance matrix was then computed by pairwise comparison of all consensus sequences, representing each identified monomeric family. Identical families were merged, thereby removing redundant HORs. Finally, HOR variants were defined by clustering together all HORs that shared more than half of their constituent unique monomeric families.
For phylogenetic analysis, five sequences were randomly selected from each family, whether forming monomeric arrays or HORs, and analyzed together with the human α-satellite consensus sequence (GenBank: X07685.1). Sequences were aligned using the MUSCLE algorithm for non-coding DNA, and a phylogenetic tree was constructed in MEGA (*78*) using the Maximum Likelihood method with the General Time Reversible model. The bootstrap consensus tree inferred from 1000 replicates was used to represent the evolutionary history of the taxa analyzed, and branches supported in fewer than 50% of bootstrap replicates were collapsed.

#### Results

α-sat were identified in all chromosomes 10 of the analyzed individuals, showing marked variation in length between haplotypes (Table S10). In PAB16, hap1 contains 118,652 bp (35.41–35.53 Mbp) and hap2 214,978 bp (35.30–35.53 Mbp). In PPY15, hap1 has 201,203 bp (35.38–35.58 Mbp) and hap2 242,496 bp (35.30–35.53 Mbp) (Fig. S2). PPY17_hap1 harbors a single array of 191,482 bp (35.34–35.53 Mbp), whereas hap2 uniquely contains two separate α-sat arrays, one spanning 2,116,114 bp (34.3–36.4 Mbp) and a second of 211,639 bp (38.8–39.0 Mbp), separated by ~2.4 Mbp (Fig. 2; Fig. S2). Notably, arrays smaller than ~250 kbp are not composed of continuous α-sat sequences; instead, particularly at the array start, α-sat monomers are interrupted by a fairly repetitive pattern of LINE elements, suggesting a mosaic composition of satellite and interspersed repeats (Fig. 2; Fig. S2).

To investigate epigenetic activity, CDRfinder was used to profile methylation across α-sat regions. Hypomethylated intervals, indicative of potential centromeric activity, were detected only in PAB16_hap1 (1,200,000 bp, 35.41–35.53 Mbp) and in the longer α-sat(+)_HOR array of PPY17_hap2 (1,161,114 bp, 34.25–36.37 Mbp). In putative neocentromeric regions, homologous to the sequence described by Tolomeo et al.(*19*) and identified via pairwise sequence identity heatmaps and SVbyEye plot, hypomethylation was observed in PAB16_hap2 (120,000 bp, 81.80–81.92 Mbp), PPY15_hap1 (100,000 bp, 82.96–83.06 Mbp), PPY15_hap2 (100,000 bp, 83.01–83.11 Mbp), and PPY17_hap1 (120,000 bp, 82.95–83.06 Mbp) (Fig. 2, Fig. S2, Table S11).

Among all the analyzed sequences, CENdetectHOR revealed the presence of HOR structures in a ~2Mb region in hap2 of PPY17 solely (α-sat(+)_HOR region; PPY17_hap2:34,250,000-36,400,000), as depicted in Fig. 2. This region is predominantly composed of a 12-mer organization (C10_hap2_M12.2), spanning 353,369 bp, together with its variants, which collectively account for ~1.1 Mb of the region, as detailed in Fig. S3A, Table S12-S13. In addition to the 12-mer, three other HORs were detected within this interval: C10_hap2_M9 (9-mer; 4,955 bp), C10_hap2_M10 (10-mer; 6,834 bp), and C10_hap2_M33 (33-mer; 21,504 bp). Together, these account for approximately 33 kb of the HOR array. The remaining portion of the region consists of monomeric α-sat arrays composed of the C10_hap2_F1 family (Table S13).

Phylogenetic clustering based on five representative sequences per family, including both monomeric and HOR-associated families (Fig. S3B), reveals a clear separation into two major clades. The HOR-associated clade (depicted in red) comprises all families from the single haplotype exhibiting HOR organization (PPY17_hap2), including both HOR-associated and monomeric families. In contrast, the second clade contains families from the remaining haplotypes, which are exclusively organized as monomeric arrays.

ModDotPlot analysis (Fig. 2; Fig. S2) revealed that most α-sat(+)_Mono regions lack the typical long-range sequence identity of the centromere, independent of their activity state. This reduced similarity is consistent across the PAB16, PPY15, and PPY17_hap1 haplotypes. Only PPY17_hap2, containing an α-sat(+)_HOR array, exhibits a structure partially reminiscent of canonical centromeres. In contrast, neocentromeric intervals exhibit an almost complete absence of sequence identity, with only sparse, highly diverged structures present, generally showing <85% similarity.

#### 5. PPY and PAB alphoid characterization with PCR and Immuno-FISH

##### Methods

**FISH experiments with PCR amplified probes**

Primer pairs were designed with Primer3Plus (v3.3.0) using default settings, with a target amplicon size of 100-200 bp; primers were synthesized and desalted by Thermo Fisher Scientific. Primer sequences and the genomic regions on which primer design for each experiment was based are provided in Table S14.

In experiment PPY_α-sat(+)_HOR, we designed a primer pair targeting the hypomethylated region of the α-sat centromeric domain on chromosome 10 PPY17_hap2 (α-sat(+)_HOR_active in Fig. 2). BLASTn(*79*), with default settings, was used to align the expected amplicon sequence to the seven chr10 α-sat sequences characterized in this study (PAB16_hap1_α-sat(+)_Mono_active; PAB16_hap2_α-sat(+)_Mono_inactive; PPY15_hap1_α-sat(+)_Mono_inactive; PPY15_hap2_α-sat(+)_Mono_inactive; PPY17_hap1_α-sat(+)_Mono_inactive; PPY17_hap2_α-sat(+)_HOR_active; and PPY17_hap2_α-sat(+)_Mono inactive). PCR was performed on PPY17 genomic DNA to generate a probe to screen all PAB and PPY specimens and to investigate the presence/absence of the α-sat with HOR-based organization on chromosome 10.

In experiments PAB_chr1_α-sat(+)_HOR, PAB_chr2_α-sat(+)_HOR, and PAB_chr4_α-sat(+)_HOR, we selected primer pairs targeting the hypomethylated region of the active centromeric domains on chromosomes 1, 2, and 4, of the NHGRI_mPonAbe1-v2.1_pri reference genome, intending to design a set of probes that covers the diversity of centromeric organization across all chromosomes in the PAB species, revealing any differences with chromosome 10 α-sat region.

Each PCR was performed in a 25 µL reaction containing 5 ng of genomic DNA, 0.25 µM of each primer, and 1X PCR Master Mix (Thermo Fisher Scientific). Negative control reactions (also called 'white'), lacking template DNA, were included to monitor for contamination. Amplification was performed using the manufacturer-recommended thermal cycling conditions: initial denaturation at 95°C for 3 min; 35 cycles of denaturation at 95°C for 30 s, annealing at 60°C for 30 s, and extension at 72°C for 1 min; and a final extension at 72°C for 7 min. PCR products were separated by electrophoresis on 1% agarose gel. Gel visualization was performed using the Bio-Rad ChemiDoc imaging system. Each PCR product was then labeled as follows: 1 µL of the amplified product was taken from the PCR reaction mixture and used as a template for incorporation of fluorescently labeled dUTP, enabling it to be used as a probe in FISH experiments. The PCR products obtained from PPY_α-sat(+)_HOR and PAB_chr1_α-sat(+)_HOR were labeled with Cy3-dUTP, whereas those from PAB_chr2_α-sat(+)_HOR and PAB_chr4_α-sat(+)_HOR were labeled with Fluorescein-dUTP and Cy5-dUTP, respectively. Each labeling reaction was performed in a total volume of 25 µL, using 1X PCR buffer (Thermo Fisher Scientific), 2 mM $MgCl_2$ (Thermo Fisher Scientific), 0.2 µM of each primer, 0.2 mM of each unlabeled nucleotide, 0.1 mM of labeled dUTP, 0.4% BSA, and 0.06 U/µL Taq DNA polymerase (Thermo Fisher Scientific), and incubated following a single-step temperature program on the thermocycler: initial denaturation at 95°C for 30 s; 1 cycle of denaturation at 95°C for 30 s, annealing at 60°C for 1 min, extension at 72°C for 1 min; and final extension at 72°C for 10 min. In addition, two control reactions were performed for each labeling: one with the PCR negative control (reaction mix without genomic DNA template) and one with the genomic DNA template (diluted 1:25) to confirm the absence of nonspecific signals. Each probe was precipitated, denatured, and hybridized, without the Human Cot-1 DNA, due to the highly repetitive nature of the α-sat sequences used as probes. Post-hybridization washes (under high-stringency conditions), fluorescence detection, and image acquisition were also performed as described in the main text.

**Immuno-FISH with CENP-C monoclonal antibodies**

Immunofluorescence was performed to detect the CENP-C protein on the metaphase spreads prepared from AG05252 (PPY_T2T) and PAB16 cell lines. Metaphase spreads were dropped onto glass slides and aged for 72 h at 37°C. An immunofluorescence assay was performed on metaphase spreads using mouse CENP-C monoclonal antibodies, as previously described with minor modifications(*80*). Slides were rehydrated in 1X PBS-azide (10 mM NaPO4, pH 7.4, 0.15 M NaCl, 1 mM EGTA, and 0.01% NaN3) for 8 min at RT and washed three times (each for 2 min) in 1X TEEN, 0.5% TritonX, and 0.1% BSA buffer. Slides were incubated with 100 µL/slide of mouse anti-CENP-C monoclonal antibody (Abcam) at 0.001 µg/µL for 2 h at 37°C. After incubation, slides were washed three times (2, 5, and 3 min, respectively) in 1X KB buffer (10 mM Tris-HCl, pH 7.7, 0.15 M NaCl, and 0.1% BSA) and incubated with 100 µL/slide goat anti-mouse IgG secondary antibody conjugated to FITC (Abcam, diluted 1:100) for 45 min at 37°C. Slides were washed in 1X KB buffer for 2 minutes at RT; pre-fixed in a 4% paraformaldehyde solution prepared in 1X KB for 45 minutes at RT; washed in distilled water for 10 minutes at RT; fixed in Methanol:Acetic acid (3:1) solution and air-dried before proceeding with FISH, performed with the 101 bp designed probe (experiment PPY_α-sat(+)_HOR), labeled with dUTP-Cy3. The denaturation step prior to hybridization was performed for 8 min at 70°C.

#### Results

**Detection of α-sat(+)_HOR arrays across specimens**

BLASTn(*79*) analyses of the expected PCR product from experiment PPY_α-sat(+)_HOR (Table S14) showed lower sequence identity with the other six α-sat sequences used as queries than with the centromeric region from which the amplicon was derived (PPY17_hap2_α-sat(+)_HOR) (Table S15). The resulting PCR product yielded a strong band on 1% agarose gel at approximately 100 bp, consistent with the expected 101 bp amplicon (Fig. S4). Faint secondary bands at ~500 bp and ~1000 bp were also observed, likely associated with the HOR organization of these sequences.

The 101 bp PCR product (PPY_α-sat(+)_HOR) in co-hybridization with the BAC probe RP11-107L10 was used to screen all the cohort samples. Among the PPY specimens, the presence of a hybridization signal, likely representing the long stretch of α-sat, was detected only in one chr10 homolog of PPY12, PPY17, PPY19, and AG05252 (Fig. S5; Table S16). No probe hybridization was detected on the canonical centromeres of the other chromosome 10 homologs. No hybridization signals were detected on the canonical centromeric region of chromosome 10 in all the PAB individuals (Fig. S6; Table S16).

**Assessment of the HOR structure on the PAB chromosome 10**

PAB_chr1_α-sat(+)_HOR, PAB_chr2_α-sat(+)_HOR, and PAB_chr4_α-sat(+)_HOR PCR performed on the PAB16 lymphoblastoid cell line using chromosome-specific α-sat primers, produced main amplicons of the expected sizes (106 bp for chr1, 101 bp for chr2, 190 bp for chr4). Faint secondary bands were also observed: for chr1, multiple bands spaced ~100–200 bp, and for chr2, an additional band at ~500 bp, possibly reflecting HOR organization. For chr4, only a strong, specific band was present (Fig. S7).

To capture a broad range of variability in centromeric organization across PAB chromosomes, a three-color FISH experiment was performed on metaphase spreads from each PAB individual in the cohort, using the three amplicons as probes (Fig. S8). A strong hybridization signal was detected from all centromeres, except for the canonical centromeric domain of chromosome 10, which was never recognized by any of the three probes in PAB individuals, highlighting the unique organization of this region.

**Immuno-FISH with CENP-C monoclonal antibodies**

Immuno-FISH was performed on AG05252 (Fig. S9A) to co-visualize CENP-C binding sites and hybridization of the 101 bp probe (PCR product PPY_αSat(+)_HOR) on chromosome 10, comparing the distribution of the centromeric protein along the two homologs with the organization of the α-sat array in the centromeric domain. A clear CENP-C signal co-localizing with the α-sat probe was detected on the HOR-based α-sat active centromeric domain on chromosome 10. On the homolog lacking the probe signal, CENP-C binding was still observed at the corresponding monomeric α-sat active site, with a very weak signal that was also detectable further downstream, in the region where a neocentromere is expected to form, in approximately 17% of metaphases analyzed. The immuno-FISH experiment was also performed on PAB16 (Fig. S9B), which is heterozygous for the neocentromere on chromosome 10. CENP-C signals were detected on the PAB16_hap1_α-sat(+)_Mono_active centromeric domain and on PAB16_hap2_α-sat(-)_Neo_active neocentromeric region. No FISH signal was detected on either chromosome 10, as previously shown.

599

#### 6. Analysis of 3D genome organization

##### Methods

Micro-C reads for PAB16, PPY15, and PPY17 were aligned to the corresponding assemblies using BWA-MEM 2(*81*) with the -SP5M parameters, recommended for HiC analysis. For haplotype-specific analyses, the alignments were phased and split by haplotype using Duet version 0.6(*82*) with default parameters except the inclusion of the parameter --*include_all_ctgs* to enable processing chromosomes with nonstandard names. Duet enables haplotype phasing and splitting by combining Clair3(*83*) and WhatsHap(*84*). Aligned reads were then saved in separate haplotype files using SAMtools view(*85*). Identification of ligation junctions was performed using pairtools(*86*). Contact matrices were generated using the cooler cload pairs command from the cooltools suite(*87*). Topologically associating domains were identified as minima in insulation scores, computed with the insulation command in cooltools at a resolution of 100 kbp. Chromosomal regions were assigned to A and B compartments based on eigenvectors of cis-interactions using the eigs-cis command in cooltools, at a resolution of 1 Mbp. The sign of the A and B compartments was assigned based on chromosome-wide gene density.

##### Results

A total of 1.6, 1.7, and 1.4 billion read pairs were obtained by sequencing Micro-C libraries for PAB16, PPY15, and PPY17, respectively. The number of unique intra and interchromosomal contacts, as identified from the primary alignment of Micro-C reads, is summarized in Table S17.

The global 3D organization of chromosome 10 is highly comparable between the three cell samples and follows patterns as previously observed in human cells (Fig. S10). This includes a preference for nearby interactions, as evidenced by the higher number of contacts near the diagonal. Moreover, we observe a reproducible "checkerboard" pattern of interactions (particularly noticeable around 50 Mbp), which is consistent with an A/B compartment organization, whereby multi-Megabase active and gene-dense A compartments preferentially interact with other A compartments, and inactive and gene-poor B compartments with other B compartments(*88*). Centromeric (Fig. S11) and neocentromeric (Fig. S12) regions are gene-poor and reside in the B compartment in all three samples (blue regions in First Eigenvector track). The main difference between samples was observed at heterozygous positions: PPY samples show a sharp decrease in heterozygosity in CEN and neocentromere, whereas this is not observed in PAB16.

Zoom-ins on centromere and neocentromere reveal marked differences in chromatin organization for both structures (Figs. S11 and S12). The centromere, in all three samples, is localized at the centre of a ~3 Mbp Topologically Associated Domain (TAD) (Fig. S11, with red arrows highlighting TAD boundaries in the insulation score track). Within the TAD, a noticeably "sub-TAD" can be observed that exactly overlaps the centromere. A further zoom-in confirms this precise overlap, showing that the sub-TAD overlaps the domain of DNA methylation that surrounds the location of the centromere (Fig. S13). Local peaks in DNA methylation signal form the boundaries of this sub-TAD. The centromeric domain is therefore characterized by strong intra-domain contacts, yet contacts within the surrounding TAD remain enriched as well. The TAD is located within a larger B compartment, characterized by a similar

First Eigenvector pattern in all cell types, with its left boundary serving as the separation from a neighboring, gene-dense TAD that forms a smaller A compartment (Fig. S11).

The annotated neocentromere, in all cell types, exactly overlaps a wide TAD boundary, as identified by the extended minimum in the insulation score track (Fig. S12; red bars in the insulation score track). This TAD boundary is localized within a large B compartment that globally follows a similar First Eigenvector pattern across all cell types. Yet, a minor difference can be observed: the minimum signal is noticeably lower in PPY15 (where the neocentromere is active on both alleles) as compared to the other cell types. Moreover, upon detailed inspection of the PPY15 contact matrix, two "stripes" of depleted signal are observed that emanate from a small domain with increased signal (Fig. S12; PPY15 column: red and blue arrows). A further zoom-in confirms the presence of this domain, which contains the hypomethylated region of the CENP-A signal (Fig. S13). The active neocentromere in PPY15 has thus adopted a specialized 3D chromatin organization, whereby the small structure reduces its contacts with neighboring TADs.

While a similar domain may be observed in PAB16 and PPY17 (where the neocentromere is active on one allele), here a more defined TAD is visible on the right as well (Fig. S12; PAB16 and PPY17 columns: red and blue arrows and Fig. S13). We speculate that the active and inactive neocentromere in PAB16 and PPY17 reside in different chromatin environments:

- The active neocentromere, like in PPY15, occupies the small hypomethylated and CENP-A carrying domain.
- The inactive neocentromere resides at the boundary of a larger TAD.

#### 7. Repeat annotation, LINE, and segmental duplication (SD) analyses

##### Methods

**Repeat annotation on chromosome 10 and the evolutionary analysis of segmental duplication**

Repetitive sequences in the genome assemblies were initially masked using three tools: TRF (v4.1.0;(*89*)) with the *command trf [asm.fa] 2 7 7 80 10 50 2000 -l 30 -h -ngs*, RepeatMasker (v4.1.5;(*90*)) with the command RepeatMasker *-s -e ncbi -xsmall -species human [asm.fa]*, and WindowMasker (v2.2.22;(*91*)), executed in two steps: *windowmasker -mk_counts -mem 16384 -smem 2048 -infmt fasta -sformat obinary -in [asm.fa] -out [asm.count]* followed by *windowmasker -infmt fasta -ustat [asm.count] -dust T -outfmt interval -in [asm.fa] -out [asm.interval]*. Genomic sequences were soft-masked using the combined BED files generated from all three repeat annotations. SDs were then identified using SEDEF (v1.1;(*92*)) on the repeat-soft-masked genomes. Detected SDs were filtered to retain only those with pairwise sequence identity greater than 90%, lengths exceeding 1 kbp, and satellite content below 70%.

**Phylogenetic tree analysis of a 250 kbp SD sequence**

Phylogenetic analysis was conducted using the maximum likelihood method implemented in IQ-TREE (v2.3.6;(*93*)). A multiple sequence alignment (MSA) of the 250 kbp duplicated region was generated using MAFFT (v7;(*76*)). Based on this MSA, a phylogenetic tree of the SD was constructed using the TVM+F+I+R4 substitution model, which was selected as the best-fit model according to the Bayesian Information Criterion. Each tree was generated with 1,000 ultrafast bootstrap replicates (-B 1000). The resulting trees were then time-calibrated using a divergence time of 28.8 Mya to macaque (T2T-MFA;(*94*), as previously reported by the time tree database(*95*), while also incorporating IQ-TREE parameters *--date-tip 0 --date-ci 100*. To infer a more precise timing of the NAHR between 250 kbp SDs, we first checked that the recombination breakpoint falls within the duplication by performing alignment between two copies of the duplicated sequence and the recombined single copy. This was done by assessing the reduction of sequence identity (<99%) of the synteny alignment block. Then, a 50 kbp sequence was subset from the left and right of the breakpoint for phylogenetic tree reconstruction. The time of the recombination event was estimated from the coalescent time between the recombined copies.

**Comparative analyses of orangutan SD sequences across primates**

BLAST(*79*) analyses were performed to determine whether the Bornean orangutan SD is species-specific or also occurs in other primate species. SD1 (chr10_hap2_hsa12: 32858690-33111201) and SD2 (chr10_hap2_hsa12: 36152001-36404139) sequences (from the alternative haplotype chromosome 10 of mPonPyg2-v2.1 reference genome) were aligned against all available primates T2T reference genomes: NHGRI_mPanTro3-v2.1_pri (GCF_028858775.2), NHGRI_mPanPan1-v2.1_pri (GCF_029289425.2), NHGRI_mGorGor1-v2.1_pri (GCF_029281585.2), NHGRI_mPonAbe1-v2.1_pri (GCF_028885655.2), T2T-CHM13v2.0 (GCF_009914755.1), and T2T-MMU8v2.0 (GCF_049350105.2). The Megablast algorithm was used with default parameters, and for each alignment, both bitscore and query coverage were used as key metrics to identify the best hits. This first approach allowed us to identify the chromosome that carries these sequences in all other species. A complementary analysis with minimap2 defined the precise genomic coordinates of the sequences orthologous to the analyzed SDs. The region that spans from SD1 to SD2 on chr10_hap2_hsa12: 32858690-36404139 (mPonPyg2-v2.1 reference genome) was aligned against PAB (NC_071995.2) chromosome 10 sequence, with the command *minimap2*

*-x asm5 -c*; and against the PTR (NC_072408.2), PPA (NC_073259.2), and GGO (NC_073234.2) chromosomes 10, the HSA chromosome 12 (NC_060936.1) and the MMU chromosome 11 (NC_133416.1), with the command *minimap2 -x asm20 -c*, suited for the comparison of most divergent genomes. Only alignments longer than 30 kbp were considered. To ensure the uniqueness of the identified regions within their respective genomes, the Megablast algorithm with default options was used to align each minimap-aligned sequence to the corresponding full genome.

##### LINE content within the α-sat region of chromosome 10

RepeatMasker tracks available for the T2T genome assemblies(*21*) for each species were parsed to retain only entries annotated as α-sat (ALR/Alpha) or LINE1 elements. To focus on potentially recent and structurally relevant insertions, only LINE1 annotations with a length greater than 5 kbp were retained for downstream analysis. This threshold was chosen to exclude highly fragmented elements and to enrich for relatively intact LINE copies that may correspond to recent or moderately recent insertions. Only the filtered LINEs that interdigitated with the α-sat were then considered for comparison of their content across the various species, taking into account their classes of belonging and their dating(*31*).

##### Visualization of homologous α-sat regions of chromosome 10

The homologous regions of PPY_hap2 (32-39 Mbp) sequence in great apes (primary haplotypes of chimpanzee, human, gorilla, Bornean and Sumatran orangutans;(*21, 96*)) and macaque (T2T-MFA;(*94*)) were identified using minimap2 (v2.28). Progressive alignments were generated with minimap2 again with the extracted sequences, allowing for secondary alignment with the following parameters "*-x asm20 -c --eqx*". The visualization was made using the SVbyEye R package (https://github.com/daewoooo/SVbyEye;(*97*)).

#### Results

##### Comparative analyses of orangutan SD sequences across primates

Hits obtained from every Megablast alignment performed using SD1 and SD2 queries were summarized for each targeted chromosome within the genome, and the best match was identified based on a combination of maximum bitscore and query coverage (Table S18). These results show that the orthologous sequences of the analyzed SDs consistently map to the chromosome homologous to the orangutan chromosome 10 across all tested species (chimpanzee, bonobo, gorilla, Sumatran orangutan, human, and macaque). Considering only hits longer than 30 kbp, minimap2 results (Table S19) allowed us the precise identification of the genomic coordinates of the domain orthologous to both the Bornean orangutan analyzed SDs: NC_072408.2: 63267079-63493351 in PTR; NC_073259.2: 64531336-64763832 in PPA; the sequence NC_073234.2: 68993633-69226910 in GGO; the sequence NC_071995.2: 33119293-33371437 in PAB; the sequence NC_060936.1: 31872704-32096064 in HSA; and the sequence NC_133416.1: 34265535-34393967 in MMU; thereby confirming that both SDs map to the same target genomic domain.

These regions identified as orthologous to the Bornean orangutan SD sequence were then aligned against each T2T reference genome (Table S20), confirming the sequence's uniqueness within each analyzed genome: the top match aligns perfectly with the query, while the remaining hits represent spurious alignments.

The alignment of neighboring sequences (PPY17_hap2) confirms the SDs (overlapping with *BICD1*) are located in syntenic sequences across the apes and a macaque (Fig. S14). Examining the sequences, we also find that there are two stretches of long and short α-sat arrays in PPY17_hap2, and that the short array corresponds to homologous locations with respect to other apes, suggesting the longer α-sat array is orangutan-specific. We find that the position of the centromere is different in the macaque located further downstream with respect to both short and long stretches of α-sat of PPY17_hap2.

**LINE content within the α-sat region of chromosome 10**

Across all assemblies examined, the α-sat region of chromosome 10 contained multiple LINE1 insertions longer than 5 kbp, indicating that the centromeric array is interspersed with transposable element sequences rather than being composed exclusively of α-sat DNA. The complete list of filtered RepeatMasker annotations is provided in Table S21, while a summary of the total number of LINE1 elements longer than 5 kbp detected in each species is reported in Tables S22-S26. Classification of the LINE elements revealed several relatively recent L1 subfamilies, including L1P1, L1PA2, L1PA3, and L1PA4. In human (HSA) and chimpanzee (PTR), the α-sat region contained LINE1 elements belonging to multiple subfamilies, including L1P1 together with several L1PA elements. Bonobo (PPA) showed a higher number of long LINE1 insertions overall, with elements assigned to L1PA2, L1PA3, and L1PA4. Gorilla (GGO) displayed a more restricted composition, with only L1PA3 and L1PA4 elements detected within the α-sat block. In contrast, in the PPY haplotype carrying both HOR and monomeric α-sat arrays, all LINE1 elements longer than 5 kbp identified within the α-sat region belonged to the L1PA3 subfamily. Because L1 subfamilies have well-characterized relative ages(*31*), their distribution provides temporal information about the history of the centromeric region. L1P1 elements correspond to very recent insertions, whereas L1PA2–L1PA4 subfamilies are older but still belong to the group of primate-specific LINE1 elements that were active during great ape evolution. The presence of multiple L1 subfamilies in human, chimpanzee, bonobo, and gorilla indicates that transposable element activity has repeatedly affected the chromosome 10 centromeric region since the divergence of these lineages. Notably, the estimated ages of the detected L1 subfamilies suggest that most insertions are younger than the divergence between orangutans and the African great apes (~19 Mya). L1PA4 elements are estimated at ~18 Mya and L1PA3 at ~12.8 Mya, indicating that LINE activity within the α-sat region likely occurred after the orangutan speciation. This is consistent with lineage-specific accumulation of LINE elements contributing to the structural diversification of the chromosome 10 centromeric region.

806

#### 8. RNA sequencing and gene annotation

##### Methods

**Kinnex full-length RNA**

RNA was extracted from 3-6 million LCLs using the Qiagen RNeasy Mini Kit (Qiagen, 74104). RNA was quality-checked using UV-Vis spectroscopy (Denovix DS-11 FX) and the Agilent Bioanalyzer 2100 with the Total RNA Nano 6000 kit (Agilent, G2939A & 5067-1511). Kinnex full-length RNA libraries were generated per manufacturer's recommendations (PacBio, 103-072-000). Samples were sequenced on the Revio platform on SMRT Cells 25M with Revio SPRQ Chemistry (PacBio, 103-520-200) with Adaptive Loading and 30-hour movies. Data were postprocessed using SMRT Link v13.3 with the "Read Segmentation and Iso-Seq" pipeline to segment and classify reads.

**Allele-specific expression and gene annotation**

For full-length transcript analysis, reads were aligned to each of the six orangutan haplotype assemblies using minimap2 (v2.28;(*71*)). The consensus gene annotation of RefSeq and comparative annotation toolkit (CAT) generated by a previous study(*21*) was projected onto six orangutan haplotype assemblies via liftoff (v1.6.3;(*98*)). The index files of six orangutan haplotype genome assemblies were first generated using minimap2 (parameters: *minimap2 -ax splice -f 1000 --secondary=no --eqx -K 100M*). Then, reads were aligned with the command *minimap2 -ax splice -f 1000 --secondary=no --eqx* to the genomes assembled as part of the present study, and filtering of secondary and supplement alignments was performed using *samtools view -F 2308*. To investigate haplotype-specific gene expression, the alignment was repeated three times, using hap1- or hap2-only assemblies, and a composite assembly of both haplotypes. Stringtie3(*99*) was used to assess gene models underlying Iso-Seq data and to quantify read counts.

##### Results

**Kinnex full-length RNA**

Kinnex-seq sequencing on the Revio SPRQ platform generated high-quality, highly consistent data across samples; 0.33 SMRT cells were used per sample, yielding between 19 and 22 million HiFi reads with an average read length of ~2.3 kbp. The average full-length non-chimeric read length ranged from 2,305 to 2,322 bp, with median lengths around 2.1 kbp and N50 values between 2.4 and 2.42 kbp. Overall, each sample produced between 44.6 and 50.4 Gbp of data, with more than half of the reads exceeding 2 kbp in length (Table S27).

**Gene annotation projection and transcript analysis**

To investigate if the emergence of a neocentromere leads to differences in gene expression, we examined transcriptome sequencing from the three orangutan individuals. Prior to quantifying gene expression, we examined full-length transcript sequencing data to generate consensus gene models by aligning them to the matched genome assembly. We identified 964-1198 genes/haplotypes across Chromosome 10 (Table S28), including 139-262 putative novel genes without gene annotation. We observe that the PAB16 sample, which contains a heterozygous neocentromere, shows skewed gene expression in chromosome 10 (Fig. S15A). On the other hand, chromosome 10 of PPY17, which contains a neocentromere in hap1 and a normal centromere in the alternative haplotype, showed a comparable gene expression pattern to other chromosomes, suggesting the skewed expression is a PAB-specific property. Examining the genes flanking the original centromere, we find that the expression levels in the PAB16_hap1 possessing the normal chromosome 10 centromere, are greater than the alternative haplotype with neocentromeres (Fig. S15B-D). However, we observe that this differential expression is not only restricted to genes near the centromeres but also affects the entire chromosome. The higher-ranked gene-expression level in PAB16_hap1 was observed universally across chromosome 10, indicated by overall increased density of reds (Fig. S15E), as opposed to the lower expression of the PAB16_hap2.

We searched for evidence of expressed transcripts corresponding to the 252 kbp locus that was duplicated in the orangutan (termed SD1 and SD2). We identified two transcript models of the bicaudal D1 gene (*BICD1*), each supported by more than 10 full-length Iso-Seq transcript reads, and one additional, lowly expressed model supported by only five reads (Fig. S16). The latter was predicted to have a disrupted open-reading frame. Between the two stronger-support models, we find that the longer gene model 1 maps solely to the downstream copy (SD2), whereas the shorter gene model 2 shows expression at both duplicated loci (SD1 and SD2), corresponding to the PPY17_hap2. Given that the BICD1 protein product in humans homodimerizes(*100*), the presence of multiple isoforms could have functional consequences, such as a dominant-negative effect, as previously reported for short *BICD1* transcripts(*101*). Investigating the predicted translated protein sequence, we note an amino acid change (115 D>E) in the longer copy of all orangutan haplotypes compared to PPY17_hap2 (Fig. S17). In addition, we observe a substitution of the 12$^{th}$ amino acid (H>R) in the shorter *BICD1* transcript, distinguishing the additional copy in the PPY17_hap2 from the rest. Alphafold3(*30*), however, does not suggest any three-dimensional folding change as a result of this amino acid substitution.

880

#### 9. Phylogenetic analyses, TMRCA estimation, and selection analyses

##### Methods

**Short-read sequence data processing**

Short-read sequences produced in this study were processed together with samples from the literature(*20, 57, 66*). Adapters from fastq files were removed with AdapterRemoval(*102*) with the additional options: *--trims --minlength 35*. Mapping was performed with BWA-MEM(*103*) with the reference genome Susie_PABv2 (GCA_002880775.3). The resulting SAM files were converted into BAM format and sorted with SAMtools(*85*). Picard was used to remove duplicates with the command *MarkDuplicates -REMOVE_DUPLICATES true* and to assign individual read group names in the final BAM files with *AddOrReplaceReadGroups*. Single-nucleotide polymorphism (SNP) calling was performed with GATK(*104*), producing individual-specific GVCF files using the command *HaplotypeCaller* with the option *-ERC GVCF*. Later, individual GVCFs were combined (command *CombineGVCFs*), genotyped (command *GenotypeGVCFs*), and then SNPs were extracted (command *SelectVariants* with the option *--select-type-to-include SNP*). To use only reliable markers in subsequent analyses, a strict SNP filtering was performed in several steps: (1) SNPs called in positions where the reference base is ambiguous (N) were removed. (2) A depth filter was applied, removing the positions where the cumulative depth (DP value) was 50% higher or lower than the average computed on all the sites. (3) Sites with Mapping Quality fraction (MQ0F) higher than 0.001 were excluded. (4) GATK was used to filter out SNPs based on the parameter indicated in the expression *QUAL < 50.0 || QD < 2.0 || FS > 60.0 || MQ < 40.0 || SOR > 3.0 || MQRankSum < -12.5 || ReadPosRankSum < -8.0* with the command *VariantFiltration*. (5) Multiallelic variants were removed since they may be representative of sequencing errors or other artifacts. (6) Finally, SNPs where there is a great disproportion of read depth in the two alleles in at least one heterozygous sample were discarded, in particular if the ratio between the number of alternative allele reads and the total reads was higher than 0.8 or lower than 0.2.

**Multiple alignment of long-read assemblies**

The alignment of long-read assemblies for chromosome 10 was performed with Progressive Cactus v2.9.8(*105*). The dataset included eleven sequences from two orangutan individuals (PAB and PPY), each represented by a primary and an alternate haplotype(*21*) (GenBank accessions GCA_028885655.3, GCA_028885685.2, GCA_028885625.3, GCA_028885525.2), the human T2T-CHM13 reference (NC_060936.1;(*96*)), and six haplotypes from in this study (PAB16_hap1 and hap2, PPY15_hap1 and hap2, PPY17_hap1 and hap2). HAL output was converted into MAF format with hal2maf using the options *--onlyOrthologs --noAncestors --noDupes --unique --maxRefGap 500 --maxBlockLen 500000*. Redundant blocks were removed with mafDuplicateFilter, and FASTA sequences were generated from the filtered MAF with mafToFastaStitcher, which concatenates orthologous blocks for each assembly(*106*). For the neocentromeric region, the core coordinates from each assembly were extracted, along with 250 kbp of upstream and downstream sequence. These sequences were aligned with MAFFT v11.0.13(*76*) using the options *--reorder --adjustdirection*, producing a focused alignment while retaining genomic context. SNP calling was performed with SNP-sites(*107*), generating a multi-sample VCF containing single-nucleotide variants.

**Pairwise alignments**

Pairwise alignments of chromosome 10 assemblies were performed between haplotypes (including intra-individual pairs) using Progressive Cactus v2.9.8. HAL files were converted to MAF using the same filtering options as in the previous section and were further processed with mafDuplicateFilter(*106*).

**Phylogenetic analyses**

Phylogeny was reconstructed using an MSA of both the chromosome-wide and neocentromere regions. We used IQ-TREE v2.4.0(*93*) with model selection (-m MFP) and 1000 ultrafast bootstrap replicates. Trees from both datasets were visualized in MEGA11 v11.0.13(*78*).

**TMRCA estimation**

The time to the most recent common ancestor (TMRCA) was estimated using the .mfa FASTA files generated from pairwise alignments of chromosome 10(*105, 106*). Analyses were performed in RStudio (version 2022.12.0+353) with the R packages Biostrings v2.74.1(*108*), stringr v1.5.1 (https://github.com/tidyverse/stringr) and ggplot2 v3.5.2(*68*), using a sliding-window approach (500 kbp windows with a 100 kbp step). Only non-gap(*68*) sites were retained, and windows with more than 80% missing data were excluded. For each window, divergence was also measured independently in MEGA (v11.0.13;(*78*)), using the option Compute Pairwise Distances with the method Number of Differences, which reports the absolute count of variable sites between sequences. Sites containing gaps or missing data were excluded with pairwise deletion. TMRCA in years was calculated as follows: $TMRCA = d/2\mu \times g$, with a mutation rate $g=1.5\times10^{-8}$ substitutions per site per generation and a generation time $g=25$ years(*57*).

**Selection**

Starting from the same SNPs dataset described in the section ‘Short-read sequence data processing’, two distinct VCF files were obtained, one for each species, with BCFtools v1.17 (*option view -S*)(*85*). PPY19 was excluded because of its hybrid origin. Tajima’s D (*109*) was computed with VCFtools v0.1.16(*110*) on windows of 1 Mbp along chromosome 10 with the option --*TajimaD 100000*. The same species-specific databases were used to identify putative positive selection sweeps with SweepFinder2(*111*). First, the VCF files were converted into the allele frequency file required by SweepFinder2 with the command:

```
bcftools query -f '%CHROM\t%POS[\t%GT]\n' INPUT.vcf | awk
'BEGIN{OFS="\t"}{x=0;n=0;for(i=3;i<=NF;i++)if($i!="./."){split($i,a,"[/|]");x+=(a[1]=="1"
)+(a[2]=="1");n+=2}if(n)print $2,x,n,1}’
```

where INPUT.vcf represents the original VCF files. Subsequently, the scan for selective sweeps along chromosome 10 was performed with the original SweepFinder method(*112*) on windows of 1 Mbp with the command *SweepFinder2 -sg 1000000.*

#### Results

To investigate the evolutionary history of the neocentromeric region, we performed a phylogenetic analysis of the entire chromosome 10 and of the neocentromeric region alone. Maximum-likelihood inference from the full chromosome recapitulates the established evolutionary history of the genus by grouping the six PPY and the four PAB haplotypes into two distinct monophyletic clades (Fig. S18). The presence of these two discrete clusters is consistent with the separation between the Bornean and Sumatran lineages observed in the PCA (Fig. 1D).

In contrast, the phylogeny inferred from the neocentromeric region alone reveals signals that deviate from the established evolutionary relationships between the two species. Notably, the PAB haplotype carrying the neocentromere (PAB16_hap2) clusters within the PPY clade, whereas the remaining non-neocentromeric PAB haplotypes form a separate cluster (Fig. 4a). This pattern indicates that the neocentromeric haplotype of PAB is phylogenetically more similar to the haplotypes (both neocentromeric and non-neocentromeric) of PPY than to those of its own species and suggests that the neocentromeric region may have followed an evolutionary trajectory distinct from that of the surrounding genome. Moreover, within PPY, it is not possible to discriminate between neocentromeric and non-neocentromeric individuals, indicating that the presence of the neocentromere does not affect their phylogenetic relationships.

We also estimated the TMRCA of the neocentromeric region by comparing sequences from the two species. When we compare the neocentromeric region between PAB CEN and PPY NEO, we obtain a coalescent time of approximately ~3.33-3.61 Mya, which is comparable to that inferred for the entire chromosome and is ≈3,4 Mya (Figs. 4c-d). In contrast, comparisons involving the neocentromeric haplotype of PAB, such as PAB NEO vs. PPY NEO or PAB NEO vs. PPY CEN, yield coalescent times of ~0.32-0.42 Mya, which postdate the species divergence and are markedly more recent than those estimated from the entire chromosome. Such a recent coalescent time between PAB NEO and the corresponding region in PPY (both NEO and non-NEO) might suggest a secondary introgression event from PPY into PAB, potentially introducing the neocentromeric region, harbouring an already active neocentromere or one predisposed to activation, into the PAB genomic background. To assess the relevance of the centromeric regions in the evolutionary dynamics of the two species, we performed selection scans along chromosome 10 separately for PPY and PAB (Fig. S19). In PPY, it is possible to observe strong selective sweeps, both for the centromeric and the neocentromeric region, while PAB shows no sign of selection. This result highlights strong selective pressures on centromeres in PPY, suggesting that complex evolutionary dynamics are operating in these regions.

1009

#### 10. Incomplete lineage sorting (ILS) and posterior decoding

##### Methods

**Processing genome alignments using MafFilter**

We started a Multiple Alignment Format (MAF) file containing haplotype-resolved blocks of orangutan chromosome 10 aligned against the human T2T reference (T2T-CHM13) homologous chromosome 12. The targeted set of species for the analysis were PAB16_hap1 (α-sat(+)_Mono), PAB16_hap2 (α-sat(-)_Neo), and PPY15_hap1 (α-sat(-)_Neo). We used T2T-CHM13 as an outgroup to consider the following tree:

```
(((PAB16_hap1, PAB16_hap2), PPY15_hap1), T2T-CHM13)
```

MAF input was filtered with MafFilter (v1.3.1)(*113*) using the *Subset* filter to extract four-way orthologous alignment blocks. Only blocks containing at least one sequence from each selected species have been kept (*strict=yes*), and blocks with duplicated sequences from at least one species were removed (*remove_duplicates=yes*). Additional input flags were used to interpret dotted characters in block sequences as gaps (*input.dots="as_gaps"*), and inconsistency of size sequences was specified (*input.check_sequence_size=F*). The output consisted of a four-way-filtered MAF file with no masking sequences (*mask=no*).

```
maffilter input.file="{input}" input.check_sequence_size=F input.dots="as_gaps" \
maf.filter="Subset{species=({specie1}, {specie2}, {specie3}, {outgroup}), strict=yes,
remove_duplicates=yes}, Output(file="{output}", mask=no)"
```

Consecutive alignment blocks were merged using the *Merge* command. T2T-CHM13 was used as a reference for synteny checking of blocks due to its high-quality assembly, to avoid loss of sequences and prevent the retention of obsolete sequences in the merged consecutive alignment blocks.

```
maffilter input.file="{input}" \ maf.filter="Merge{species=({outgroup}), dist_max=0},
Output(file="{output}")"
```

We also discarded small blocks (*min_length=2000*) with no masking sequences in the output file (*mask=no*).

```
maffilter input.file="{input}" \ maf.filter="MinBlockLength(min_length=2000),
Output(file="{output}",mask=no)"
```

The resulting MAF file was used as input for ILS analysis together with demographic parameters derived from a maximum likelihood estimation (MLE) carried out by TRAILS software (v0.0.1;(*114*)).

**Demographic parameters optimization**

The optimization process aimed to estimate demographic parameters of the observed genomic alignment data. These include the population size parameters (*N_AB*; *N_ABC*), divergence time parameters (*t_A*; *t_B*; *t_C*; *t_2*; *t_upper*), and other relevant parameters such as the recombination rate ($\rho$). Moreover, fixed parameters were used in the optimization process, including the number of discretized time intervals before the split events in the reconstruction of the phylogenetic tree (*n_int_AB_original=3*; *n_int_ABC_original=3*) and the mutation rate

($\mu$ = 2 × 10^-8). The optimization procedure employed the Nelder-Mead algorithm for model fitting, starting from a nine-way Cactus alignment (MAF file) with the following species selected for the demographic inference: PAB16_hap1 (α-sat(+)_Mono), PAB16_hap2 (α-sat(-)_Neo), PPY15_hap1 (α-sat(-)_Neo), and T2T-CHM13. Demographic parameters optimization was performed using a Python (v3.12.9) script adapted from TRAILS software (v0.0.1). A second Python script was used to extract the optimized parameters with the highest likelihood. All optimized parameters are listed in Table S29.

**Posterior decoding**

Posterior probabilities were computed by TRAILS software (v0.0.1), which implements a probabilistic model based on interconnected continuous-time Markov chains (CTMCs). These CTMCs model transitions between hidden states, each corresponding to a distinct genealogical topology and a pair of discretized coalescent time intervals, through the coalescent with recombination process across two adjacent genomic sites. Emission probabilities of hidden states, representing the probability of observing the alignment data corresponding to a given genealogy, were calculated using separate CTMCs that model a mutational process. All CTMCs were parameterized using the demographic parameters estimated during the optimization process, including the effective population size (Ne), speciation time (t), recombination rate ($\rho$) and mutation rate ($\mu$). TRAILS used transition and emission probabilities of hidden states to estimate the likelihood of each genealogy, defined by a unique combination of topology and two coalescent time intervals, at each nucleotide position within posterior probabilities. Additionally, other parameters required for the posterior decoding—such as the discretized time intervals—were set consistently with those estimated in the demographic parameters optimization (*n_int_AB_original* = *n_int_AB*; *n_int_ABC_original* = *n_int_ABC)*. Posterior decoding was computed using a Python (v3.12.9) script executable via command-line arguments using the argparse module.

```
python argparse_compute_posteriors.py --maf “{input}” --sp1 “{specie1}” --sp2 “{specie2}”
--sp3 “{specie3}” --outgroup “{outgroup}” --n_int_AB_original “3” --n_int_ABC_original
“3” --n_int_AB “3” --n_int_ABC “3” --parameters “{parameters}” --output “{output}”
```

**Processing posterior decoding data**

The ILS pattern and coalescent time in PAB16_hap1, PAB16_hap2, PPY15_hap1, and T2T-CHM13 quartet were analyzed using R Studio (v2024.12.0). First, a weighted posterior average probability was calculated for each possible genealogy, excluding missing genomic positions. Then, the ILS proportion was estimated by summing the mean posterior probabilities of hidden states that share ILS topology (A and B) and dividing the sum by the total probabilities across all topologies. Using a similar procedure, a weighted mean posterior probability was calculated for each discretized time interval for the first and second coalescent events, again using 1 kbp windows. In particular, the first and second coalescences were calculated by assigning integers from 1 to 3 to each discretized interval in chronological order, backward in time, and then computing a weighted mean of the integers using the corresponding posterior probabilities per 1 kbp window as weights. Windows with more than 50% missing data were then excluded, and additional filters were applied to remove outliers below the 1$^{st}$ percentile and above the 99$^{th}$ percentile for ILS and coalescence analysis. To highlight the variation in ILS and the coalescent time across the entire orangutan chromosome 10 (hsa12), particularly around the neocentromeric region, the processed values were summarized in 100 kbp windows by averaging 1 kbp window values. All data processing was performed using Rscript (RStudio v2024.12.0) executable via command-line arguments using the optparse module (v1.7.5).

```
Rscript tab_ILS_V2_V3_1stC_2ndC.R --input "{input}" --hidden_states "{hidden_names}"
--bin_size "{bin_size}" --chr 10 --output "{output}"
```

To optimize and exemplify the workflow, a Snakemake (v9.0.1) script was used.

**Gene annotation**

Gene annotation was performed using the filtered dataset of 100 kbp windows. To highlight the gene annotation along chromosome 10 (hsa12), the genomic coordinates of the twenty 100 kbp windows with the highest and the twenty with the lowest proportion of ILS (for a total length of 4 Mbp, ~3% of the length of the chromosome) were extracted using the slice_max and slice_min functions from the *data.table* package in R Studio (v2024.12.0). The extracted coordinates were saved in a BED format. Gene annotations were carried out using the intersect function from BEDTools (v2.31.1), which compares each feature in the query file (flag -a) with the features in a reference annotation file (flag -b) in search of overlaps. As a reference, we used a Verkko 2.2.1_v1 Liftoff mapped to PAB16_hap1 coordinates, in accordance with the TRAILS analysis outputs, where genomic coordinates are reported relative to the first species of the phylogenetic tree. The flags -wa and -wb were used to output the original entries from both input files for each detected overlap among them. The gene annotation was performed via the command line:

```
#highest 20
bedtools intersect -a /path/to/top20_highILS_100Kb_windows_clean.bed -b
/path/to/PAB16hap1_Verkko2.2.1_v1_liftoff.gtf -wa -wb >
outputs/genes_top20_highILS_100Kb_windows.tsv

#lowest 20
bedtools intersect -a /path/to/top20_minILS_100Kb_windows_clean.bed -b
/path/to/PAB16hap1_Verkko2.2.1_v1_liftoff.gtf -wa -wb >
outputs/genes_top20_minILS_100Kb_windows.tsv
```

#### <u>**Results**</u>

**Demographic parameters estimation**

Transition and emission probabilities were modelled using continuous-time Markov chains (CTMCs) parameterised by demographic parameters, including speciation times, ancestral effective population sizes, and recombination rate ($\rho$). Speciation times are expressed in units of "*generations x mutation rate*," and the effective population size in units of "*individuals x mutation rate*." Assuming a mutation rate of $\mu = 1.5 \times 10^{-8}$ per site per generation and a generation time of $g = 25$ years(*57*), the inferred speciation time appears substantially older than the divergence time reported in the literature of *Pongo* lineages. For instance, the two haplotypes from the PAB individual show a "splitting time" of approximately 5.5-6.4 Mya, and the PAB-PPY split is inferred at ~9.2 Mya, whereas the most recent common ancestor with the outgroup (*Homo Sapiens*, CHM13) is placed at ~27 Mya. These values should not be interpreted as direct dates of speciation. Instead, they likely reflect a single-chromosome estimate of coalescent times, which can differ substantially from species divergence events, especially in genomic regions characterised by different recombination—a low recombination could increase genealogical depth(*115-117*)—or in regions characterised by complex evolutionary structure, such as regional variation in mutation rate(*118*), which may impact the time conversions. The converted recombination rate ($\rho$) estimate was approximately $9.477 \times 10^{-9}$. Effective population size (*Ne*) estimates, converted using the same mutation rate ($\mu$), yielded two closely similar values in PAB16_hap1-PAB16_hap2 ancestry and in PAB16_hap1-PAB16_hap2-PPY15_hap1 ancestry, namely 38,926 and 39,022 individuals, respectively, which are very close to previous estimates(*21*). Using the converted estimates for t2 (in generations) and N_AB, we calculated the proportion of ILS as 30,8%, an estimate that is slightly higher than the one obtained by harnessing the posterior decoding approach. Although the converted demographic parameter estimates appear slightly inflated, they are internally consistent, supporting the overall coherence of the inferred demographic model parameters.

**ILS in the neocentromeric region**

We investigated the ILS pattern along orangutan chromosome 10 (hsa12) by analyzing 100 kbp long windows in the PAB16_hap1, PAB16_hap2*, PPY15_hap1*, and T2T-CHM13 phylogenetic quartet. Across the entire chromosome, the mean of the ILS proportion was ~21.8%, of which ~10.5% supports the PAB16_hap1 and PPY15_hap1* ILS phylogeny –Topology A– and ~11.3% the PAB16_hap2* and PPY15_hap1* ILS phylogeny –Topology B–. Notably, the neocentromeric region exhibited a markedly elevated ILS proportion of 60.9%, with pronounced asymmetry between the two alternative ILS topologies. Only ~3.6% of windows supported the ILS-Topology A, whereas ~57.3% of windows supported the ILS-Topology B (Fig. S20). The PAB16 haplotype, which carries the neocentromeric region, appears to cluster more closely with PPY15, which is homozygous for the presence of the neocentromere. Moreover, in the distribution of ILS-Topology A windows, the ILS proportion observed at the neocentromere approaches the mean of the distribution. In contrast, in the distribution of ILS-Topology B, it exceeds the $99^{th}$ percentile ($99^{th}$ percentile threshold=0.427), indicating a robust affinity among the PAB16_hap2* and PPY15_hap1* haplotypes in the neocentromeric region (Fig. S20). We further examined first and second standardised coalescent times along the orangutan chromosome 10 (hsa12) using the same window-based framework (Fig. S21). Windows overlapping the neocentromeric region exhibited a first coalescent time approximately 1.54 standard deviations (s.d.) below the mean and a second coalescent time approximately 2.1 s.d. above the mean, resulting in one of the highest values in the distribution. These patterns indicate a relatively shallow coalescent between PAB16_hap1 and PAB16_hap2*, as intrinsically expected for two haplotypes derived from the same individual, and a markedly deep coalescent between the PAB16 and PPY15_hap1*, consistent with an ancient Most Recent Common Ancestor (MRCA).

**Gene annotation at ILS hotspots and deserts**

Gene annotation performed on the twenty 100 kbp windows exhibiting the highest and lowest ILS proportion revealed a spectrum of configurations. These included deserts or hotspots characterised by a specific topology in which one or more genes were present, as well as windows in which no genes were detected (Fig. 4C).

At the neocentromere, one ILS hotspot ranked among the top twenty highest ILS 100 kbp windows across the chromosome (specifically, the $4^{th}$ highest ILS value; Table S30). This window displayed a predominant proportion of Topology B, yet no genes were detected within it. Multiple additional hotspots were identified in the neighbourhood of the neocentromere, evolving an extended cluster of elevated ILS. In all of these windows, no genes were detected except for two hotspot 100 kbp windows: 83-83.1 Mbp and 83.1-83.2 Mbp, in which *SLC6A15* (chr10:83,073,275-83,126,278) and *LOC112136027* (chr10:83,106,351-83,106,540) were identified, respectively (Table S30). A detailed visualization of this pattern and the corresponding gene annotation was obtained by a zoom-in around the neocentromere region (Fig. S22).

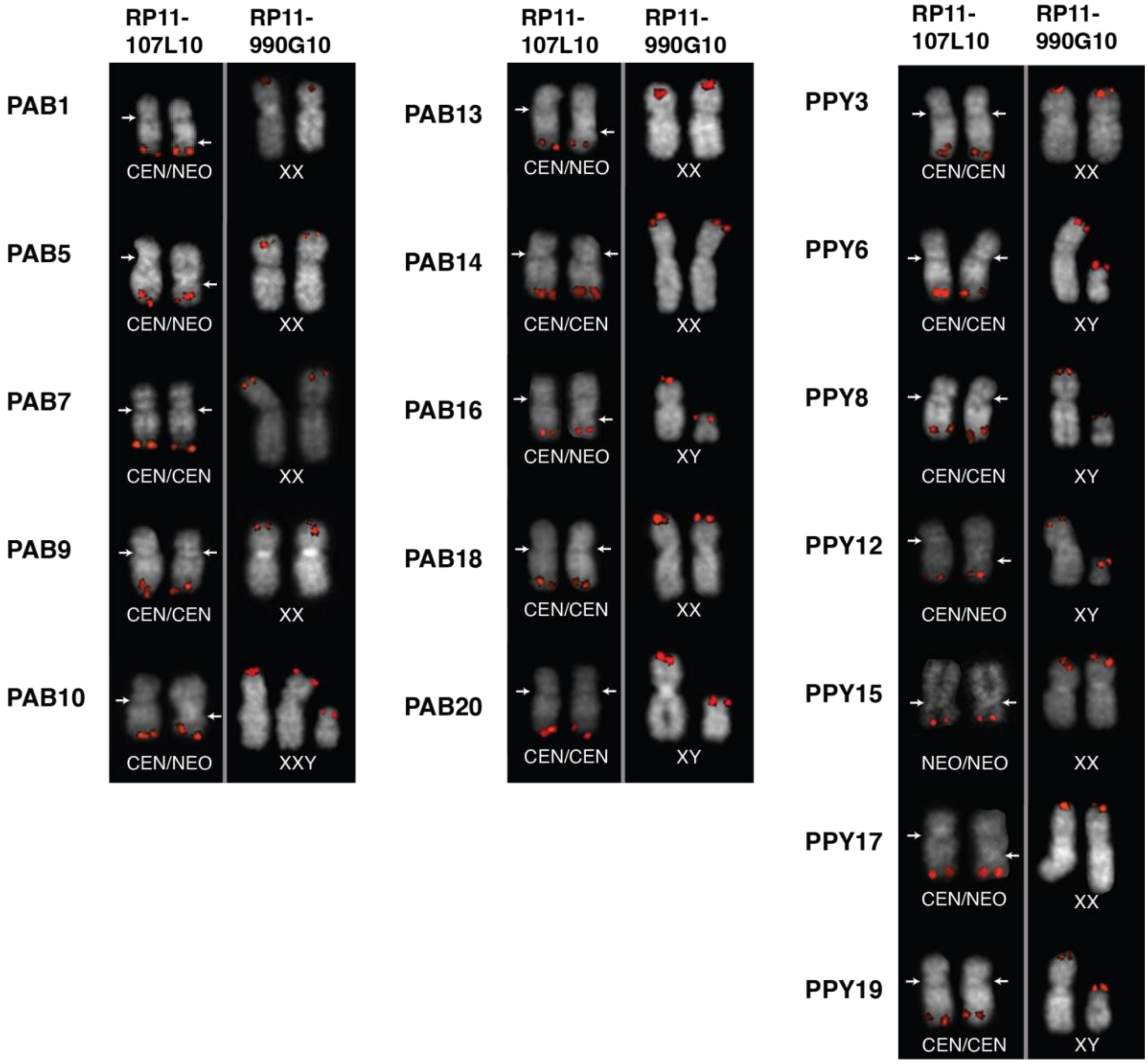


**Fig. S1. FISH experiments.** FISH experiments were performed on the 17 orangutan LCLs using the RP11-107L10 probe to investigate the neocentromeric status and the RP11-990G10 probe, specific for the Pseudo Autosomal Region, to determine chromosomal sex. Chromosomes were stained with DAPI; signals were pseudo-colored in red. Arrows indicate the position of the active centromere on each chromosome 10.

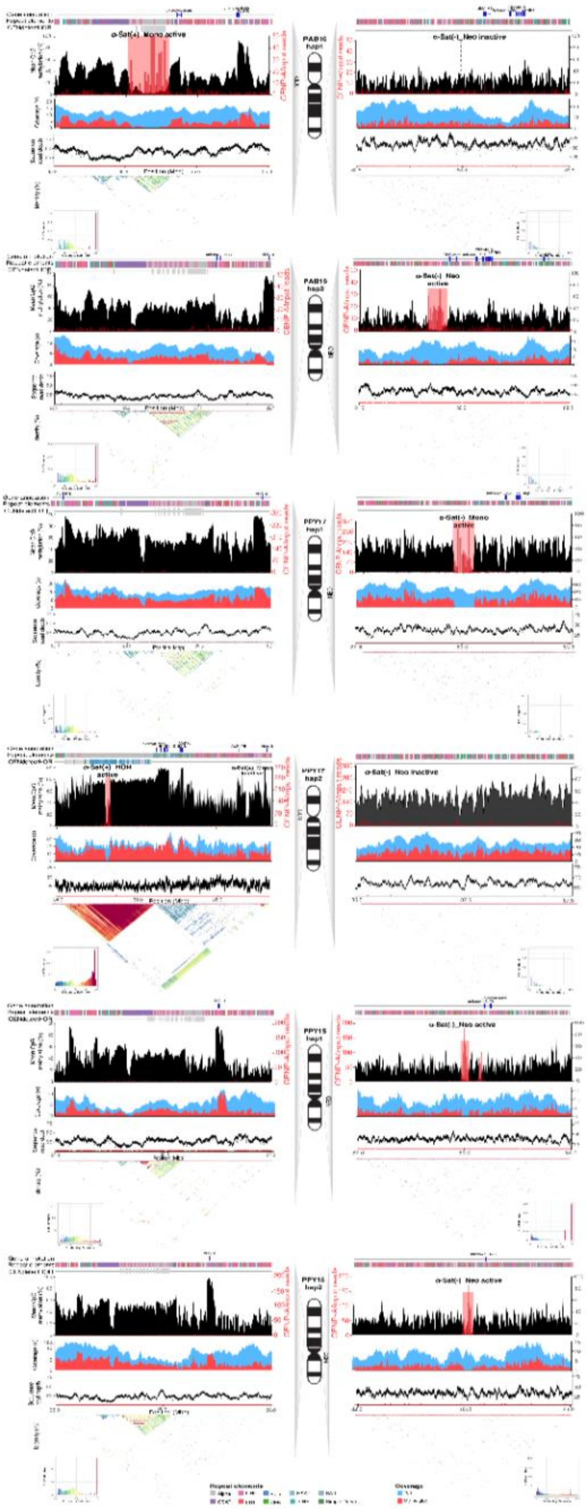

**Fig. S2. Centromere organization of chromosome 10 in samples PAB16, PPY17, and PPY15, for both haplotypes (hap1 and hap2).** For each haplotype, for the centromeric (left) and neocentromeric (right) regions, the following are shown from top to bottom: (i) gene annotation; (ii) composition of repeat element classes; (iii) organization of α-sat units, distinguishing between α-Sat(+)_Mono, α-Sat(+)_HOR, and α-Sat(−)_Neo, further classified as active or inactive based on methylation profiles (black) and ratio (red) of CENP-A chip reads/input reads (5 kbp bins); (iv) coverage; (v) sequence read depth; and (vi) sequence identity percentage.

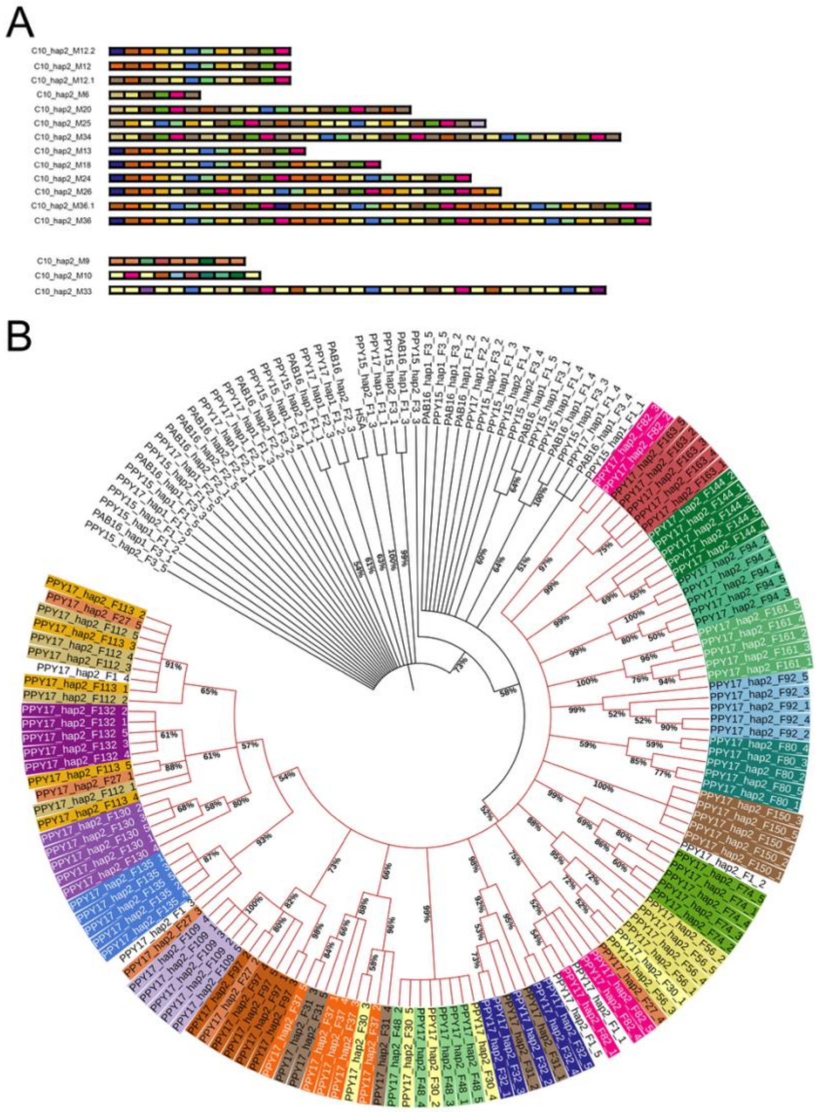


**Fig. S3. HOR organization and phylogenetic relationships of α-sat monomer families.** HOR organization and phylogenetic relationships of α-sat monomer families. (A) HOR structures identified in hap2 of PPY17, showing the detailed composition of C10_hap2_M12.2 and its variants in terms of monomeric families. C10_hap2_M9, C10_hap2_M10, and C10_hap2_M33 create additional HORs. Blocks of the same color represent monomers belonging to the same family. (B) Phylogenetic tree based on five representative sequences per family from all α-satellite families forming either HORs or monomeric arrays across the six analyzed haplotypes, together with the human α-satellite consensus sequence (HSA). Tip labels are colored according to monomeric family, consistent with panel A. Bootstrap support values (percentage of 1000 replicates) are indicated for major nodes. Two major clades are observed: the HOR-associated clade (in red) comprises all families from PPY17 haplotype 2, including both HOR-associated and monomeric families, whereas the second clade contains families from the remaining haplotypes, which are exclusively organized as monomeric arrays.

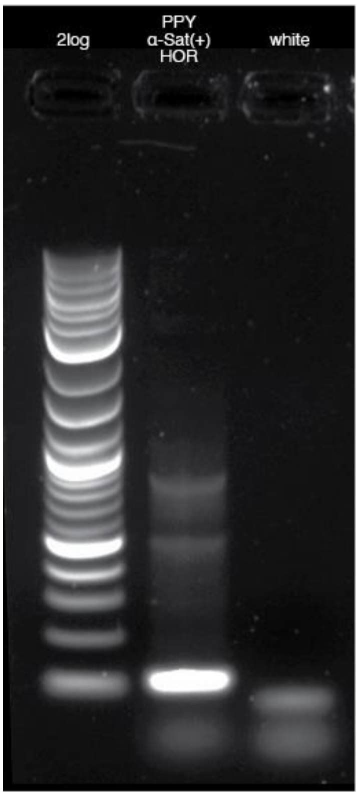


**Fig. S4. 1% agarose gel electrophoresis showing the PCR amplification products obtained from the PPY17 genomic template in PPY_α-sat(+)_HOR experiment.** Lane 1 (2log): 2-log DNA ladder used as size marker. Lane 2 (PPY_α-sat(+)_HOR): PCR products obtained in the PPY_α-sat(+)_HOR experiment. Lane 3 (white): negative control, showing no amplification.

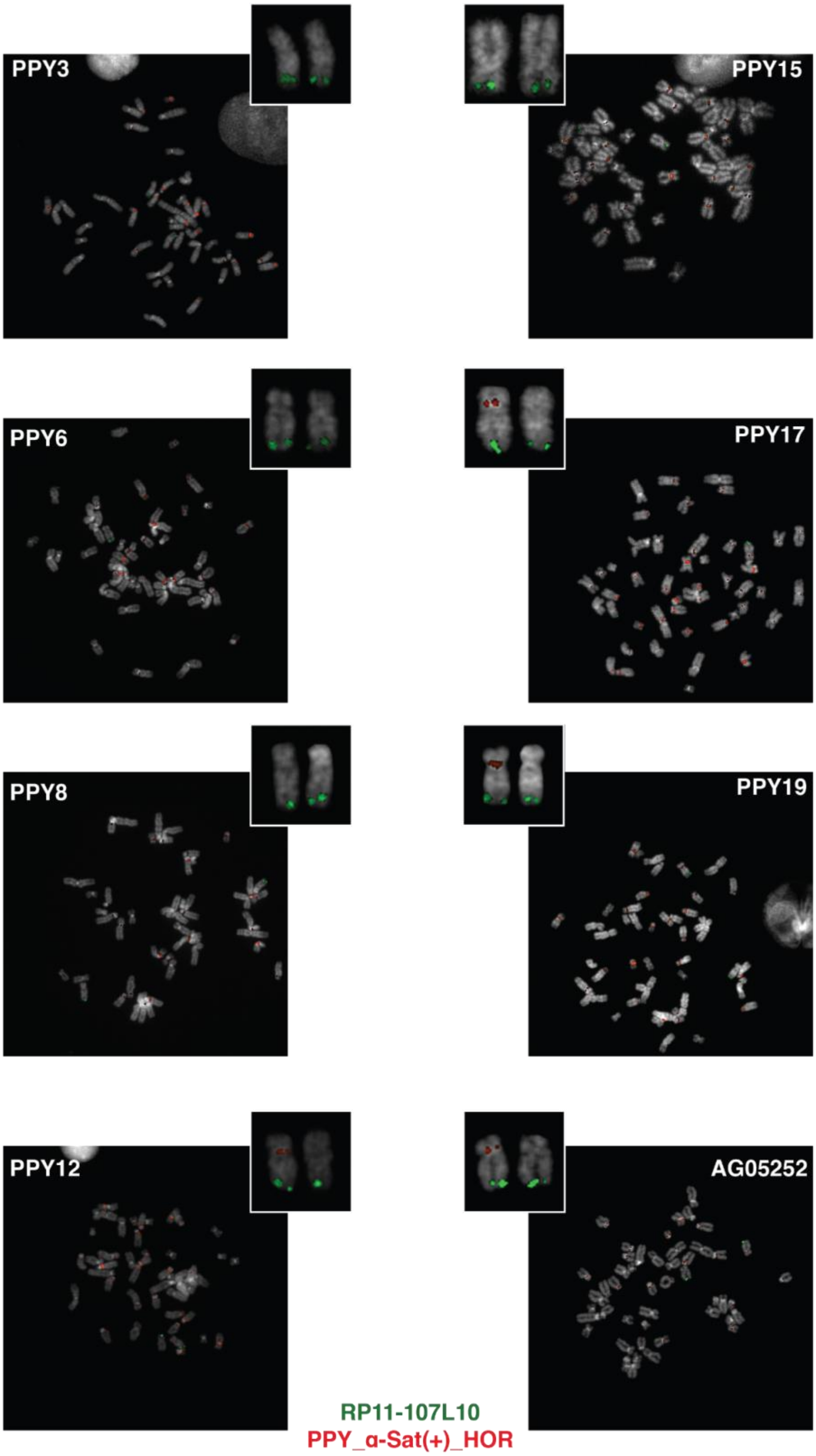


**Fig. S5. FISH analyses on the PPY individuals with the labeled 101 bp PCR product designed on PPY_α-sat(+)_HOR sequence.** PPY_α-sat(+)_HOR (red signals), co-hybridized with the BAC probe RP11-107L10 (green signals), performed on the PPY individuals, including the AG05252 cell line.

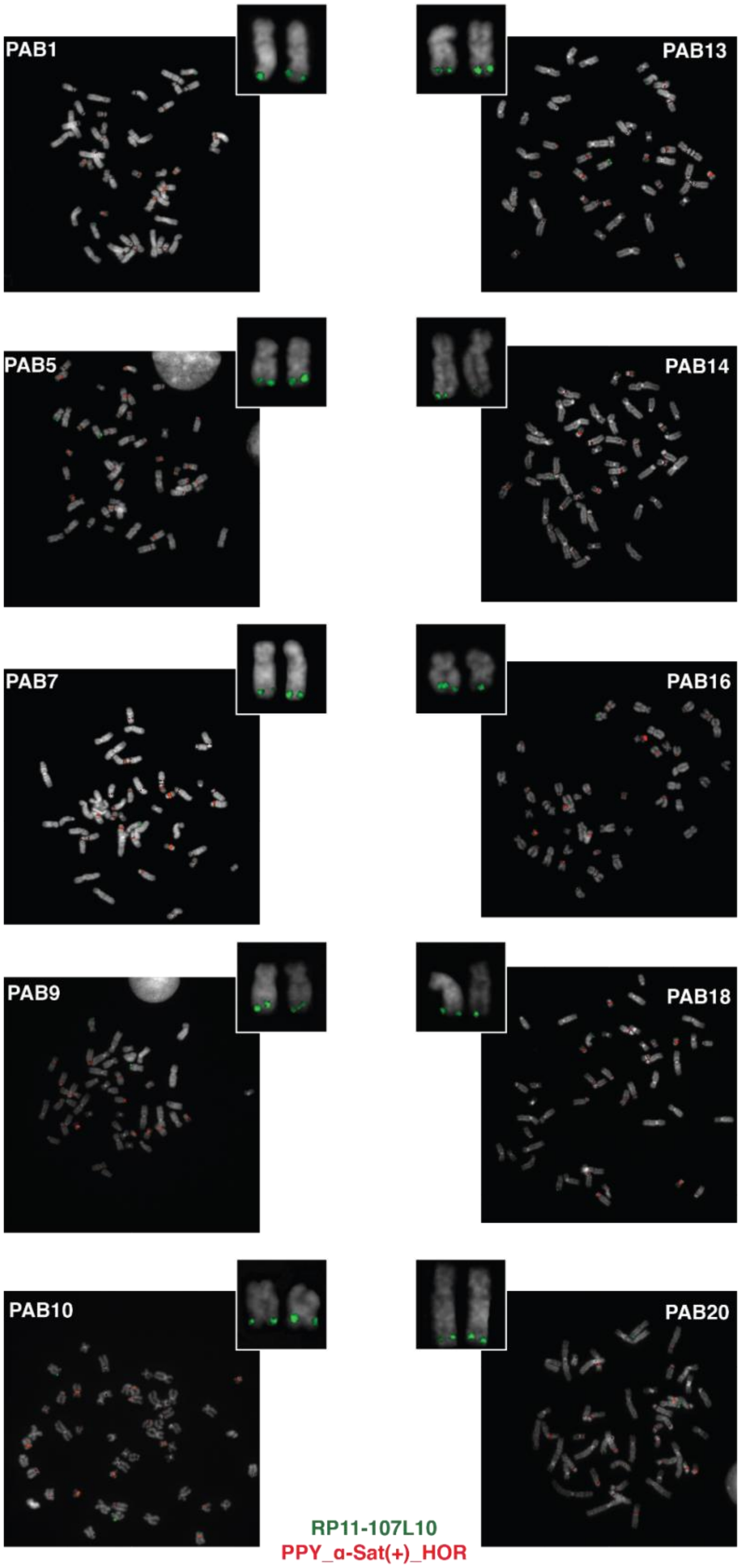


**Fig. S6. FISH analyses on the PAB individuals with the labeled 101 bp PCR product designed on PPY_α-sat(+)_HOR sequence.** PPY_α-sat(+)_HOR (red signals), co-hybridized with the BAC probe RP11-107L10 (green signals), performed on the PAB individuals.

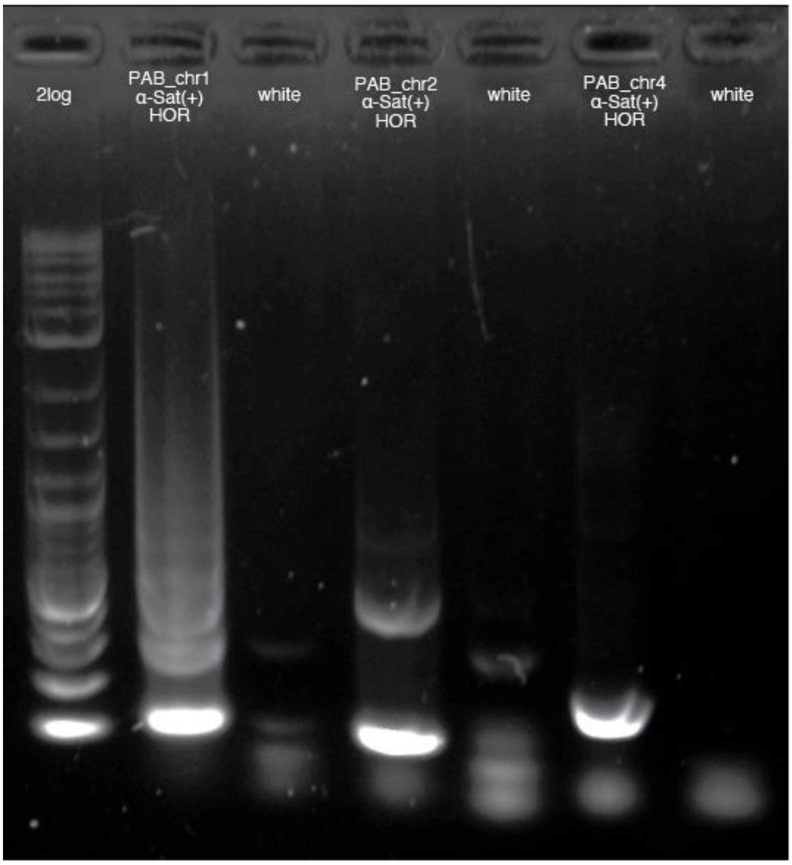


**Fig. S7. 1% agarose gel electrophoresis showing the PCR amplification products obtained from the PAB16 genomic template in experiments PAB_chr1_α-sat(+)_HOR, PAB_chr2_α-sat(+)_HOR, and PAB_chr4_α-sat(+)_HOR.** Lane 1 (2log): 2-log DNA ladder (1 kbp ladder). Lanes 2, 4, and 6: PCR products obtained with primer pairs designed on chromosomes 1, 2, and 4, respectively. Lanes 3, 5, and 7: corresponding negative controls showing absence of amplification.

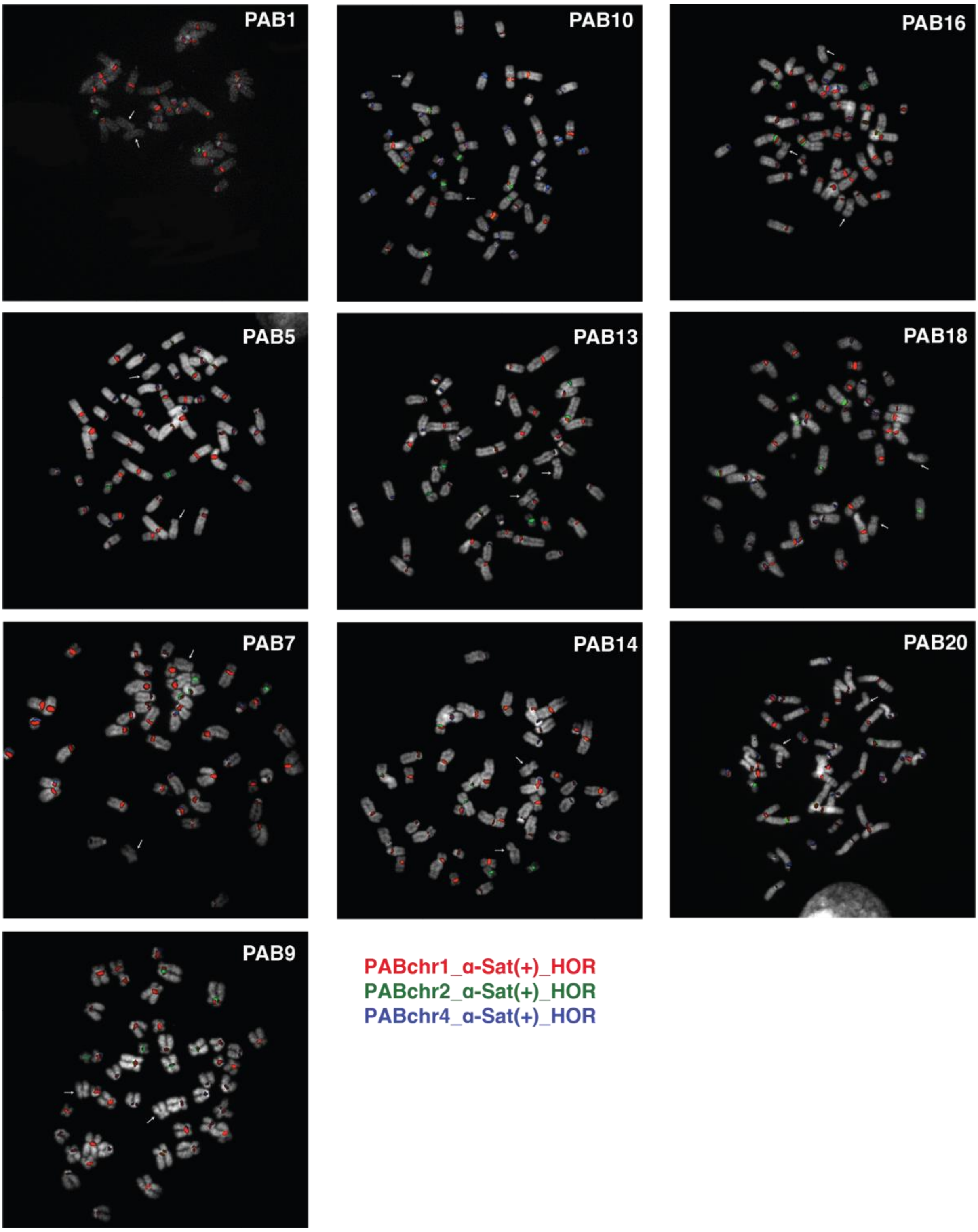


**Fig. S8. Results of the three-color FISH analyses.** FISH performed on PAB individuals co-hybridizing the three labeled PCR products obtained from experiments PAB_chr1_α-sat(+)_HOR (106 bp probe, pseudo-colored in red), PAB_chr2_α-sat(+)_HOR (101 bp probe, pseudo-colored in green), and PAB_chr4_α-sat(+)_HOR (190 bp probe, pseudo-colored in blue). Chromosomes were stained with DAPI. Arrows indicate each copy of chromosome 10.

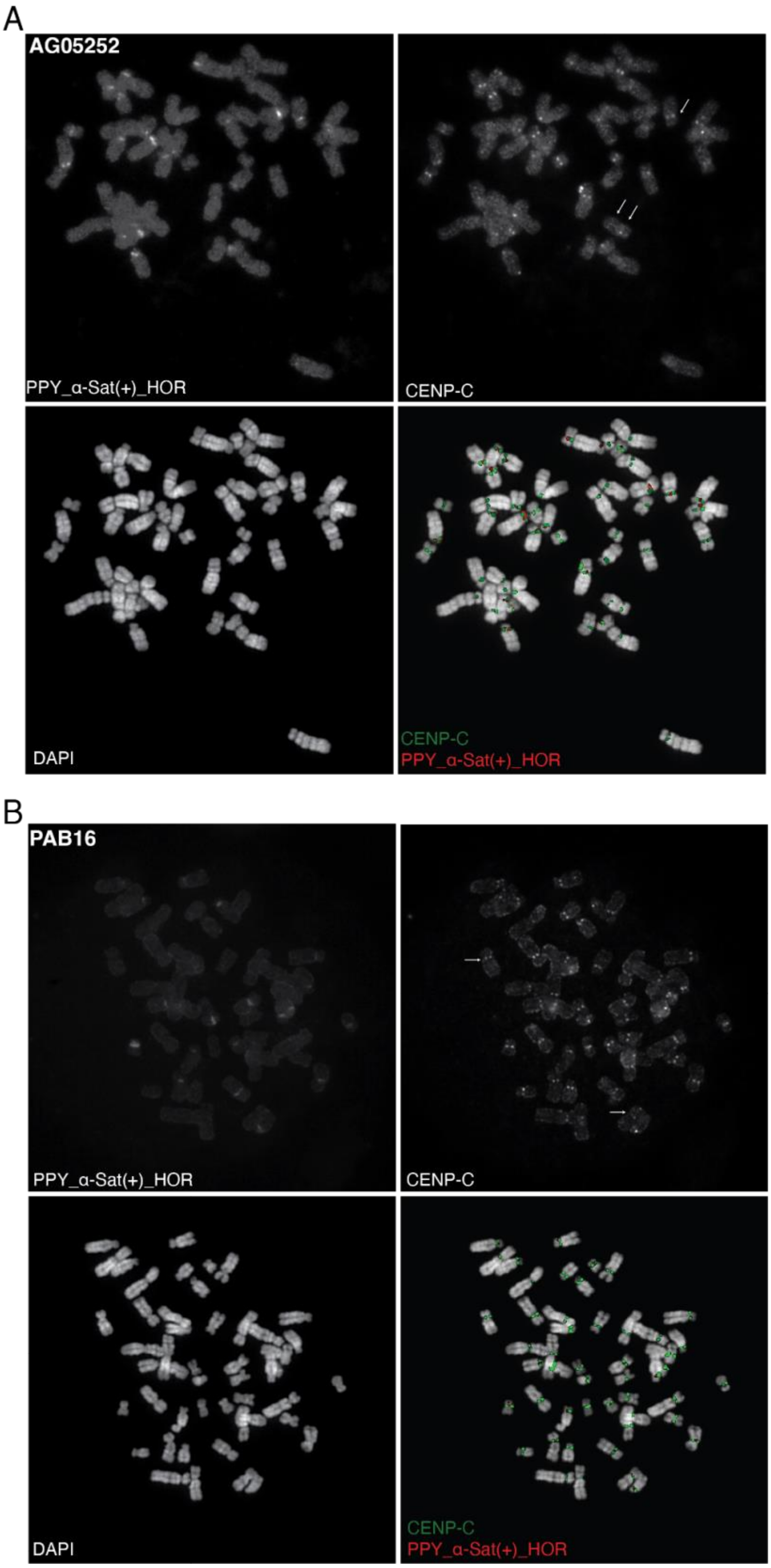


**Fig. S9. Immuno-FISH on AG05252 and PAB16 metaphases using CENP-C antibody and PCR product PPY_α-sat(+)_HOR.** Immuno-FISH on **(A)** AG05252 and **(B)** PAB16 metaphases. Panels show PCR product PPY_α-sat(+)_HOR FISH probe (Cy3), CENP-C immunostaining (Fluorescein), and DAPI, as greyscale images, and merged images for each sample. Arrows in the Fluorescein panels indicate CENP-C binding sites on chromosome. 10.

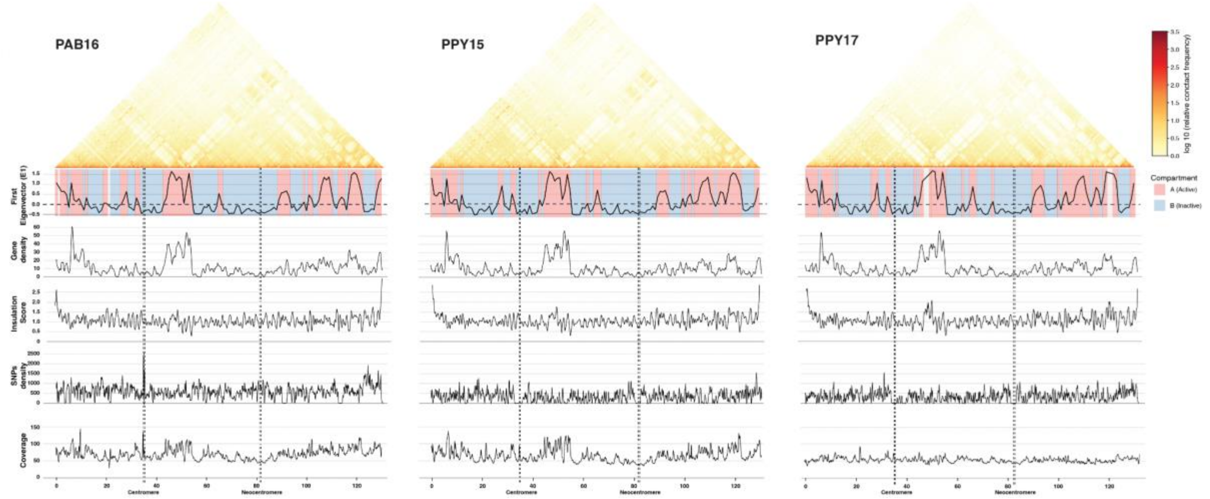


**Fig. S10. Genome-wide Micro-C contact maps and chromatin features along chromosome 10 in samples PAB16, PPY15, and PPY17.** The upper-triangular contact matrices show the normalized contact frequency (log10 scale), with enrichment of short-range interactions near the diagonal. Below each map, from top to bottom, the tracks display (i) the first eigenvector (E1) used to assign A (active, red) and B (inactive, blue) Hi-C compartments; (ii) gene density; (iii) insulation score, with minima in the signal indicating TAD boundaries; (iv) SNP density; and (v) coverage. Dashed lines indicate the position of the centromere.

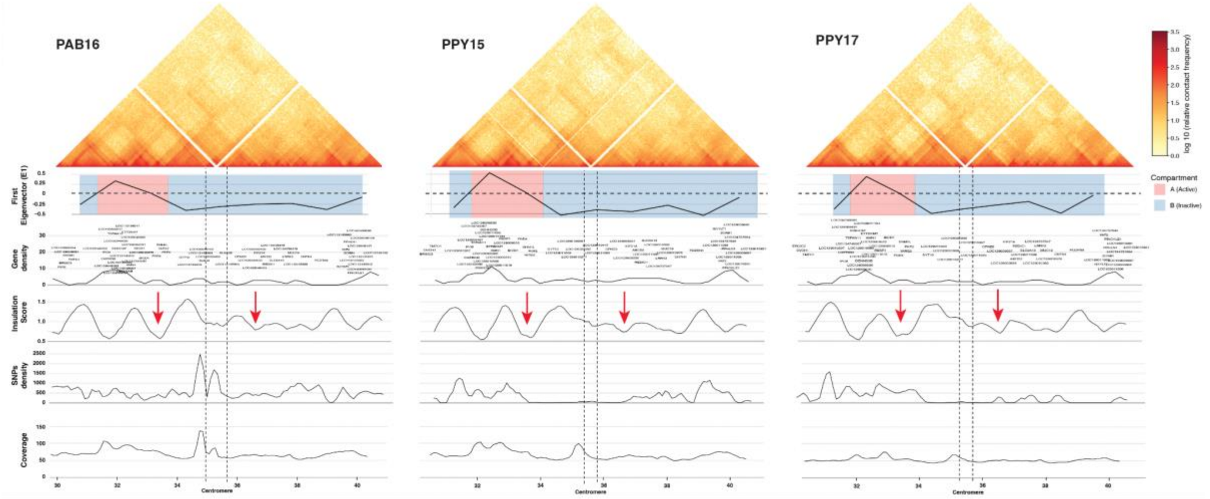


**Fig. S11. Zoom-in on Micro-C contact maps and chromatin features in a 10 Mbp region surrounding the centromere on chromosome 10.** The upper-triangular contact matrices show the normalized contact frequency (on a log10 scale). Below each map, from top to bottom, the tracks display (i) the first eigenvector used to assign A (active, red) and B (inactive, blue) Hi-C compartments; (ii) gene density; (iii) insulation score, with red arrowheads highlighting minima that indicate TAD boundaries surrounding the centromere; (iv) SNP density; and (v) coverage. Dashed lines indicate the position of the centromere.

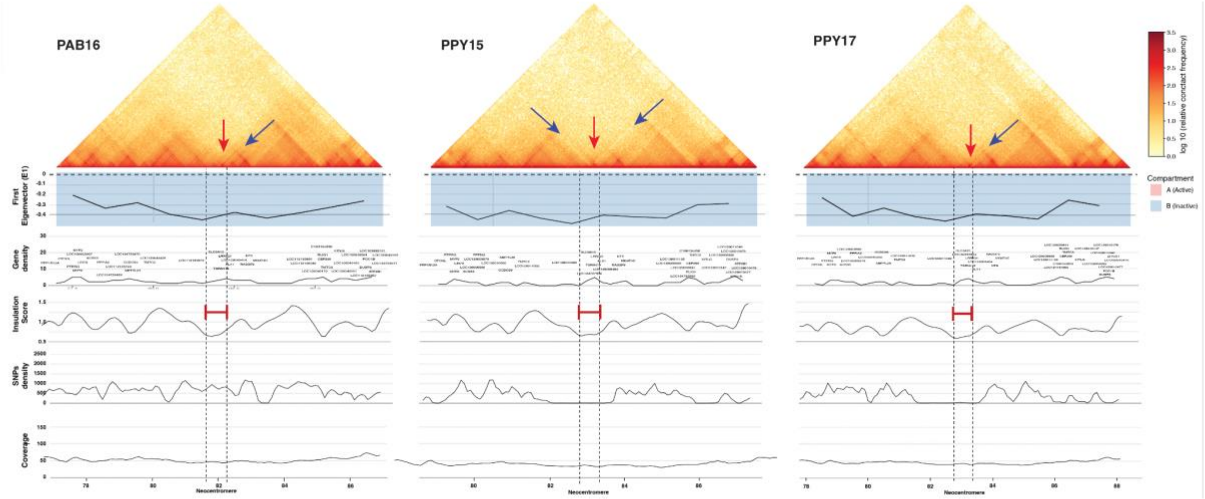


**Fig. S12. Zoom-in on Micro-C contact maps and chromatin features in a 10 Mbp region surrounding the neocentromere on chromosome 10.** The upper triangular Micro-C contact matrices show the normalized contact frequency (log10 scale). Red arrows indicate a small domain with depleted interactions in the surroundings in PPY15 (blue arrows indicating stripes with low contact signal in left and right orientations), whereas in PAB16 and PPY17 the domain forms a loop towards the TAD on the right (blue arrow indicating a stripe with increased contact signal). Below each map, from top to bottom, the tracks display (i) the first eigenvector used to assign A (active, red) and B (inactive, blue) Hi-C compartments; (ii) gene density; (iii) insulation score, with red line that highlights the extended valley that indicates the wide TAD boundary that overlaps neocentromere; (iv) SNP density; and (v) coverage. Dashed lines indicate the position of the centromere.

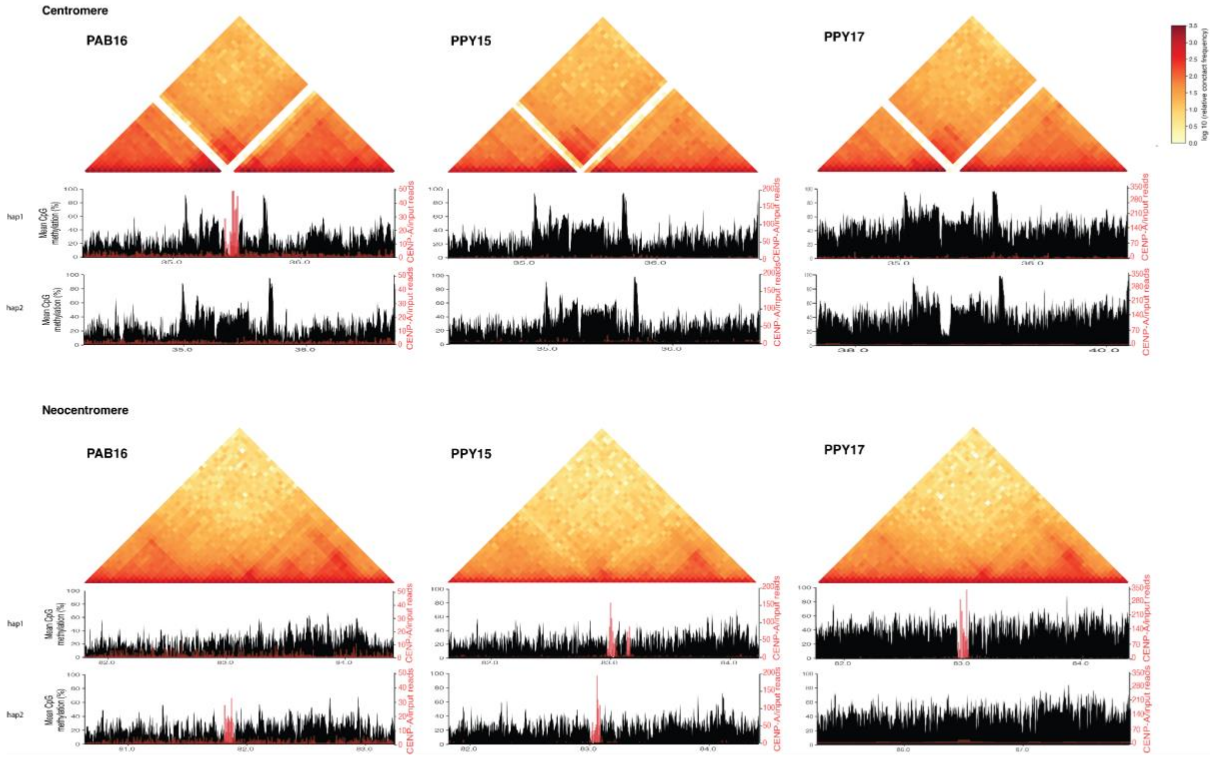


**Fig. S13. Further zoom-in on Micro-C contact maps in an approximately 2.5 Mbp region surrounding the centromere (top) and neocentromere (bottom).** Below each matrix, the mean CpG methylation and CENP-A signal for both haplotypes are provided. Non-allele-specific Micro-C maps for PAB16 were aligned to hap2, whereas PPY15 and PPY17 maps were aligned to hap1. For all cell types, alignment with the other haplotype is more approximate.

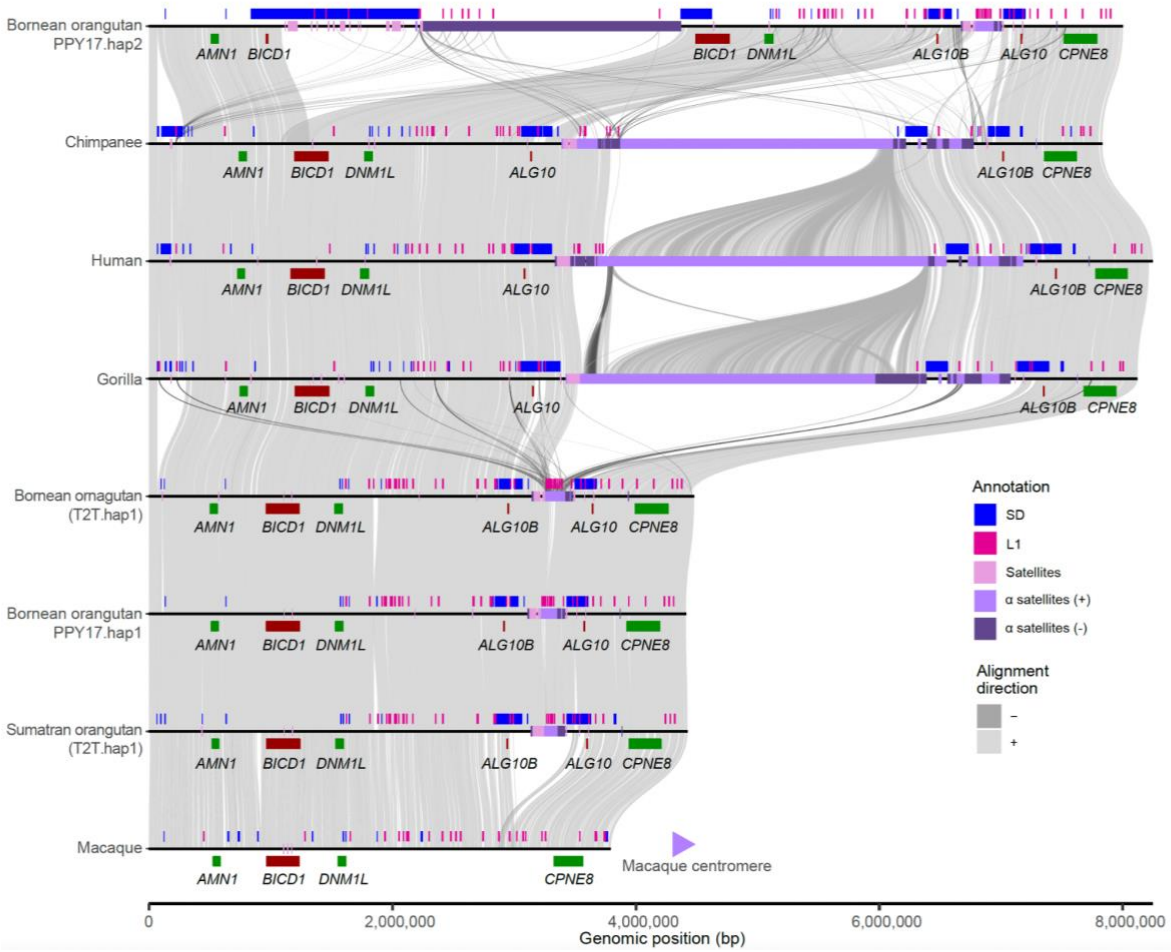


**Fig. S14. Comparisons of α-sat arrays across the ape lineage.** Sequence of PPY17_hap2 (32-39 Mbp) including the two α-sat arrays mapping to homologous regions of great apes (primary haplotypes of chimpanzee, human, gorilla, Bornean and Sumatran orangutans; (*21*, *99*)) and macaque (T2T-MFA; (*97*)). Alignment directions, whether direct or inverted, are indicated by light and dark grey, respectively. Annotation tracks including segmental duplication (SD; blue) and long interspersed nuclear elements-1 (L1; hot pink) are shown on the top track, followed by satellites (plum) in the middle. α-sats in direct and inverted orientations are indicated by light and dark purple, respectively. Below the satellite tracks are the genes showing synteny. Genes colored by red indicate those duplicated by at least one species, while green indicates unique genes.

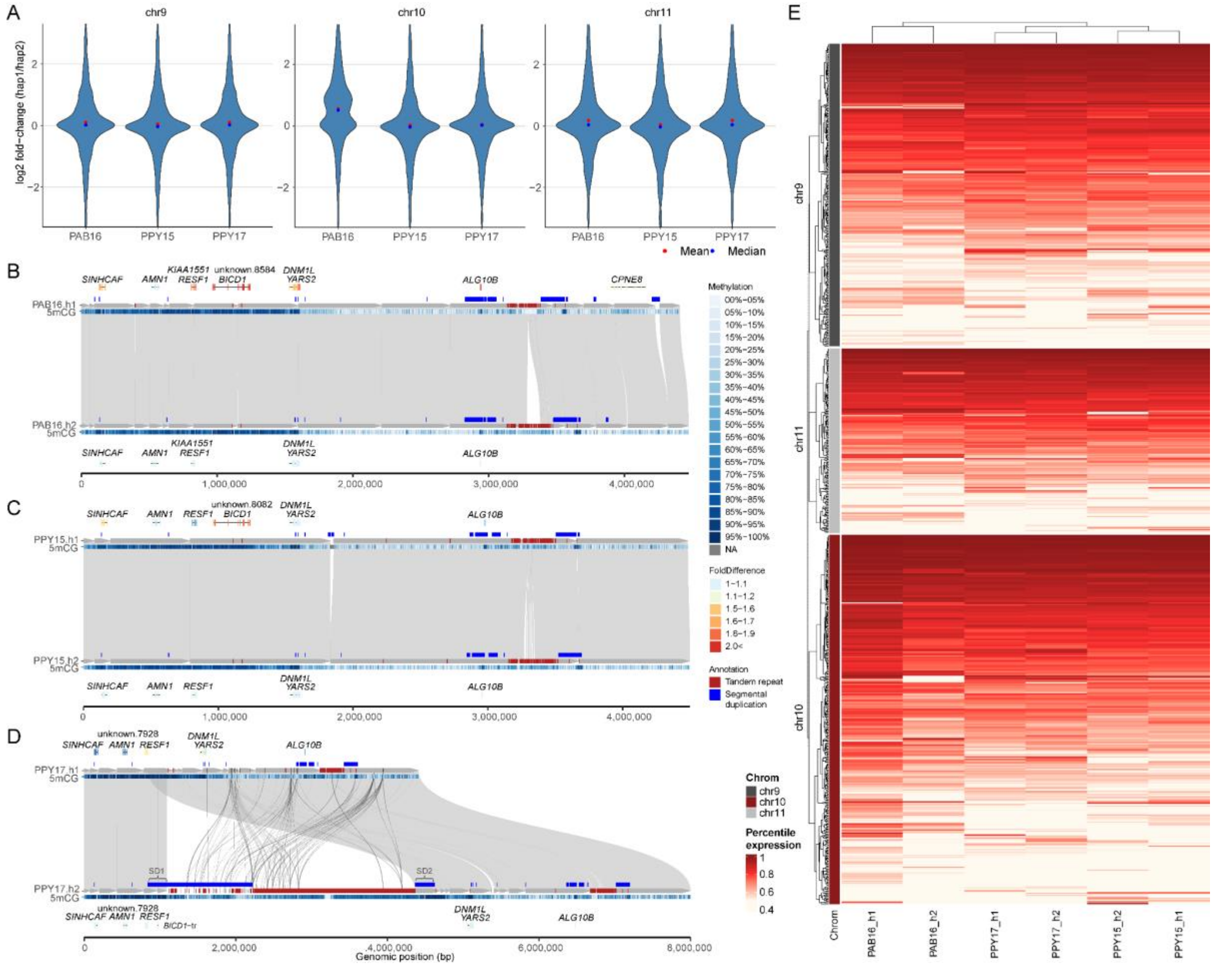


**Fig. S15. Haplotype-biased gene expression in heterozygous neocentromeric chromosome 10. (A)** Distribution of fold-change of gene expression (hap1/hap2) showing biased expression in PAB16. **(B-D)** Expression of genes neighboring the centromeric regions in **(B)** PAB16, **(C)** PPY15, and **(D)** PPY17. From the top, gene models are shown, with their colors indicating fold-change against the alternative haplotype (hap2). Below the gene, shows the SD track in blue, followed by centromeric satellites in red, as well as CpG methylation at the bottom, indicating hypermethylation in shaded of blue. **(E)** Heatmap of gene expression levels in percentile rank, indicating the highest expression in red to the lowest expression in white. The percentile rank was calculated from the genome-wide distribution, with each row representing a gene; the expression level of multi-copy genes is collapsed by summing the transcripts per million (TPM) counts.

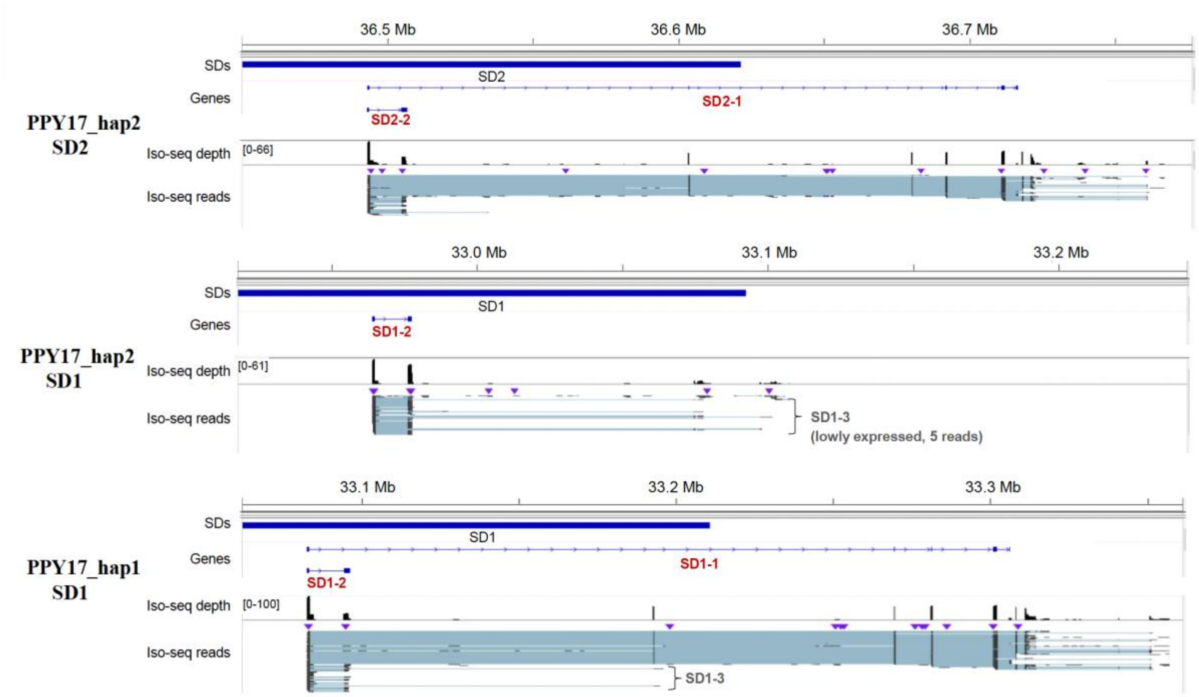


**Fig. S16. Three transcript models supported by at least five full-length transcript reads.** Evidence of gene models 1 and 2 (i.e., PPY17_hap2_SD2-1, PPY17_hap2_SD2-2, and PPY17_hap2.SD2-3) is found in SD2 (downstream of centromere) of PPY17_hap2 and the homologous sequence (SD1) in PPY17_hap1.

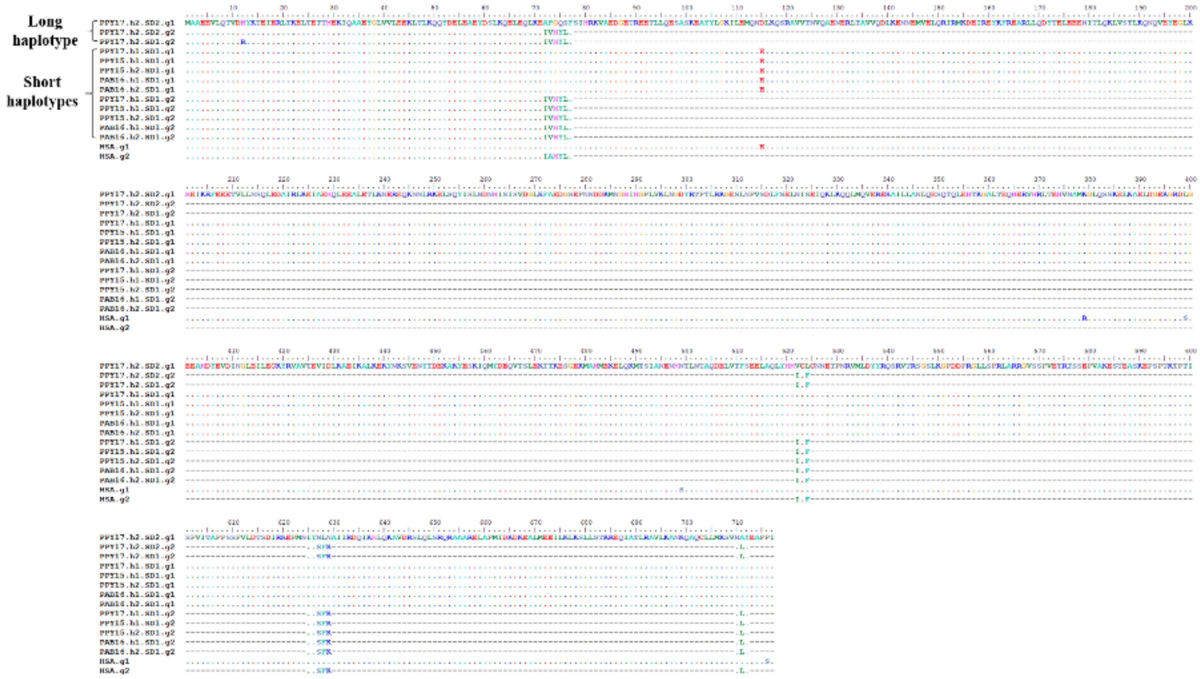


**Fig. S17. Alignment of protein sequences encoding for the two *BICD1* gene models expressed in orangutan and human.**

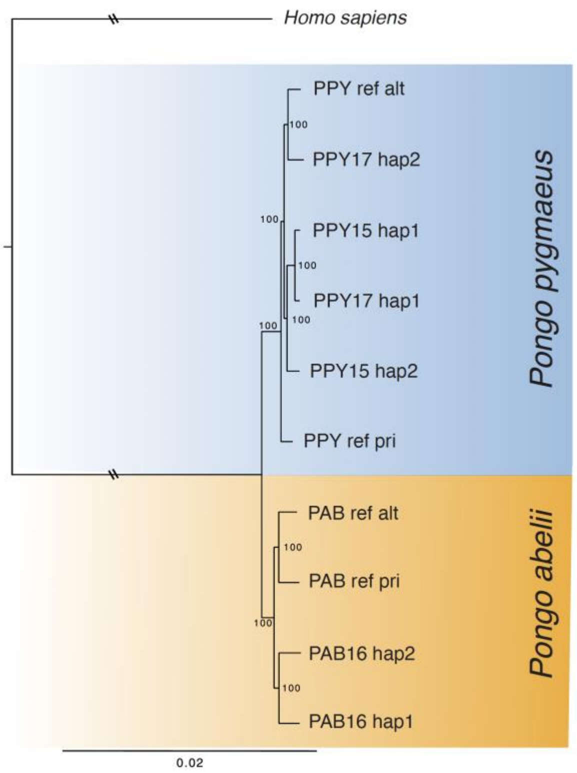


**Fig. S18. Maximum-likelihood phylogenetic tree.** Tree inferred from the entire chromosome 10 haplotypes, reflecting the established divergence between Bornean (PPY) and Sumatran (PAB) species. Homo sapiens used as the outgroup.

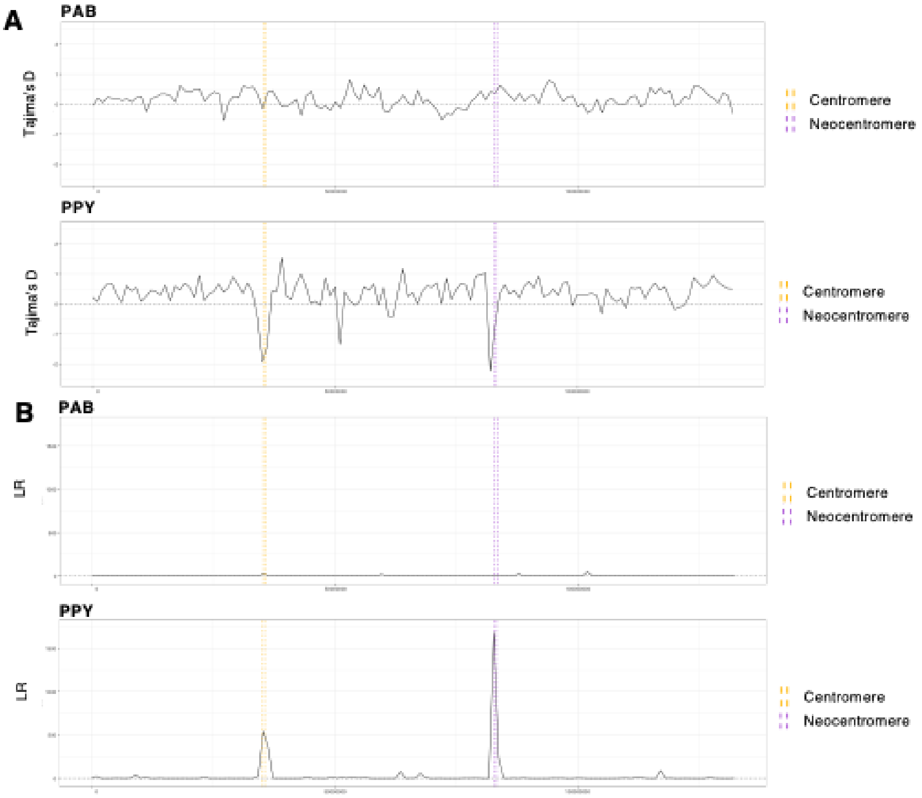


**Fig. S19. Selection scan of chromosome 10 *P. abelii* (PAB) and in *P. pygmaeus* (PPY) separately. (A)** Tajima's D shows an excess or rare variants (D<<0) in proximity of the centromeric regions in PPY, indicating a selective sweep. On the other hand, in PAB no sweep can be identified showing no sign of selection along chromosome 10. **(B)** Similarly, SweepFinder analysis highlights two peaks of Likelihood Ratio (LR) in PPY located in correspondence of the centromeric regions, while PAB shows low levels of LR throughout the entire chromosome. This again suggests strong selective sweeps in those regions occurring in PPY, with possibly a greater selective pressure in the neocentromere because of the higher peak in LR compared to the centromere. Values for the two species were reported with the same scale for a better comparison.

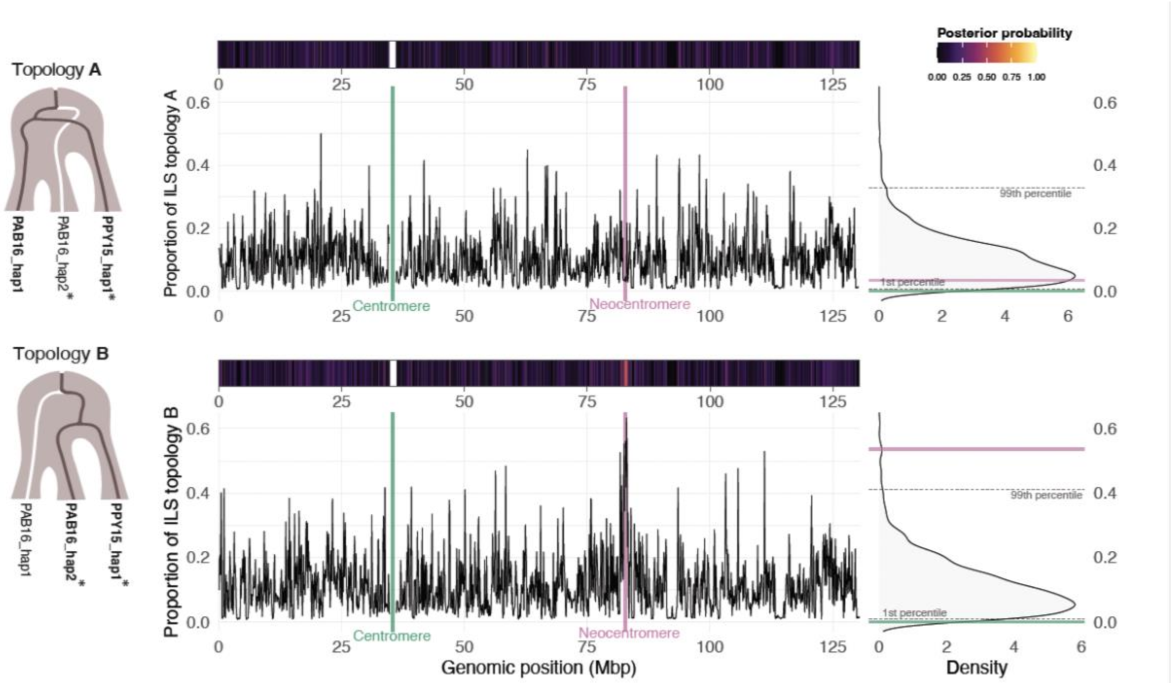


**Fig. S20. Incomplete lineage sorting (ILS) topologies along orangutan chromosome 10 (hsa12) in PAB16_hap1, PAB16_hap2*, and PPY15_hap1* phylogenetic tree.** For each topology, the posterior probability (heatmap, top) and the ILS proportion along chromosome 10 (line plot, bottom) are shown. Windows (100 kbp) encompassing the CEN and NEO highlighted in green and pink, respectively. On the right, ILS proportion distribution is reported.

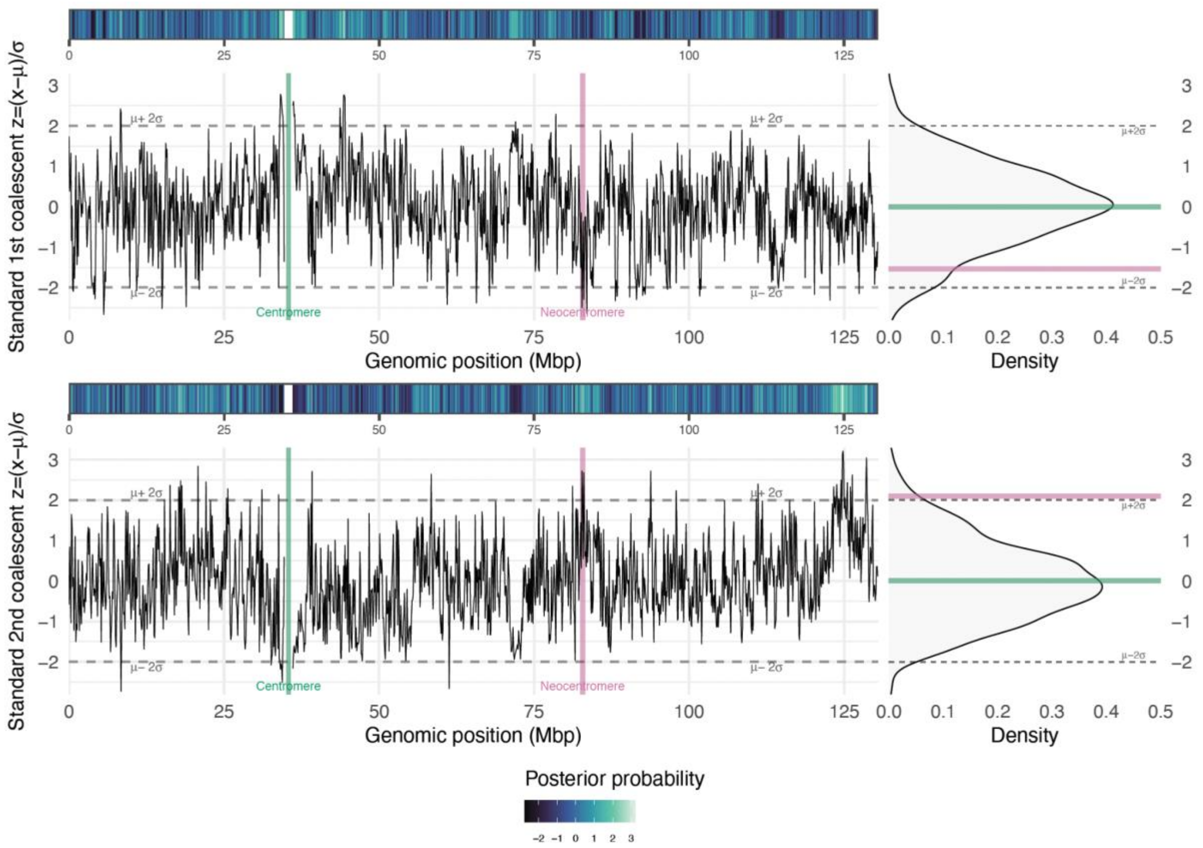


**Fig. S21. Characterization of $1^{st}$ and $2^{nd}$ coalescent patterns along orangutan chromosome 10 (hsa12) for the PAB16_hap1, PAB16_hap2*, and PPY15_hap1* phylogenetic tree.** Coalescent patterns are analyzed using 100 kbp windows. Standardized values for each window are represented as peaks (bottom) or as a color gradient (top), similar to the ILS analysis. Dotted lines indicate two standard deviations from the mean. On the right, the distribution of the first and second coalescent events is shown, with windows encompassing the CEN and NEO annotated in green and pink, respectively.

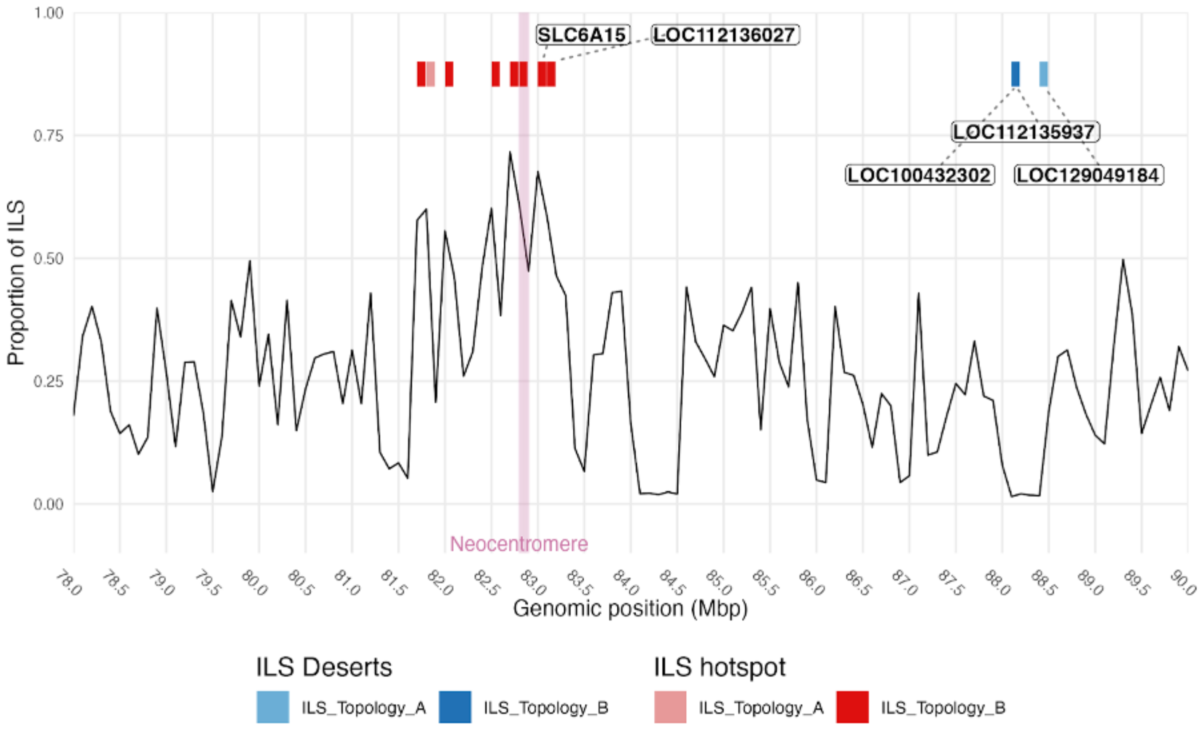


**Fig. S22. ILS proportion in the neocentromere region.** A region spanning 1 Mbp encompassing the neocentromere is shown, with hotspot and desert indicated as for Fig. 4B.

References

61. G. Chiatante, G. Giannuzzi, F. M. Calabrese, E. E. Eichler, M. Ventura, Centromere Destiny in Dicentric Chromosomes: New Insights from the Evolution of Human Chromosome 2 Ancestral Centromeric Region. *Mol Biol Evol* **34**, 1669-1681 (2017).
62. P. S. MOORHEAD, P. C. NOWELL, W. J. MELLMAN, D. M. BATTIPS, D. A. HUNGERFORD, Chromosome preparations of leukocytes cultured from human peripheral blood. *Exp Cell Res* **20**, 613-616 (1960).
63. H. N. Seuanez, H. J. Evans, D. E. Martin, J. Fletcher, An inversion of chromosome 2 that distinguishes between Bornean and Sumatran orangutans. *Cytogenet Cell Genet* **23**, 137-140 (1979).
64. K. Osoegawa *et al.*, A bacterial artificial chromosome library for sequencing the complete human genome. *Genome Res* **11**, 483-496 (2001).
65. A. Daponte *et al.* (Submitted into Current Protocols, 2026).
66. J. Prado-Martinez *et al.*, Great ape genetic diversity and population history. *Nature* **499**, 471-475 (2013).
67. S. Purcell *et al.*, PLINK: a tool set for whole-genome association and population-based linkage analyses. *Am J Hum Genet* **81**, 559-575 (2007).
68. R. A. M. Villanueva, Z. J. Chen, ggplot2: Elegant Graphics for Data Analysis (2nd ed.). *Measurement: Interdisciplinary Research and Perspectives* **17**, 160-167 (2019).
69. A. Rhie, B. P. Walenz, S. Koren, A. M. Phillippy, Merqury: reference-free quality, completeness, and phasing assessment for genome assemblies. *Genome Biol* **21**, 245 (2020).
70. J. T. Robinson *et al.*, Integrative genomics viewer. *Nat Biotechnol* **29**, 24-26 (2011).
71. H. Li, Minimap2: pairwise alignment for nucleotide sequences. *Bioinformatics* **34**, 3094-3100 (2018).
72. H. Li *et al.*, The Sequence Alignment/Map format and SAMtools. *Bioinformatics* **25**, 2078-2079 (2009).
73. N. Huang, H. Li, compleasm: a faster and more accurate reimplementation of BUSCO. *Bioinformatics* **39**, (2023).
74. C. Jain, A. Rhie, N. F. Hansen, S. Koren, A. M. Phillippy, Long-read mapping to repetitive reference sequences using Winnowmap2. *Nat Methods* **19**, 705-710 (2022).
75. M. R. Vollger *et al.*, Long-read sequence and assembly of segmental duplications. *Nat Methods* **16**, 88-94 (2019).
76. K. Katoh, D. M. Standley, MAFFT multiple sequence alignment software version 7: improvements in performance and usability. *Mol Biol Evol* **30**, 772-780 (2013).
77. P. Rice, I. Longden, A. Bleasby, EMBOSS: the European Molecular Biology Open Software Suite. *Trends Genet* **16**, 276-277 (2000).
78. K. Tamura, G. Stecher, S. Kumar, MEGA11: Molecular Evolutionary Genetics Analysis Version 11. *Mol Biol Evol* **38**, 3022-3027 (2021).
79. S. F. Altschul, W. Gish, W. Miller, E. W. Myers, D. J. Lipman, Basic local alignment search tool. *J Mol Biol* **215**, 403-410 (1990).
80. W. C. Earnshaw, H. Ratrie, G. Stetten, Visualization of centromere proteins CENP-B and CENP-C on a stable dicentric chromosome in cytological spreads. *Chromosoma* **98**, 1-12 (1989).
81. M. Pham, Y. Tu, X. Lv, Accelerating BWA-MEM Read Mapping on GPUs. *ICS* **2023**, 155-166 (2023).

82. Y. Zhou, A. W. Leung, S. S. Ahmed, T. W. Lam, R. Luo, Duet: SNP-assisted structural variant calling and phasing using Oxford nanopore sequencing. *BMC Bioinformatics* **23**, 465 (2022).
83. Z. Zheng *et al.*, Symphonizing pileup and full-alignment for deep learning-based long-read variant calling. *Nat Comput Sci* **2**, 797-803 (2022).
84. M. Martin, P. Ebert, T. Marschall, Read-Based Phasing and Analysis of Phased Variants with WhatsHap. *Methods Mol Biol* **2590**, 127-138 (2023).
85. P. Danecek *et al.*, Twelve years of SAMtools and BCFtools. *Gigascience* **10**, (2021).
86. N. Abdennur *et al.*, Pairtools: From sequencing data to chromosome contacts. *PLoS Comput Biol* **20**, e1012164 (2024).
87. N. Abdennur *et al.*, Cooltools: Enabling high-resolution Hi-C analysis in Python. *PLoS Comput Biol* **20**, e1012067 (2024).
88. E. Lieberman-Aiden *et al.*, Comprehensive mapping of long-range interactions reveals folding principles of the human genome. *Science* **326**, 289-293 (2009).
89. G. Benson, Tandem repeats finder: a program to analyze DNA sequences. *Nucleic Acids Res* **27**, 573-580 (1999).
90. M. Tarailo-Graovac, N. Chen, Using RepeatMasker to identify repetitive elements in genomic sequences. *Curr Protoc Bioinformatics* **Chapter 4**, 4.10.11-14.10.14 (2009).
91. A. Morgulis, E. M. Gertz, A. A. Schäffer, R. Agarwala, WindowMasker: window-based masker for sequenced genomes. *Bioinformatics* **22**, 134-141 (2006).
92. I. Numanagic *et al.*, Fast characterization of segmental duplications in genome assemblies. *Bioinformatics* **34**, i706-i714 (2018).
93. B. Q. Minh *et al.*, IQ-TREE 2: New Models and Efficient Methods for Phylogenetic Inference in the Genomic Era. *Mol Biol Evol* **37**, 1530-1534 (2020).
94. S. Zhang *et al.*, Integrated analysis of the complete sequence of a macaque genome. *Nature* **640**, 714-721 (2025).
95. S. Kumar *et al.*, TimeTree 5: An Expanded Resource for Species Divergence Times. *Mol Biol Evol* **39**, (2022).
96. S. Nurk *et al.*, The complete sequence of a human genome. *Science* **376**, 44-53 (2022).
97. D. Porubsky *et al.*, SVbyEye: a visual tool to characterize structural variation among whole-genome assemblies. *Bioinformatics* **41**, (2025).
98. A. Shumate, S. L. Salzberg, Liftoff: accurate mapping of gene annotations. *Bioinformatics* **37**, 1639-1643 (2021).
99. M. Pertea *et al.*, StringTie enables improved reconstruction of a transcriptome from RNA-seq reads. *Nat Biotechnol* **33**, 290-295 (2015).
100. T. Matanis *et al.*, Bicaudal-D regulates COPI-independent Golgi-ER transport by recruiting the dynein-dynactin motor complex. *Nat Cell Biol* **4**, 986-992 (2002).
101. S. V. Indran, M. E. Ballestas, W. J. Britt, Bicaudal D1-dependent trafficking of human cytomegalovirus tegument protein pp150 in virus-infected cells. *J Virol* **84**, 3162-3177 (2010).
102. M. Schubert, S. Lindgreen, L. Orlando, AdapterRemoval v2: rapid adapter trimming, identification, and read merging. *BMC Res Notes* **9**, 88 (2016).
103. H. Li, R. Durbin, Fast and accurate short read alignment with Burrows-Wheeler transform. *Bioinformatics* **25**, 1754-1760 (2009).
104. A. McKenna *et al.*, The Genome Analysis Toolkit: a MapReduce framework for analyzing next-generation DNA sequencing data. *Genome Res* **20**, 1297-1303 (2010).
105. J. Armstrong *et al.*, Progressive Cactus is a multiple-genome aligner for the thousand-genome era. *Nature* **587**, 246-251 (2020).

106. G. Hickey, B. Paten, D. Earl, D. Zerbino, D. Haussler, HAL: a hierarchical format for storing and analyzing multiple genome alignments. *Bioinformatics* **29**, 1341-1342 (2013).
107. A. J. Page *et al.*, SNP-sites: rapid efficient extraction of SNPs from multi-FASTA alignments. *Microb Genom* **2**, e000056 (2016).
108. S. Lifschitz *et al.*, Bio-Strings: A Relational Database Data-Type for Dealing with Large Biosequences. *BioTech (Basel)* **11**, (2022).
109. F. Tajima, Statistical method for testing the neutral mutation hypothesis by DNA polymorphism. *Genetics* **123**, 585-595 (1989).
110. P. Danecek *et al.*, The variant call format and VCFtools. *Bioinformatics* **27**, 2156-2158 (2011).
111. M. DeGiorgio, C. D. Huber, M. J. Hubisz, I. Hellmann, R. Nielsen, SweepFinder2: increased sensitivity, robustness and flexibility. *Bioinformatics* **32**, 1895-1897 (2016).
112. R. Nielsen *et al.*, Genomic scans for selective sweeps using SNP data. *Genome Res* **15**, 1566-1575 (2005).
113. J. Y. Dutheil, S. Gaillard, E. H. Stukenbrock, MafFilter: a highly flexible and extensible multiple genome alignment files processor. *BMC Genomics* **15**, 53 (2014).
114. I. Rivas-González, M. H. Schierup, J. Wakeley, A. Hobolth, TRAILS: Tree reconstruction of ancestry using incomplete lineage sorting. *PLoS Genet* **20**, e1010836 (2024).
115. J. Ishigohoka *et al.*, Distinct patterns of genetic variation at low-recombining genomic regions represent haplotype structure. *Evolution* **78**, 1916-1935 (2024).
116. M. Nordborg, in *Handbook of Statistical Genetics*. (2003).
117. T. Zhu, T. Flouri, Z. Yang, A simulation study to examine the impact of recombination on phylogenomic inferences under the multispecies coalescent model. *Mol Ecol* **31**, 2814-2829 (2022).
118. P. F. Arndt, T. Hwa, D. A. Petrov, Substantial regional variation in substitution rates in the human genome: importance of GC content, gene density, and telomere-specific effects. *J Mol Evol* **60**, 748-763 (2005).
